# Persistent gene flow suggests an absence of reproductive isolation in an African antelope speciation model

**DOI:** 10.1101/2022.12.08.519574

**Authors:** Xi Wang, Casper-Emil Tingskov Pedersen, Georgios Athanasiadis, Genis Garcia-Erill, Kristian Hanghøj, Laura D. Bertola, Malthe Sebro Rasmussen, Mikkel Schubert, Xiaodong Liu, Zilong Li, Long Lin, Emil Jørsboe, Casia Nursyifa, Shanlin Liu, Vincent Muwanika, Charles Masembe, Lei Chen, Wen Wang, Ida Moltke, Hans R. Siegismund, Anders Albrechtsen, Rasmus Heller

**Author notes:** Contributed equally (Xi Wang, Casper-Emil Tingskov Pedersen, Anders Albrechtsen, Rasmus Heller).

## Abstract

African antelope diversity is a globally unique vestige of a much richer world-wide Pleistocene megafauna. Despite this, the evolutionary processes leading to the prolific radiation of African antelopes are not well understood. Here, we sequenced 145 whole genomes from both subspecies of the waterbuck, an African antelope believed to be in the process of speciation. We investigated genetic structure and population divergence and found evidence of a mid-Pleistocene separation on either side of the eastern Great Rift Valley, consistent with vicariance caused by a rain shadow along the so-called ‘Kingdon’s Line’. However, we also found pervasive evidence of not only isolated and recent, but also widespread historical gene flow across the Rift Valley barrier. By inferring the genome-wide landscape of variation among subspecies, we found 14 genomic regions of elevated differentiation, including a locus that may be related to each subspecies’ distinctive coat pigmentation pattern. We investigated these regions as candidate speciation islands.

However, we observed no significant reduction in gene flow in these regions, nor any indications of selection against hybrids. Altogether, these results suggest a pattern whereby climatically driven vicariance is the most important process driving the African antelope radiation, and suggest that reproductive isolation may not set in until very late in the divergence process.

## Introduction

The African continent harbors the world’s most biodiverse community of large mammals^1,2^. Antelopes belonging to the family Bovidae are especially well represented, and a large number of lineages have evolved over the last 20 million years and adapted to a range of different habitats and ecosystems^3^. African antelopes are therefore one of the most fascinating examples of an adaptive radiation among extant large mammals, and the only such radiation that has remained relatively intact since the Pleistocene era, despite many of its member species being currently under severe threat from human-induced factors^4^. Notwithstanding this globally important biodiversity, which is furthermore of huge economic importance in many countries, the evolutionary drivers of the antelope radiation are not well understood.

The waterbuck (*Kobus ellipsiprymnus*) has been considered a model for incipient speciation in African antelopes^5,6,7^. Usually, two subspecies are recognized^8,9,10^, the common (*K. e. ellipsiprymnus*) and defassa (*K. e. defassa*) waterbuck, while some authors suggest these should be elevated to full species level^7^. The two subspecies have different karyotypes^6^, and they are easily distinguishable due to strikingly different patterns of rump coloration: defassa has a completely white rump, whereas common waterbuck has a white elliptical ring around the rump. They are presently parapatrically distributed with defassa occurring to the west of and common waterbuck to the east of the eastern Rift Valley (Fig. 1). The two subspecies are believed to have a contact zone in and around the Rift Valley and its southern extensions in Zambia, and hybrid populations and individuals have been described based on intermediate phenotypes in several localities along this contact zone^5,8^. A genetic study based on microsatellites and mtDNA confirmed that hybridization takes place in the Kenyan populations in Nairobi and possibly Samburu, but also concluded that admixture was a recent phenomenon and established that the subspecies are genetically highly divergent^10^. Accordingly, the current understanding is that hybridization is the result of recent secondary contact between two previously allopatrically diverging evolutionary lineages, and that it only takes place in a few isolated locations along a narrow hybridization zone.

**Fig. 1.**
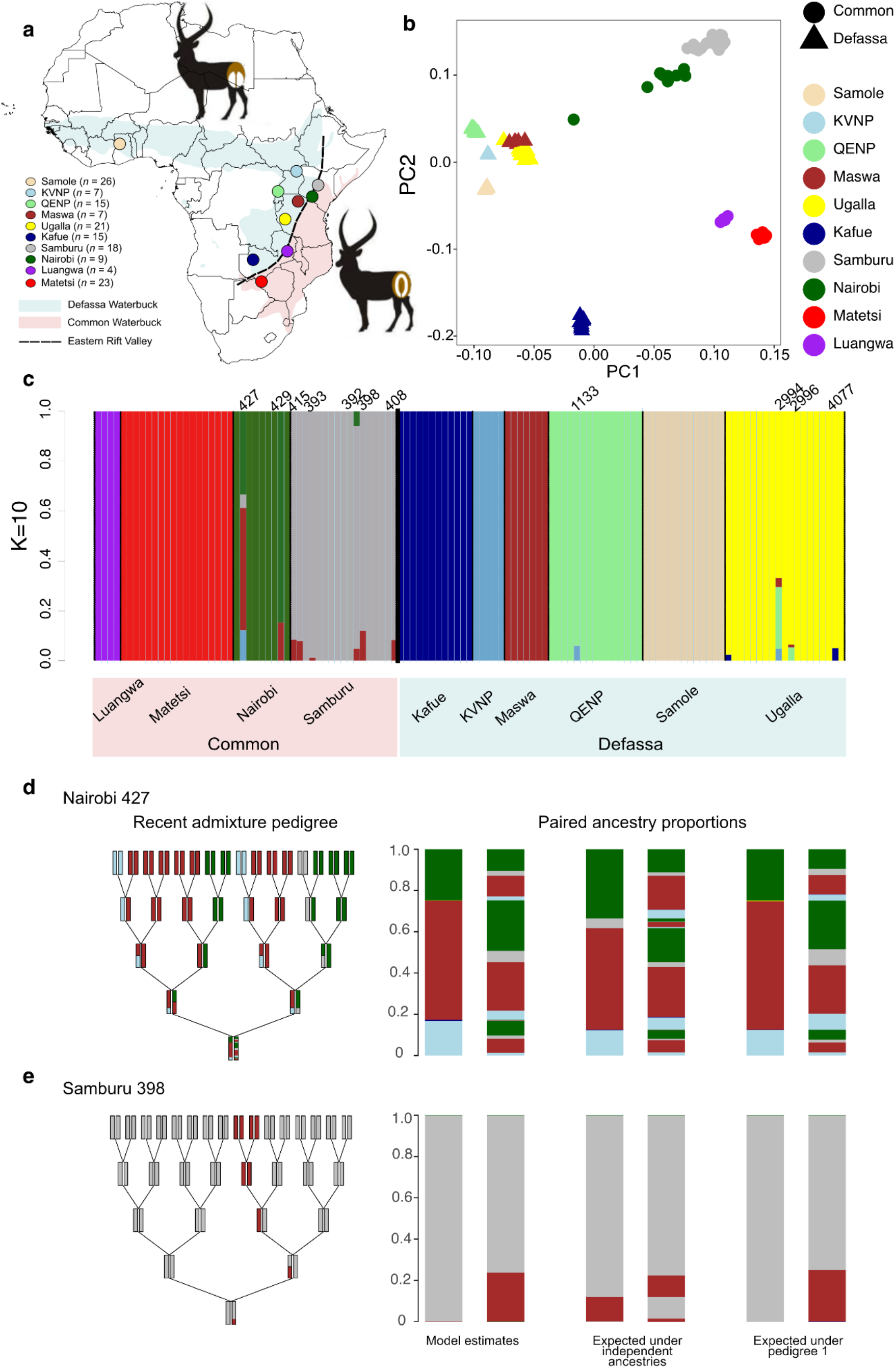
Sampling locations, population structure and recently admixed samples. **a**, Location of origin of the samples analyzed in this study. The dashed line represents the eastern branch of the Great Rift Valley. **b**, PCA plot of waterbuck samples, colored by sample country, showing the first two principal components inferred with PCAngsd. **c**, Admixture proportions of waterbuck samples, estimated with NGSadmix assuming ten ancestral clusters. Admixed individuals are indicated with their sample number. **d**, the observed pattern (left) and the predicted independence pattern (right) of sample 427 from Nairobi and **e**, sample 398 from Samburu using a recently developed method (Garcia-Erill et al. in prep) to show signs of very recent admixture within four generations.

Speciation is a tenet of biodiversity formation, particularly during adaptive radiation when the rate of speciation is accelerated. The speciation process itself has attracted renewed scientific attention since the availability of whole genome data, especially after studies identified a highly heterogeneous genomic landscape of differentiation in diverging populations^11,12,13,14,15,16,17,18^. These findings have led to the realization that speciation can progress in the presence of gene flow, as long as some loci in the genome -sometimes referred to as “speciation islands” -are resistant to gene flow, i.e. there must be selection against admixture in those loci^19,20^. More generally, these and other studies have addressed fundamental questions regarding speciation, including the role of selection in initiating and completing the speciation process^21,22^, i.e. whether or not reproductive isolation tends to occur as an accidental outcome of allopatric population divergence or is itself subject to natural selection^23^. However, as a heterogeneous landscape of genome-wide differentiation can arise for many different reasons, some of which are not related to reproductive isolation^20^, regions of increased divergence do not prove the existence of reproductive isolation. It is therefore necessary to supplement such observations with other analyses that directly address whether there is selection against individuals carrying mixed ancestry in such loci before they can be claimed to play an important role in speciation.

Several features make the waterbuck ideally suited for investigating the genomic aspect of the speciation process: its distribution on either side of an obvious geographical feature, the presence of two evolutionary lineages that are morphologically distinguishable, the known presence of hybrids and the relatively specialized habitat requirement of the waterbuck, which causes its distribution to be tightly linked to standing water^8^. Motivated by this, we used whole genome data to investigate genetic structure, population divergence and gene flow among ten different waterbuck populations across the species range, including two populations in the subspecies contact zone. We first focused on inferring patterns of gene flow, both recent and historical. Next, we performed genome-wide scans to infer the genomic landscape of variation and differentiation to assess whether there are regions of the genome that are resistant to gene flow, and to address whether there are indications of selection against subspecies hybridization. Together, these analyses provide a detailed account of the genomics of speciation in one of the most important mammal radiations on Earth.

## Results

### Sample and sites filtering

Whole-genome resequencing data was generated for 145 waterbuck individuals (Dataset A, Supplementary Table 1) spanning most of the natural distribution and representing 10 locations (Fig. 1a). The mean sequencing coverage was ∼3.4x per individual (Supplementary Table 2). We also sequenced six samples from closely related species: mountain reedbuck, southern reedbuck, Senegal kob, white-eared kob, Uganda kob and puku, and downloaded whole genome resequencing data from a Bohor reedbuck (91.2x) and a red lechwe (94.0x)^24^. Raw reads were mapped to both an internal scaffold-level defassa waterbuck draft genome^24^, and the chromosome-level goat reference genome. After performing rigorous quality inspection of all samples, we removed a total of 26 samples due to either low mapping rate (2 samples), extreme error rates and/or species mislabeling (8 samples), extreme heterozygosity (2 samples), sample duplication (8 samples), relatedness (5 samples) or uncertainty about the sampling locality (1 sample). Hence, the final data set consisted of 119 waterbuck samples (Dataset C) and 8 outgroup samples. We also filtered the reference genome sites and sequencing data, excluding data from regions that showed e.g. poor mapping or other problematic patterns, resulting in 1,119,867,604 (38.7%) remaining sites in the waterbuck reference genome, and 918,938,303 (31.4%) remaining sites in the goat reference genome. Details about the sample and sites filtering process can be seen in Supplementary Materials Section 1 and Section 2, and in Supplementary Figure 1 and Supplementary Table 1, Supplementary Table 2 and Supplementary Table 3.

### Population structure and recent admixture events

We first visualized the population structure of waterbuck by performing a principal component analysis (PCA) of the 119 individuals (Dataset C) using PCAngsd. The PCA shows high congruence between sampling localities and genetic structure within waterbuck (Fig. 1b; Supplementary Figure 3). The first principal component (PC1) captures an overall separation of the two subspecies, whereas PC2 conforms with a North-South gradient.

Notably, two individuals from Nairobi appear intermediate between Nairobi and the defassa populations, potentially reflecting recent admixture between subspecies of waterbuck. In addition, all Kafue samples are positioned somewhat intermediate on PC1, suggesting they could have ancestry from both subspecies.

Admixture proportions based on NGSadmix further supported recent admixture events between waterbuck (Fig. 1c; Supplementary Figure 4). Based on evalAdmix, the model with ten ancestral source populations (K = 10) provides a good fit to the data without excessive correlation of residuals between populations (Supplementary Figure 5), although some correlation of residuals within the inferred populations are still evident, suggesting some remaining substructure or distant relatedness among samples. Overall, individuals clustered almost exclusively to one of the ten assumed ancestral groups, indicating that each sampling locality can be regarded as a distinct population. Eleven individuals from Nairobi, Samburu, Ugalla and QENP, however, showed signs of recent admixture. Among those eleven samples, seven individuals from Nairobi and Samburu had admixture proportions from both subspecies, suggesting that recent admixture is at least as common between subspecies as between populations within subspecies.

Out of the eleven potentially admixed samples identified by NGSadmix, two samples (398 from Samburu and 427 from Nairobi) have paired ancestry proportions that are consistent with recent admixture within the last four generations (Fig. 1d; Fig. 1e; Supplementary Table 4). In the case of sample 427 from Nairobi, it shows a complex recent admixture history involving at least four populations (Fig. 1d), but with a relatively high inconsistency index (Supplementary Table 4). This indicates that the inferred admixture pedigree is not a perfect fit to the paired ancestry. A likely explanation is that the individual is recently admixed with a source that is not well represented in the dataset. The recent admixture history of sample 398 from Samburu is best explained as a single admixture from a Maswa ancestor three generations ago (Fig. 1e). In this case, the inferred admixture pedigree fits nearly perfectly to the paired ancestry, and the inconsistency index is low (Supplementary Table 4), which gives strong support for the sample being a double backcross. These two samples involve admixture between defassa and common waterbuck populations, confirming ongoing gene flow between the subspecies. For the remaining recently admixed individuals we found no evidence of the admixture event being within the last four generations. However, this does not rule out that they are the result of admixture.

For example, they could derive from admixture slightly older than four generations ago, they could involve admixture from unsampled (ghost) populations or they could be the result of recent admixture where both parents have similar admixture proportions.

To further visualize the evolutionary relation between the populations, we excluded the eleven recently admixed samples (Dataset D) and built a neighbor-joining (NJ) tree including the eight outgroups. The NJ tree supported the PCA and admixture results, showing a basal split between the two waterbuck subspecies and a clear North-South evolutionary division within each subspecies (Supplementary Figure 6). Individuals from each sampling locality cluster together, consistent with results from the PCA and admixture analyses (Fig. 1b; Fig. 1c; Supplementary Figure 3; Supplementary Figure 4). We therefore concluded that sampling locations are a reasonable proxy for genetic populations and retained them as the unit of genetic analyses. In subsequent analyses where we are mostly interested in evolutionary and demographic events on a longer time scale, we excluded the eleven recently admixed samples unless otherwise noted (Dataset D).

### Genetic diversity and population differentiation

We inferred genome-wide heterozygosity using realSFS for the 108 un-admixed individuals (Supplementary Figure 7). Overall, the two subspecies harbor similar levels of genetic diversity, with an average heterozygosity of 0.00359 per bp in the common waterbuck and 0.00355 per bp in the defassa waterbuck. The defassa populations KVNP and QENP had the lowest mean heterozygosity values (0.00312 and 0.00296, respectively), whereas the two Zambian populations Luangwa and Kafue had the highest values (0.00536 and 0.00426, respectively). The levels of genetic diversity from Luangwa and Kafue are high compared to other large African mammals we have investigated, e.g. 0.00396 in Cape buffalo (Quinn et al. in prep), 0.00207 in desert warthog and 0.00262 in common warthog^25^, and 0.00201 in African leopard^26^, and also higher than many other species of large mammal^27^. Additionally, heterozygosity distributions were almost entirely non-overlapping when considering adjoining populations, providing further support to consider each sampling location as genetically distinct populations.

Using realSFS we also quantified the genetic differentiation between populations and found that F_st_ was generally high between the two subspecies (average F_st_ between subspecies 0.27; highest F_st_ 0.39 between Matetsi and QENP), while F_st_ was lower within subspecies (average F_st_ within common 0.11 and within defassa 0.13; lowest F_st_ 0.06 between Matetsi and Luangwa) (Supplementary Figure 8), in agreement with the PCA and NJ tree (Fig. 1b; Supplementary Figure 3; Supplementary Figure 6). Kafue in central Zambia had almost the same F_st_ value with all other populations, including overlapping values when compared with other defassa populations (F_st_ 0.15-0.22) and common populations (F_st_ 0.17-0.27). The ambiguous pattern of differentiation observed in Kafue coincides with Kafue’s intermediate position on PC1 and with Kafue having the highest genetic diversity of any defassa population, both of which support that this population could have an admixed origin.

The spatial distribution of genetic variation was further investigated using EEMS^28^. The main feature is a gene flow barrier overlapping the eastern branch of the Great Rift Valley. However, there are additional barriers emerging from the EEMS analysis, suggesting that the Rift Valley is not the only important barrier to gene flow, and the subspecies divergence is not the only important genetic divide in the waterbuck (Fig. 2c). Interestingly, the overall solid gene flow barrier overlapping with the Rift Valley appears more permeable in two places, including the region surrounding Nairobi (Fig. 2c) where hybrids have been identified.

**Fig. 2.**
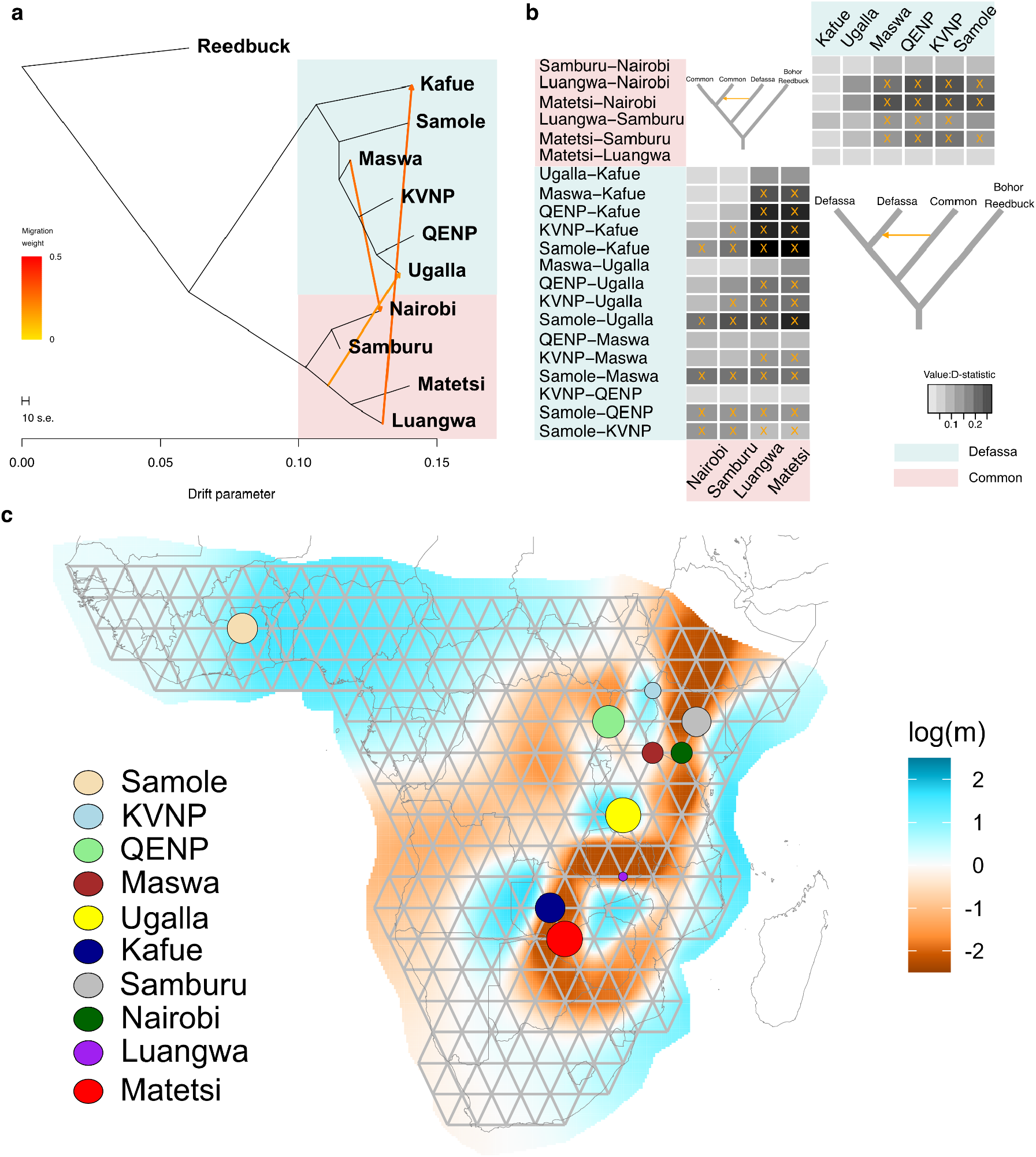
Historical admixture between defassa waterbuck and common waterbuck. **a**, Population tree inferred with TreeMix assuming three migration events. Three reedbuck samples (Mountain reedbuck, Southern reedbuck and Bohor reedbuck) were used as outgroups. Arrows represent migration edges, with colour indicating the migration weight (proportion of the admixed population estimated to derive from the source population). **b**, D-statistics when using waterbuck population as P1, P2 and P3, and Bohor reedbuck as outgroup P4, where the x-axis representing P3 and y-axis representing P1-P2. Star represents the significant median P value for each combination. D-statistics was estimated by single read sampling for all possible combinations of samples using ANGSD. **c**, EEMS analysis showing the location of barriers to gene flow between sampling localities. The color scale depicts the inferred value of *m*, the effective rate of gene flow, with blue representing areas of relatively high gene flow and red representing areas of relatively low gene flow.

The divergence time between the waterbuck subspecies has not been estimated before using genetic data. Using Fastsimcoal2 we estimated it to be 374 kya (95% CI:365-386; Supplementary Figure 9; Supplementary Table 5). A model without gene flow can be regarded as a lower bound on the inferred divergence time, as adding gene flow would push back the divergence time.

### Historical admixture between waterbuck subspecies

To investigate the historical process of population divergence and infer patterns of historical admixture between populations of waterbuck, we performed a TreeMix analysis on 108 un-admixed individuals, assuming from zero to three migration edges (Fig. 2a). All trees showed a deep split between a defassa and a common lineage, and a further North-South basal divergence within both waterbuck lineages, in line with results from PCA, NJ tree and the pairwise F_st_ values. All of the inferred migration edges connected populations across the subspecies: between Nairobi and Maswa in the northern part of the range, and between Kafue and Luangwa, and between Ugalla and the ancestor of Matetsi-Luangwa in the southern part of the range, indicating geographically widespread gene flow between the two subspecies of waterbuck across Kingdon’s Line.

To corroborate these gene flow observations we used D-statistics (ABBA-BABA) to test all possible population trees consistent with the overall subspecies topology, using the 108 unadmixed samples to exclude confounding from recent admixture (Fig. 2b; Supplementary Figure 10; Supplementary Figure 11). We observed strong signals of ancient gene flow between subspecies. Out of 60 combinations of defassa-defassa-common tests, 34 had median D-stat values that were significant, suggesting gene flow between populations from different subspecies. The highest D-stat values were found in combinations including Kafue, a northern defassa population (Samole, QENP, KVNP or Maswa) and a southern common population (Luangwa or Matetsi). Similarly, 15 out of 36 common-common-defassa combinations had significant median D-stat values, and here the highest were found in combinations with Nairobi, one of the southern common populations (Luangwa or Matetsi) and one of the northern defassa populations (Samole, QENP, KVNP or Maswa). These results corroborate the TreeMix results by suggesting extensive past gene flow between the two subspecies, which likely occurred both in the northern and in the southern part of the species range.

### Investigating genomic islands of divergence

After having found pervasive evidence for gene flow between the two subspecies, we performed a range of sliding window genome scans to detect ‘islands of divergence’ between the two subspecies, which may in turn be candidates for ‘speciation islands’. First, we identified 14 regions of highly elevated F_st_ between the pooled subspecies samples, defined as being in the top 0.01% of all F_st_ windows and combining adjacent windows (distance < 1Mbp) into one single region (Fig. 3a; Fig. 3b).

**Fig. 3.**
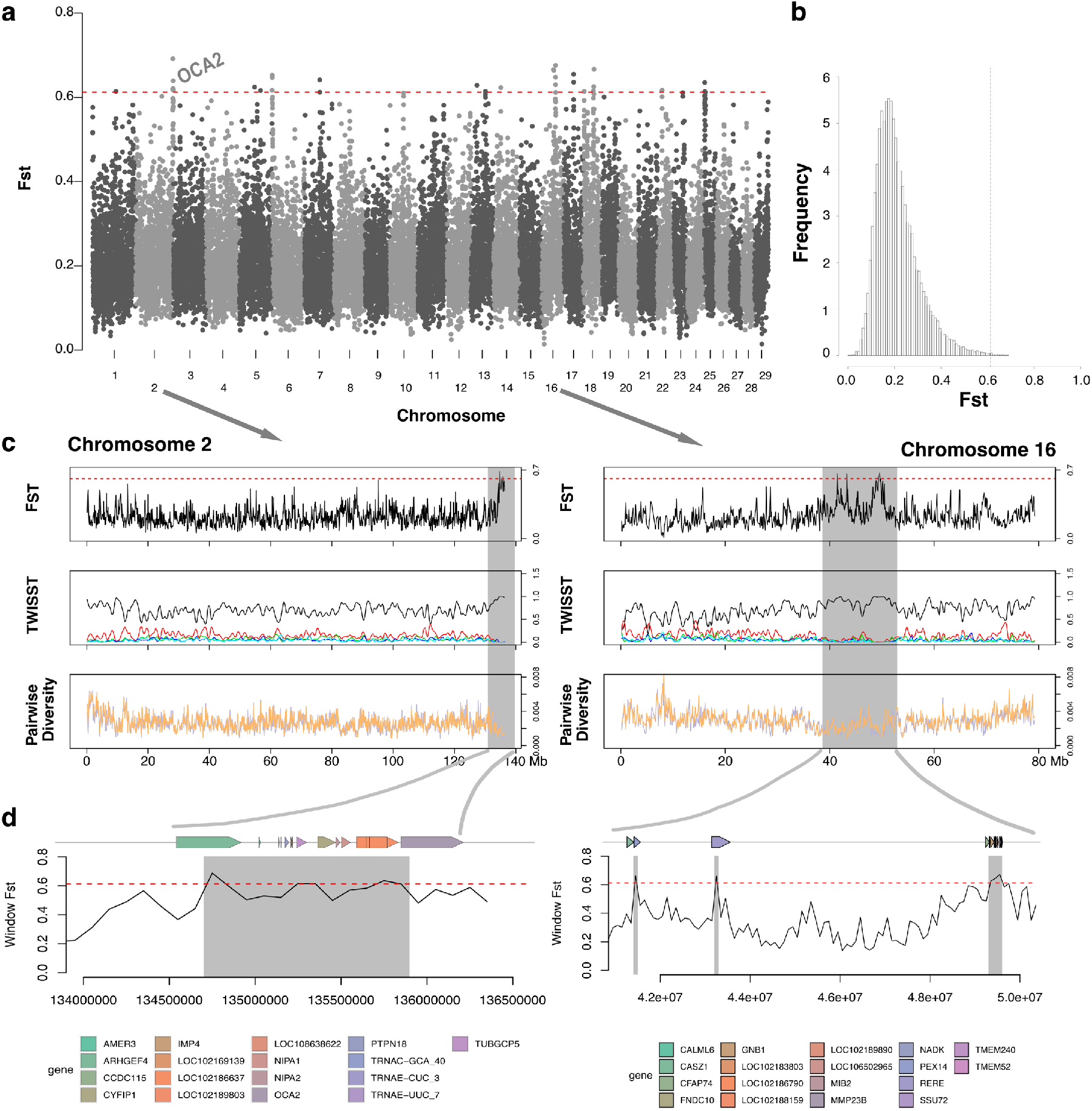
Genome-wide scans to detect genomic islands of divergence between defassa waterbuck and common waterbuck. **a**, Manhattan plot of F_st_ values estimated for 100 kb windows, estimated from genotype likelihood between the pooled two subspecies samples. **b**, Distribution of the F_st_ values plotted in a, indicating the 99.9% quantile (red dashed line) used as a threshold to detect outliers. **c**, Plots of genome wide F_st_ and variation in local topological relationships between the two subspecies using TWISST, and plots of pairwise diversity for defassa waterbuck (orange) and common waterbuck (light purple). Supplementary Figure 13 shows plots for all 11 chromosomes in the genome scan. **d**, Plot of F_st_ within four outlier F_st_ windows distributed in chromosome 2 and 16, and its surrounding genomic region, together with its annotated protein coding genes. Supplementary Figure 14 shows plots for all 14 F_st_ windows within 11 chromosomes.

Second, we investigated the variation in local topological relationships between the two subspecies using TWISST. This analysis supported that the 14 islands of divergence are enriched for population topologies that have defassa and common waterbuck as monophyletic units (Fig. 3c; Supplementary Figure 12; Supplementary Figure 13; Supplementary Table 6).

To evaluate the role of natural selection in the divergence between subspecies of waterbuck, we estimated pairwise diversity, Tajima’s D, and the pattern of localized linkage disequilibrium across the whole genome(Fig. 3c; Supplementary Figure 13; Supplementary Table 6). We observed reduced pairwise diversity, reduced negative Tajimas’ D, and increased linkage disequilibrium across all 14 divergence islands, suggesting that natural selection, possibly interacting with the recombination landscape, is likely to have played a role in elevating divergence between two subspecies of waterbuck in these particular regions of the genome. Interestingly, the region on chromosome 2 contains the genes *HERC2* and *OCA2*, which we hypothesize may be related to the distinctive coat pigmentation patterns in each subspecies of waterbuck (Fig. 3a; Fig. 3d; Supplementary Figure 14).

### No sign of reduced gene flow in divergence islands

To qualify as genomic islands of speciation, regions should show a significant reduction of gene flow and selection against hybrid ancestry^19,20^. We investigated gene flow in the divergence islands by performing a series of analyses focusing on the 14 regions identified above (Fig. 4). In all regions we found that individuals grouped into two distinctive clusters, each representing one of the two subspecies, and in agreement with the analyses used to define these as divergence islands. However, in all 14 regions we also observed a small number of individuals that were almost exactly intermediate in local ancestry as estimated using PCA (Fig. 4a; Supplementary Figure 15). Furthermore, the intermediate individuals consistently showed elevated heterozygosity in these regions compared to the two clusters representing each subspecies (Fig. 4b; Supplementary Figure 15), indicating that these individuals are heterozygous for two highly divergent and subspecies-associated haplotypes. In addition, in some divergence islands we found a small number of individuals that grouped entirely with the “wrong” subspecies (ten individuals in total distributed across nine islands), indicating that they are homozygous for a haplotype that is almost exclusively found in the other subspecies (Fig. 4c; Supplementary Figure 15). Collectively, we will refer to these individuals as the “disparate” individuals in each divergence island. The disparate individuals were not identical across the 14 islands, but there was a clear tendency of populations close to the contact zone being overrepresented among them. For example, the ten individuals that grouped entirely with the other subspecies in one of these highly discriminative regions came from Ugalla (2993, 2998, 4078, 5618), Maswa (8880, 5624), Nairobi (421, 428, 434) and Samburu (401), and only the geographically remote Samole population has no disparate individuals. As expected, the recently admixed individuals identified using NGSadmix were overrepresented among disparate individuals, but there were several other disparate individuals that showed no sign of recent admixture according to NGSadmix. We further assessed whether the disparate individuals are likely to be the result of semi-recent admixture (admixture too old to be detected by NGSadmix) by checking whether they have unusual D-statistic values relative to other members of their population, but this was not the case (Supplementary Figure 16). Given the prominent role of sex chromosomes during population divergence and speciation^29^, we also separately investigated the X chromosome using only the female samples. The X chromosome has higher overall F_st_ and more complete lineage sorting than the autosomes (Supplementary Figure 17), but we found only two additional candidate divergence islands, which showed similar patterns as the autosomal divergence islands (Supplementary Figure 18). Based on all these observations, we draw five conclusions: i) these patterns can only be explained by gene flow even in these highly differentiated regions of the genome, ii) a non-negligible proportion of this gene flow is not very recent, iii) there is no indication of selection against hybrids in these regions, iv) the reduced recombination rate inside these regions means that migrant haplotypes remain unbroken, making migrant individuals easy to detect, and v) gene flow must have been considerable, given the detected number of individuals with migrant ancestry in these regions, and the finding even of individuals carrying two migrant haplotypes. In summary, there is no support for these divergence islands also being candidate speciation islands.

**Fig. 4.**
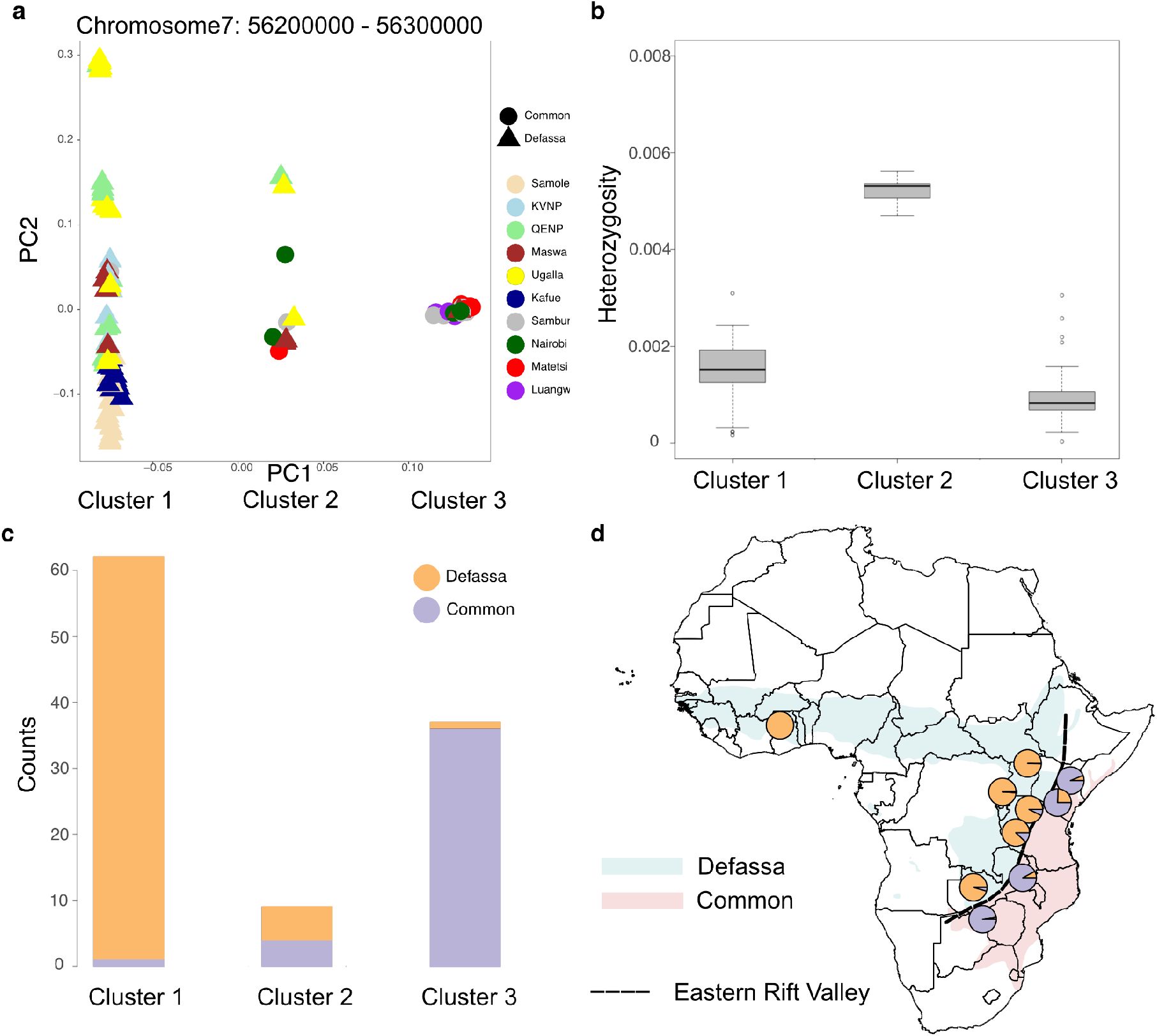
Patterns of gene flow in divergence islands. A representative island on chromosome 7 is depicted in panels a, b and c, and results across all 14 autosomal islands are summarized in panel d. **a**, Population structure within a divergence island on chromosome 7 inferred by PCAngsd, showing highly discrete structure. **b**, Heterozygosity estimation in the divergence island on chromosome 7 and grouping individuals into three clusters according to PC1 in a. Heterozygosity is clearly elevated in Cluster 2. **c**,The number of individuals from each subspecies that belong in each of the three clusters in the divergence island on chromosome 7. Supplementary Figure 15 shows the corresponding plots for all 14 divergence islands. **d**, The average proportion of membership in “defassa typic” (orange) or “common typic” (light purple) PCA clusters across all 14 autosomal divergence islands. Individuals in the intermediate cluster (Cluster 2) are counted as having half membership in each cluster.

## Discussion

We confirm here that the waterbuck separates into two distinct genetic lineages which correspond to the two subspecies, common and defassa waterbuck. We estimate that these two lineages separated at least 374 kya (95% CI: 365-386), much earlier than the early Holocene date suggested by^8^. However, we also show that there is substantial further structure within each of the two subspecies, with high genetic differentiation between northern and southern populations within each subspecies. These findings present the first highly-resolved picture of genetic structure in waterbuck, and show that both the Samole and Kafue populations are genetically distinct from all other defassa populations with F_st_ ranging from 0.11-0.20 and 0.15-0.22, respectively. We hypothesize that the strong genetic structure even within subspecies could be caused by the narrow habitat preference of waterbucks^8^, inhabiting a widespread, but patchy habitat type. In particular, the strong dependency on standing water would have made the waterbuck sensitive to any climatic changes that affect the distribution of free water. These types of climatic changes were prevalent throughout the volatile climate in the Pleistocene, in which tropical Africa alternated between wet pluvials and dry interpluvials^30,31^. Accordingly, the eastern edge of the Rift Valley is associated with a rain shadow so prominent that it is known in biogeography as ‘Kingdon’s Line’^32,33^. During dry periods of the Pleistocene, very little free surface water would have been available along this rain shadow^34^, which could therefore have been the immediate agent of vicariance between the two waterbuck lineages.

Despite the clear genetic separation between subspecies, we found multiple lines of evidence for gene flow between them. Elaborating on the findings of Lorenzen et al.^10^ we found seven individuals in Samburu and Nairobi that appear recently admixed. Using a new method we infer the most likely admixture pedigrees for these individuals and show that one individual is a result of a double backcross between subspecies. Interestingly, the seven cases of recent between-subspecies admixture outnumber the four within-subspecies population admixture cases, which underlines that ongoing gene flow could be as prevalent between subspecies as within subspecies.

Moreover, after removing potentially recently admixed samples we also found clear evidence of ancient gene flow between subspecies, which contrasts with previous studies concluding that admixture is exclusively a recent phenomenon^10^. Both the TreeMix and D-stat analyses showed strong evidence of excess allele sharing between many combinations of defassa-common waterbuck populations, notably suggesting ancient gene flow both in the southern and in the northern part of the species range. These results are supported by the EEMS analysis, which shows patches of increased permeability in the otherwise solid genetic barrier along the Rift Valley. We also found that Kafue, in particular, is almost equally genetically distant to nearby common as nearby defassa populations. We therefore conclude that there must have been at least periodically substantial amounts of gene flow between defassa and common waterbuck, and that this gene flow is also taking place presently.

Reconciling the findings of extensive gene flow despite clear genetic structure we propose that waterbucks may have followed a mixing-isolation-mixing (MIM) model^35^, whereby populations would have become separated when their preferred habitat got fragmented, and reconnected under more benign conditions. This is in line with the assumed continental refugial biogeography for many African ungulates^31^, and we conclude that the waterbucks may be more prone to climatic fluctuations due to their water dependence than most other species, explaining why their genetic structuring is more pronounced than many other African mammal species^26^.

The presence of extensive and widespread gene flow between populations belonging to different waterbuck subspecies questions whether these can be considered on the path to speciation, as has been suggested several times^5,6,7^. However, as a large body of recent literature shows, speciation is possible even in the face of gene flow, notably if some parts of the genome resists introgression^21,36,37^. We therefore investigated whether there are indications of such speciation islands forming between the waterbuck subspecies. We did identify 14 highly differentiated and highly lineage sorted regions of the genome. Despite this, our investigations showed that variation in these regions is not consistent with speciation islands. For example, we found evidence that there are several individuals of both subspecies that carry haplotypes from the other subspecies in these regions. These are most likely the result of gene flow rather than old standing variation, as we found them more often in populations surrounding the subspecies contact zone. Such introgression of the “divergence islands” appears not to be under any negative selection, neither in heterozygous nor in homozygous state, and we found indications that it persists for a long time and at relatively high frequencies in the recipient populations. Therefore, we conclude that they are very unlikely to qualify even as incipient speciation islands of the genome.

Given that the highly diverged regions of the genome do not confer reproductive isolation, there are several other possible explanations for their occurrence. Most of them show evidence of being under positive selection, but they are also characterized by reduced recombination rates. Regions with reduced recombination rates are more likely to become divergence islands due to the higher efficiency of linked background selection^36,38,39^. In addition, these regions show several characteristic patterns of large-scale inversions, e.g. clear clustering according to PCA and elevated heterozygosity in the intermediate samples^40^. Interestingly, we identified one divergence island corresponding to a location on goat chromosome 2 that contains the genes *HERC2* and *OCA2*, which has been linked with hair and skin color polymorphism in many animals, including humans^41,42,43^, medaka^44^, the Mexican cave tetra^45^, and zebrafish^46^. It would be interesting to investigate whether the conspicuous rump hair color pattern difference that distinguishes common and defassa waterbucks could be related to the strong differentiation in this genomic region. Even so, physical observations of intermediate rump patterns in several overlapping parts of the subspecies range^8,47^ are consistent with our conclusions that such diagnostic differences are not associated with reproductive isolation, and could simply be the result of background selection or even genetic drift, possibly coupled with an inversion and low recombination in the underlying parts of the genome.

Based on our finding of widespread and recurrent gene flow between the two waterbuck subspecies, including the parts of the genome that are most highly differentiated, we conclude that reproductive isolation has not occurred to any substantial degree in this model system for incipient antelope speciation, despite a relatively old divergence time dating to at least 374,000 years ago and despite (polymorphic) differences in karyotypes^6^. Rather, we find that waterbuck population divergence is primarily, or perhaps exclusively, due to vicariance, i.e. allopatric separation driven by barriers that come and go in response to habitat changes, in particular those that influence the distribution of standing water.

Moreover, reduncines separated by up to 3.8 million years of evolutionary divergence^24^ can readily hybridize both in captivity and in nature^8^, suggesting that reproductive isolation may take very long to set in these taxa. These observations have important implications for understanding speciation in the African antelope guild. If population divergence is predominantly vicariant, i.e. driven by barriers that emerge due to climate changes, and not driven by ecogeographic separation^48^, then outbreeding depression during secondary contact caused by local adaptations and divergent selection^49^ may be insignificant. In such a biogeographic scenario, the retention of reproductive compatibility for long periods of evolutionary time may even be a selective advantage, as it would facilitate mixing of gene pools when geographic conditions favor it. These findings are consistent with the growing body of evidence for cross-species gene flow even across extended periods of evolutionary divergence, and hence a more fluid interpretation of the species concept^50^.

Our results also raise some pertinent questions about the management and conservation of genetic variation in waterbuck, some of which may be applicable to other African species. First, we found that the waterbuck cannot be simply regarded as two discrete evolutionary or management units^51^ corresponding to the two subspecies, with Kafue as a notable example of a population that clearly violates this classification. Second, the prevalence and persistence of naturally occurring gene flow between the two subspecies question whether each of them contain important local adaptations that could lead to outbreeding depression in hybrids. Lastly, if genetic admixture between highly divergent lineages, or even hybridization between full species, is a widespread natural process in the complex biogeographical history of the African antelopes, these processes need to be better understood before we can make informed decisions about what the most relevant conservation units are.

## Methods

### Sample collection

A total of 145 samples (Dataset A) were collected between 1992 and 1998 from 10 localities within the waterbuck distribution range (Fig. 1a; Supplementary Table 1). Two of them, Nairobi and Samburu, are known to harbor individuals of intermediate phenotypes^10,52,53^.

We also included a single sample from each of six closely related species within the antelope tribe Reduncini: the mountain reedbuck (*Redunca fulvorufula*) and southern reedbuck (*R. arundinum*); the Senegal (*Kobus kob kob*), white-eared (*K. k. leucotis*) and Uganda kob (*K. k. thomasi*) and the puku (*Kobus vardoni*). Additionally, we downloaded publicly available whole genome sequencing data from a Bohor reedbuck (*R. redunca*) and a red lechwe (*Kobus leche leche*) generated as part of the Ruminant Genome Project^24^.

### Lab protocol

Samples consisting of skin tissue, biopsies or earnicks were kept in a DMSO buffer in the field, stored at -20 °C as soon as possible, and were further transferred to a -80 °C freezer for long-term storage. The QIAGEN DNeasy Blood & Tissue Kit (QIAGEN, Valencia, CA, USA) was used for DNA extraction, following the manufacturer’s protocol. RNase A was subsequently added to get RNA-free genomic DNA. Before using gel electrophoresis to check the quality of the genomic DNA, we further measured the DNA concentrations with a Qubit 2.0 Fluorometer and a Nanodrop.

### Sequencing

All samples were sequenced to approximately average 3.4X depth of coverage using Illumina paired-end 150 bp reads on Illumina’s HiSeq 2000 platform. The two downloaded samples (Bohor reedbuck and red lechwe) were sequenced to much higher depths of 91.2X to 94.0X. We assessed the quality of the raw reads using FastQC (http://www.bioinformatics.babraham.ac.uk/projects/fastqc/) and MultiQC^54^ before mapping.

### Mapping

For mapping, we used a modified version of PALEOMIX ‘bam’ pipeline^55^ (https://github.com/xiqtcacf/WaterbuckScripts), which is a pipeline designed for the processing of demultiplexed high-throughput short-read sequencing data.

We first trimmed Illumina universal adapters using AdapterRemoval v2.3.2^56^. No trimming of low-quality bases and ambiguous bases (Ns) was performed. We merged overlapping paired-end reads into a single sequence with default parameters in order to improve the fidelity of the overlapping region by selecting the highest quality base when mismatches are observed. Mismatching positions in the alignment where both bases had the same quality were set to ‘N’ via the --collapse-conservatively option. Finally, empty reads resulting from trimming primer-dimers were removed.

We then used BWA mem v0.7.17-r1188^57^ to map the reads to: 1) a scaffold-level defassa waterbuck draft genome^24^, and 2) the chromosome-level reference genome from a male San Clemente goat (ARS1; GenBank: GCA_001704415.1). PCR duplicates were flagged using samtools v1.11 ‘markdup’ for paired reads and PALEOMIX ‘rmdup_collapsed’ for merged reads.

The resulting BAM files were merged and filtered based on standard BAM flags to exclude unmapped reads, reads with unmapped mate reads, secondary alignments, reads that failed QC, PCR duplicates, and supplementary alignments. In addition, we excluded reads with filtered mate reads, and reads in alignments with inferred insert sizes smaller than 50 bp or greater than 1000 bp, reads where fewer than 50 bp or less than 50% of the read were aligned, and read pairs in which mates mapped to different contigs or not in the expected orientation. Statistics of the filtered BAM files were generated by samtools ‘stats’ and ‘idxstats’^58^.

### Sample filtering

To remove problematic samples prior to downstream analyses, we performed quality inspection on all samples by considering various mapping statistics: GC-content in each sample after mapping, heterozygosity, error rate, and between-sample relatedness. The pipeline for sample filtering is described in Supplementary Materials Section 1 and can be found in the github page: https://github.com/xiqtcacf/WaterbuckScripts.

### Sites filtering

To avoid biases from low-quality mapping^26^, we further performed a series of quality controls on the reference sequences and on the data sets mapped against the internal reference (defassa waterbuck) and the external reference (goat). The site filtering involves exclusion of problematic regions of the reference genomes (based on mappability, repeats and sex-linked chromosomes or scaffolds), and exclusion of sites that showed unusual depth or excess heterozygosity after mapping. For the sites filtering steps we used only the samples (Dataset B) retained after the sample filtering steps outlined above. The pipeline for sites filtering is described in Supplementary Materials Section 2.

### Genotype likelihood calculation

We estimated genotype likelihoods by using the GATK model (-GL 2) in ANGSD, inferring the major and minor allele from the genotype likelihoods (-doMajorMinor 1), estimating the allele frequencies (-doMaf 1), resulting in beagle likelihood files for both data sets, i.e. mapped against both references. Filtering was performed simultaneously, including only sites that passed the sites filtering, a minimum p-value of 1e-6 to call a SNP under the likelihood ratio test (-SNP_pval 1e-6), a minimum allele frequency filter of 0.05 (-minMaf), a minimum mapping quality of 25 (-minMapQ), and a minimum base quality score of 30 (-minQ).

### Population structure and recent population admixture

#### NGSrelate within each location

Before downstream analyses, we performed a second analysis of relatedness within each sampling location by using NgsRelate, which can be used to infer relatedness for pairs of individuals from low coverage Next Generation Sequencing (NGS) data by using genotype likelihoods instead of called genotypes^59,60^. We first generated a file with allele frequencies (-doMaf 1) and a file with genotype likelihoods (-doGlf 3) for all samples (Dataset B) passing sample filtering and sites filtering criteria using ANGSD, with additional filtering of -hwe_pval 1e-6 -minmaf 0.05 -minMapQ 30 -minQ 20 -SNP_pval 1e-6. The frequency column from the allele frequency file was then extracted and used together with the genotype likelihood file by NgsRelate. We identified six additional pairs of relatives within the sampling localities Samole (K1>0.15 and K2>0.8), Matetsi (K1>0.8), and Ugalla (K1>0.39 and K2>0.15), and removed the lower depth sample from each pair (removing 7600, 7602 in Samole; 1597, 1607 in Matetsi; and 5620 in Ugalla, Supplementary Figure 1).

#### Principal component analysis using PCAngsd

We performed principal component analysis (PCA) using PCAngsd, which is based on an iterative heuristic approach of estimating individual allele frequencies to infer population structure^61^. Low depth sequencing data sets are associated with statistical uncertainty, and a model-based approach in PCAngsd can account for this uncertainty by working directly on genotype likelihoods of the unobserved genotypes, improving accuracy in samples with low sequencing depth^61^. We used ten eigenvectors to assess population structure, equivalent to the number of study localities (Dataset C).

#### Admixture proportion analysis using NGSadmix and evalAdmix

We estimated admixture proportions for 119 remaining waterbuck individuals (Dataset C) using NGSadmix^62^ based on genotype likelihoods. We ran NGSadmix with multiple seeds from K=2 to K=10 until convergence, which we defined as a maximum difference of two log-likelihood units between the top three maximum likelihood results. For the converged run of each K value, to identify the best-fitting value of K, we subsequently evaluated the model fit using evalAdmix^63^, which estimates the pairwise correlation of the residuals matrix between individuals. A good fit of the data to the admixture model results in correlation close to 0. A positive correlation represents individuals with similar demographic histories that is not reflected in the admixture proportions, and a negative correlation represents individuals with different histories despite not being modeled as such in the admixture proportions. Both positive and negative correlations indicate a poor model fit.

#### Evaluating cases of recent admixture

We used a recently developed method (Garcia-Erill et al. in prep) to infer the admixture history of eleven individuals showing signs of recent admixture. This method is based on inferring genome-wide patterns of ancestry heterozygosity, which will differ from the expectation under statistical independence only in cases of very recent admixture (1-5 generations old). The method tests if the estimated patterns are more compatible with a recent admixture pedigree or with a pedigree where admixture is relatively older (above 5 generations). The method furthermore provides indices of how well the data fits the model assumptions, which can be used to evaluate alternative explanations for the observed admixture result (Garcia-Erill et al. in prep).

#### NJ-tree

To construct a Neighbor joining (NJ) tree we estimated an identity-by-state (IBS) matrix as a measure of the pairwise genetic distance between individuals. We did this by randomly sampling a single read at each site for each sample (-doIBS 1) and calculating the pairwise IBS matrix between individuals (-doCounts 1 -makeMatrix 1) by ANGSD. The NJ tree (Dataset D) excluding all individuals identified as recently admixed individuals by NGSadmix was inferred using the R package ape^64^.

### Ancient admixture

#### Treemix

As a first investigation of the history of population splits and ancient admixture in the waterbuck, we performed a TreeMix analysis^65^ on 108 waterbuck samples within ten populations (excluding 11 samples identified as recently admixed samples, Dataset D), and included three reedbuck samples (mountain reedbuck, southern reedbuck and Bohor reedbuck) as outgroups. We first estimated the likelihood of sample allele frequency (SAF files) for each population with ANGSD, using only sites that passed all site filters, polarized using the external reference goat genome as the ancestral state. All populations were then merged by keeping only sites with no missing data across all populations, resulting in 8.3 million sites as input data. The input allele counts per population were finally generated by calling within each population the maximum likelihood sample frequency. We ran 25 differently seeded replicates for each possible migration from 0 to 3 using TreeMix and chose the run with the highest likelihood, breaking ties at random. We forced the reedbucks to be outgroups and used a block size of approximately 14K variable input sites, corresponding to an assumed maximum LD block size of 5 Mbp.

#### ABBABABA

To further investigate the presence of ancient gene flow between defassa waterbuck and common waterbuck populations, we performed D-statistics (ABBA-BABA) tests^66^,^67^ for data sets from 108 samples (excluding all recently admixed individuals, Dataset D) mapped to the external reference Goat to avoid potential internal reference bias, with the Bohor reedbuck as outgroup. We first applied the function -doAbbaBaba 1 implemented in ANGSD, which samples a random base at each position to estimate the counts of ABBA and BABA sites between each triplet of P1, P2 and P3 individuals, using only sites that passed the above sites filtering and blocks of a predefined size of 5 MB (-blockSize 5000000). Subsequently, a block jackknife approach was used to estimate standard errors and to calculate Z scores^68^. To take into account the large number of comparisons, we performed a Bonferroni correction on the inferred raw p-values^69^.

### Diversity and divergence

#### Heterozygosity

We estimated the genome-wide heterozygosity for 108 waterbuck samples (Dataset D) using realSFS in ANGSD. For this we used the same approach as in the sample filtering (see Supplementary Materials Section 1), but here we included all sites remaining after the site filtering (see Supplementary Section 2). We first estimated the single-individual SFS using realSFS, and then divided the number of heterozygous sites by the total number of sites.

#### Pairwise global F_st_

To estimate genome-wide estimates of F_st_ for each pair of waterbuck populations, we first generated per-population site allele frequency files (saf) from genotype likelihoods using - doSaf 1 in ANGSD using the goat reference allele as ancestral. For each pairwise population, we then inferred population pairwise 2d-SFS from saf files with realSFS^70^. Using these 2dSFS as a prior, genome-wide pairwise F_st_ were estimated by ‘realSFS fst’ in ANGSD, using Hudson’s F_st_ estimator, which is less sensitive to differences in sample size between populations^71^.

#### EEMS

To evaluate the relative effective migration rates, we used EEMS (estimated effective migration surface), which produces a visual representation of population structure that can highlight potential areas of deviation from isolation by distance, which is indicative of gene flow patterns^28^. We used ANGSD to calculate a pairwise IBS matrix for 108 waterbuck samples (Dataset D) by sampling a random base at each position. This matrix was used as input for EEMS, along with the sample coordinates and the habitat coordinates. We used three independent runs of 20 million steps and a burn-in of 10 million steps, with 300 demes and all sites retained after the sites filtering to perform runeems_snps of EEMS for the data sets. The results were visualized using rEEMSplots (https://github.com/dipetkov/eems).

#### Divergence time estimation

The divergence time between defassa waterbuck and common waterbuck was investigated using a coalescent simulation based method implemented in Fastsimcoal2 v2.7^72^. The Samole population was used to represent defassa waterbuck and Matetsi was used to represent common waterbuck, based on results from Treemix and D-statistics, which suggested these populations were least likely to have received gene flow from the other subspecies. We inferred the unfolded 2d-SFS between Samole and Matetsi mapped to the waterbuck reference, unfolding by using the red lechwe to represent the ancestral allele. We assumed a simple model only considering divergence time without gene flow between the two populations. The rationale for this model is that it infers a lower boundary of the true divergence time by not considering gene flow, which is furthermore warranted by choosing populations far away from any potential hybrid zone. For this model we ran 100 independent Fastsimcoal runs to find the best-fitting parameters yielding the highest likelihood, with 500,000 coalescent simulations per likelihood estimation (-n500000), 100 conditional maximization algorithm cycles (-L100), and minimum 100 observed SFS entry count taken into account in likelihood computation (-C100). To obtain the 95% confidence interval of the model, we generated 100 non-parametric bootstraps from a global saf file of the original dataset using ‘--method pseudo-loo’, a jackknife resampling approach based function in winsfs^73^. For each bootstrap, we performed 100 independent Fastsimcoal runs, using the same settings as for the analyses of the original dataset, and calculated the 95% quantile and standard error for parameters from the maximum likelihood run from each bootstrapped SFS. A mutation rate of 1.43e-8 per site per generation^24^ and a generation time of 7.1 years^74^ were used to convert model estimates from coalescence units to absolute values (i.e. years).

### Genome-wide scans

#### Sliding window F_st_

To identify genomic regions showing exceptionally high levels of sequence divergence between the two subspecies of waterbuck, we performed a genome-wide scan by pooling respectively the four common waterbuck populations (Luangwa, Matetsi, Nairobi and Samburu) and the six defassa populations (Samole, Maswa, KVNP, QENP, Ugalla and Kafue) (Dataset D). Per-population site allele frequency files (saf) based on data sets mapped to the goat reference were then generated for both of these pooled groups using - doSaf 1 in ANGSD. Pairwise 2d-SFS from saf files were then estimated using realSFS. Based on the site allele frequencies and 2dSFS priors, we also estimated Hudson’s F_st_ in 100kb windows using ‘realSFS fst’ in ANGSD. We restricted the analysis to the sites retained after the sites filtering outlined above and furthermore removed any windows that contained < 10,000 sites, which resulted in keeping windows with information in at least 31.3% of sites (915 million sites retained out of 2922 million sites). We retained the top 0.1% of the remaining window F_st_ values and merged outlier windows into a single region if the distance between windows was < 1 Mbp. We considered these regions as candidates of ‘divergence islands’ and extracted the genes overlapping with these regions from the goat reference genome annotation (https://ftp.ncbi.nlm.nih.gov/genomes/all/GCF/001/704/415/GCF_001704415.1_ARS1/GCF_001704415.1_ARS1_genomic.gtf).

We also scanned the X chromosome, adjusting the approach to accommodate the ploidy difference compared to autosomes and taking into account the higher expected F_st_ in X chromosomes compared with autosomes^29^. For this scan we used only female individuals based on SATC results (44 individuals, Supplementary Figure 2) and re-ran the above site filters by using only the female samples. We defined the top 0.1% outlier windows based only on results from the two X chromosomal scaffolds in the goat reference genome.

#### TWISST

To examine and quantify the variation in local topological relationships between the two subspecies throughout the whole genome, we used TWISST^75^ for the 108 non-admixed individuals (Dataset D). In order to reduce the number of possible topologies, we combined populations of Nairobi and Samburu into one population of common waterbuck located in northern part (NorthCom), populations of Matetsi and Luangwa in southern part of common waterbuck (SouthCom), populations of Maswa, KVNP, QENP, Ugalla as one located in the central distribution of defassa waterbuck (CentralDef), and treated Kafue, Samole as separate populations of defassa waterbuck (Fig. 1a), hence grouping populations based on both geography and the population split tree from TreeMix (Fig. 2a). We first calculated the IBS matrix per 100 kb window based on single read sampling by -doIBS 1 in ANGSD, and then generated one NJ tree per window using the R package ape^64^. The exact weightings by considering all subtrees were then calculated using the default method ‘complete’ in TWISST, which collapses monophyletic nodes and is fast if the taxa are well resolved^75^.

#### Genome-wide summary statistics

To further investigate the genome-wide pattern of variation on a subspecies level, we estimated nucleotide diversity, Tajima’s D and linkage disequilibrium (LD) estimation in sliding windows of 100 kb. We used the data mapped to the goat reference and pooled all individuals (Dataset D) from each subspecies into one. We used ANGSD to estimate pairwise nucleotide diversity and Tajima’s D by first using ‘dosaf 1’ to calculate the per-population site allele frequency likelihood, and then estimated the global site frequency spectrum using ‘realSFS’ based on the expectation maximization algorithm^70^. The function ‘saf2theta’ was further used to calculate thetas for each site from the posterior probability of allele frequency (global SFS) and ‘thetaStat’ was finally used to output pairwise nucleotide diversity (pi) and Tajima’s D^76^ in 100 kb non-overlapping sliding windows. We estimated LD represented by the correlation coefficient between pairs of loci (r^2^) using the genotype likelihood based program ngsLD^77^, with a minimum maf filter of 5% and randomly sampling 1% of all SNP pairs.

### Detection of gene flow within divergence islands

To investigate gene flow in detail in the divergence islands, we performed a series of analyses in each divergence island using data sets mapped to goat reference for 108 samples (Dataset D) across all ten populations of waterbuck. For each island, we first inferred local population structure by using PCAngsd^61,78^. We then binned the first principal component (PC1) into five bins and calculated the individual heterozygosity for individuals within each bin. To further investigate patterns of gene flow in these divergence islands, we also investigated D-stat results for samples that had unusual inferred ancestry, i.e. ancestry from the other subspecies.

## Supporting information

Waterbuck-SupplementaryFile

## Acknowledgements

We thank Amal al-Chaer for her laboratory work on extracting DNA from the tissue samples. XW, LDB and RH were supported by an European Research Council Starting Grant (No. 853442), and RH was furthermore supported by a Carlsberg Foundation Young Researcher grant (CF21-0497).

## Data availability

The raw sequencing data generated for this project have been deposited in FastQ format in SRA with BioProject accession code XXXXXXX. Scripts used to generate all analyses and plots can be found in the github page https://github.com/xiqtcacf/WaterbuckScripts.

