## Supplementary material for "Persistent gene flow suggests an absence of reproductive isolation in an African antelope speciation model": Waterbuck-SupplementaryFile

* Contributed equally

### Supplementary Materials Section 1: Sample filtering

*Basic mapping statistics filtering*

We first evaluated statistics gathered as part of the BAM filtering step described above, including insert size distributions, mapping rates, average coverage, and the number of reads excluded based on each filtering criteria. The quality of all BAM files were further assessed using FastQC (http://www.bioinformatics.babraham.ac.uk/projects/fastqc/) and MultiQC[54] as post-mapping quality checking. We also evaluated the depth distribution per sample by using ANGSD[79] (-doCounts1 -doDepth1) (https://github.com/ANGSD/angsd,). Sample 1219 and Sample 2589 were removed because of the extremely low mapping rate (<0.1%) (Supplementary Table 2).

*Error rates*

Sample sequencing error rates were measured using the ‘perfect individual’ method[80] implemented in ANGSD, as excess mismatches between each sample and the outgroup (Goat reference), relative to the mismatches between the outgroup and a consensus sequence of a high depth sample, the ‘perfect individual’ (sample 1133). The idea is that each ingroup sample should have the same expected number of derived alleles as the ‘perfect individual’ when using an outgroup, and that a surplus of observed derived alleles in each sample is thus due to excess errors relative to the ‘perfect individual’. A consensus sequence was first created for the ‘perfect individual’ using ‘-doCounts 1 -doFasta 2’ option in ANGSD, and error rates were then estimated using all bases (-doAncError 1). For generating the consensus sequence and for the error estimation, we used quality filters including a minimum base quality of 30 and minimum mapping quality of 20. Based on this, sample 2553 showed extremely high error rate (1.34%), samples 7543, 7544 and 7593 showed relatively high error rates (0.10-0.43%), and four samples showed numerically large negative error rates: 7398, 7399, 7400, 7401 ([-0.03]-[-0.02%]), indicative of being phylogenetically closer to the outgroup than the 'perfect individual'. These individuals were removed from downstream analyses (Supplementary Table 2).

*Heterozygosity*

We calculated site allele frequency likelihood based on individual genotype likelihoods assuming Hardy-Weinberg equilibrium (HWE) for all samples using -doSaf 1 in ANGSD. A per-individual site frequency spectrum (SFS) was estimated by realSFS in ANGSD. Based on individual SFS, the genome-wide heterozygosity for all samples was finally calculated by dividing the estimated number of heterozygous sites by the total number of sites. Two samples (1244 and 7321) showed extremely high values of heterozygosity and were consequently removed from further analyses (Supplementary Table 2).

*Duplicates filtering based on King calculated from global 2D-sfs*

Using the methodology described in Waples et al.[81], we identified closely related and potentially duplicated samples by first inferring the two-dimensional site frequency spectrum (2d-SFS) for each pair of samples. Three statistics were then calculated directly from the 2d-SFS for each pair of samples: R0, R1, and the KING-robust kinship coefficient. These can be used to identify close familial relatives without the need to estimate population allele frequencies. We identified 14 duplicate pairs with KING-robust kinship values > 0.330, and proceeded to exclude all but one sample from set of duplicated pairs (samples 7597, 7595, 7594, 1614_1, 1615_1, 2997, 1612, 397, Supplementary Table 2). For a few analyses (see Results) we chose to instead combine all data from each duplicated individual to achieve higher sequencing depth, which allowed us to validate the findings from the low depth data.

### Supplementary Materials Section 2: Sites filtering

*Reference filtering: mappability, repeats and sex-linked scaffolds*

The mappability of each site of both internal and external reference genomes were estimated by GenMap (v1.2.0)[82], conservatively using 150bp k-mers with up to 2 mismatches (-K 150 -E 2) and defaults for the remaining settings. All sites with a mappability score less than one were excluded. We also discarded low complexity and repeat sequences in both reference fasta files using RepeatMasker v.4.1.1 (http://www.repeatmasker.org/). To identify sex-linked and abnormal scaffolds in the defassa waterbuck reference, we used SATC[83]. This resulted in 437 scaffolds being identified as X chromosome-linked and excluded from further analyses. In addition, we also removed all scaffolds smaller than 100 kb when using defassa waterbuck as reference. For the chromosome level goat reference, we only retained the 29 autosomal chromosomes (Supplementary Table 3).

*Global depth filtering*

We estimated global depth per site across all samples using ANGSD. We then estimated the mean and median depth per site across data sets and an average pooled depth across all individuals, and further excluded all sites that had a global depth below 375 (average depth * total number of samples) or above 1.5 times the median (655.5).

*Excess heterozygosity (HWE) filtering*

Mapping of repetitive DNA sequences to a reference genome creates ambiguous mapping of reads originating from multiple genome locations to a single location, resulting in sites with an excess of heterozygosity. Regions of extreme heterozygosity are indicative of such problematic mapping. We filtered such regions by first estimating genotype likelihoods for all waterbuck samples that passed the above sample quality control, using ANGSD with the GATK genotype likelihood model and basic quality filters (-SNP_pval 1e-6 -minmaf 0.05 -minMapQ 25 -minQ 30). Using these genotype likelihoods as input to PCAngsd[61,78], we then calculated the per-site inbreeding coefficients (F), ranging from -1 where all samples are heterozygous to 1 where all samples are homozygous, and performed a Hardy-Weinberg equilibrium (HWE) likelihood ratio test accounting for population structure. The optimal number of principal components to model the population structure was inferred based on Velicer's minimum average partial test[84] implemented in PCAngsd. Finally, we treated sites with $F < -0.9$ and p-value < 1e-6 as excessively heterozygous and conservatively excluded all sites within 10kb of such sites. We did this for both data sets, mapped to defassa waterbuck and goat reference (Supplementary Table 3).

### Supplementary Materials Figures

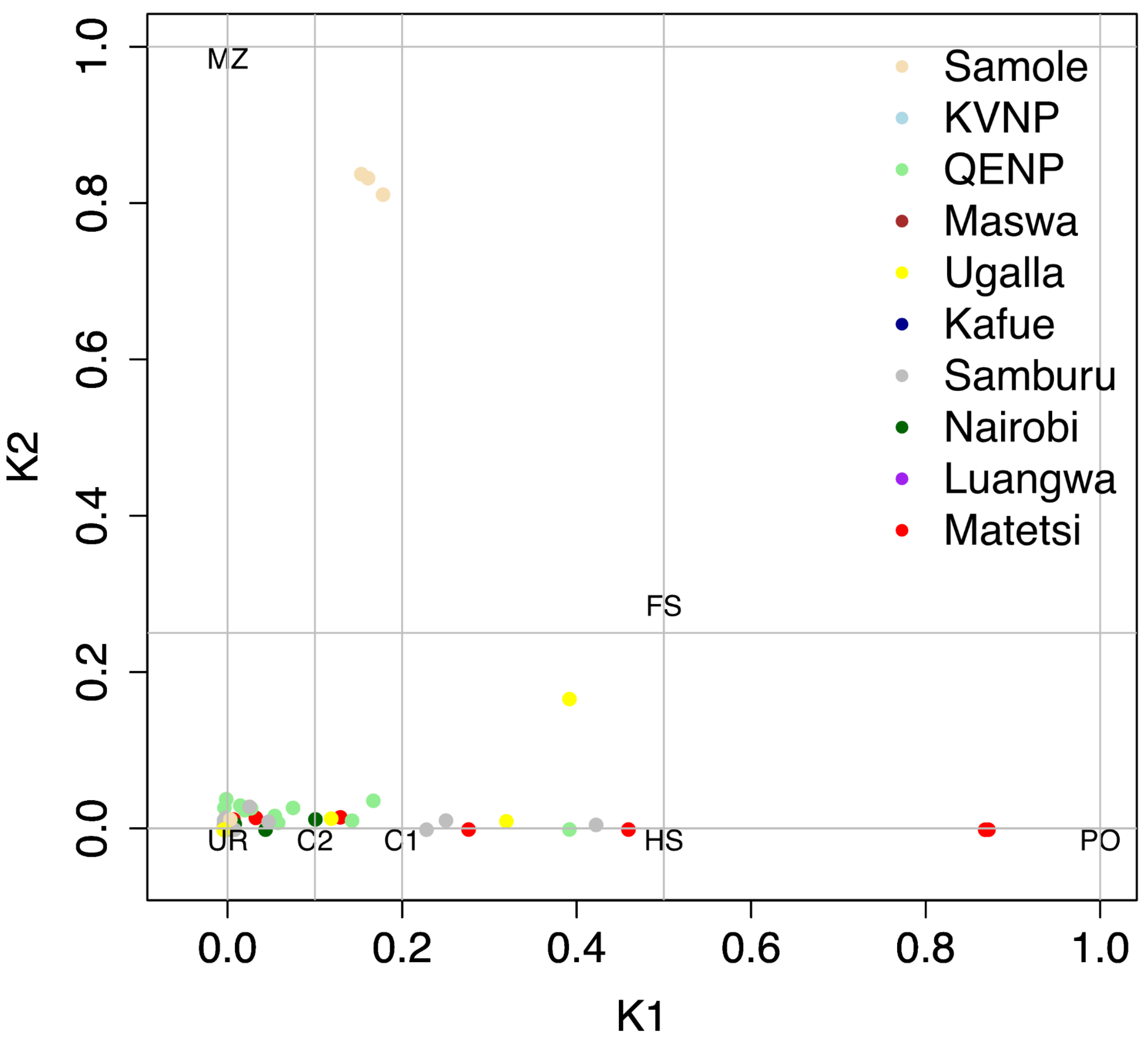

**Supplementary Figure 1**. Estimates of relatedness for waterbucks within each sampling location by NGSrelate. Six pairs of relatives within the sampling localities Samole, Matetsi, and Ugalla were identified, and the lowest coverage sample in each pair was removed (removing 7600 and 7602 in Samole, both of which repeat twice in three pairs; 1597, 1607 in Matetsi; and 5620 in Ugalla).

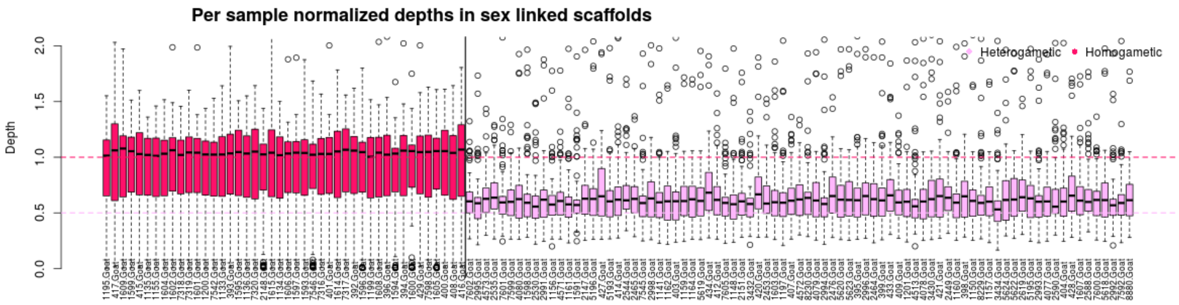

**Supplementary Figure 2**. Boxplot of normalized depth of sex-linked scaffolds for each sample. The samples are ordered based on the two inferred sex groups identified by the Graphical User Interface (GUI) version of SATC. Expected values for each group are shown by horizontal green dashed lines of 0.5 (heterogametic) and 1.0 (homogametic). The vertical line separates the inferred heterogametic and homogametic samples.

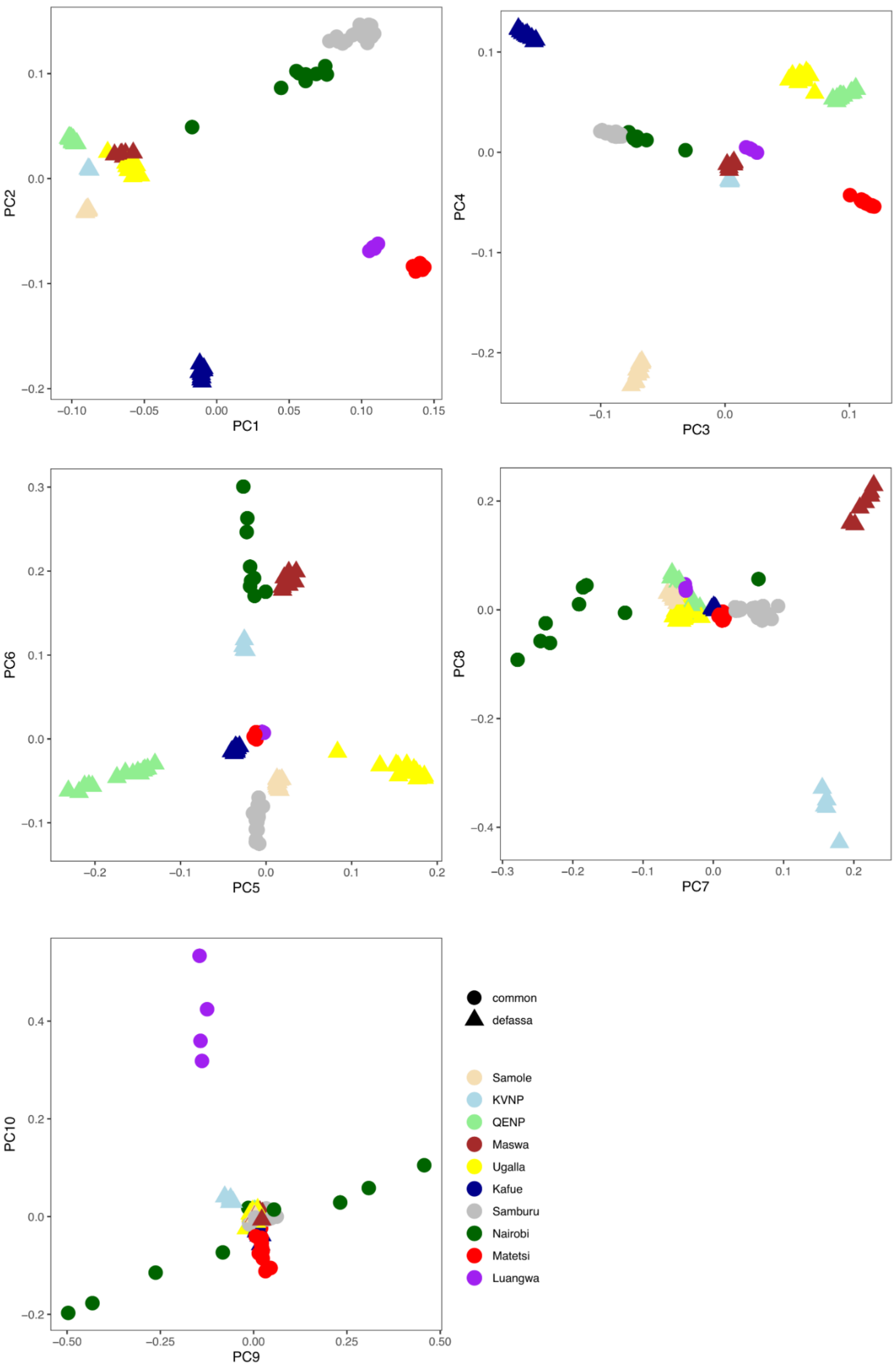
PC1
PC2
PC3
PC4
PC5
PC6
PC7
PC8
PC9
PC10
common
defassa
Samole
KVNP
QENP
Maswa
Ugalla
Kafue
Samburu
Nairobi
Matetsi
Luangwa

**Supplementary Figure 3**. PCA plot of waterbuck samples colored by sampling locality on the first ten principal components inferred with PCAngsd.

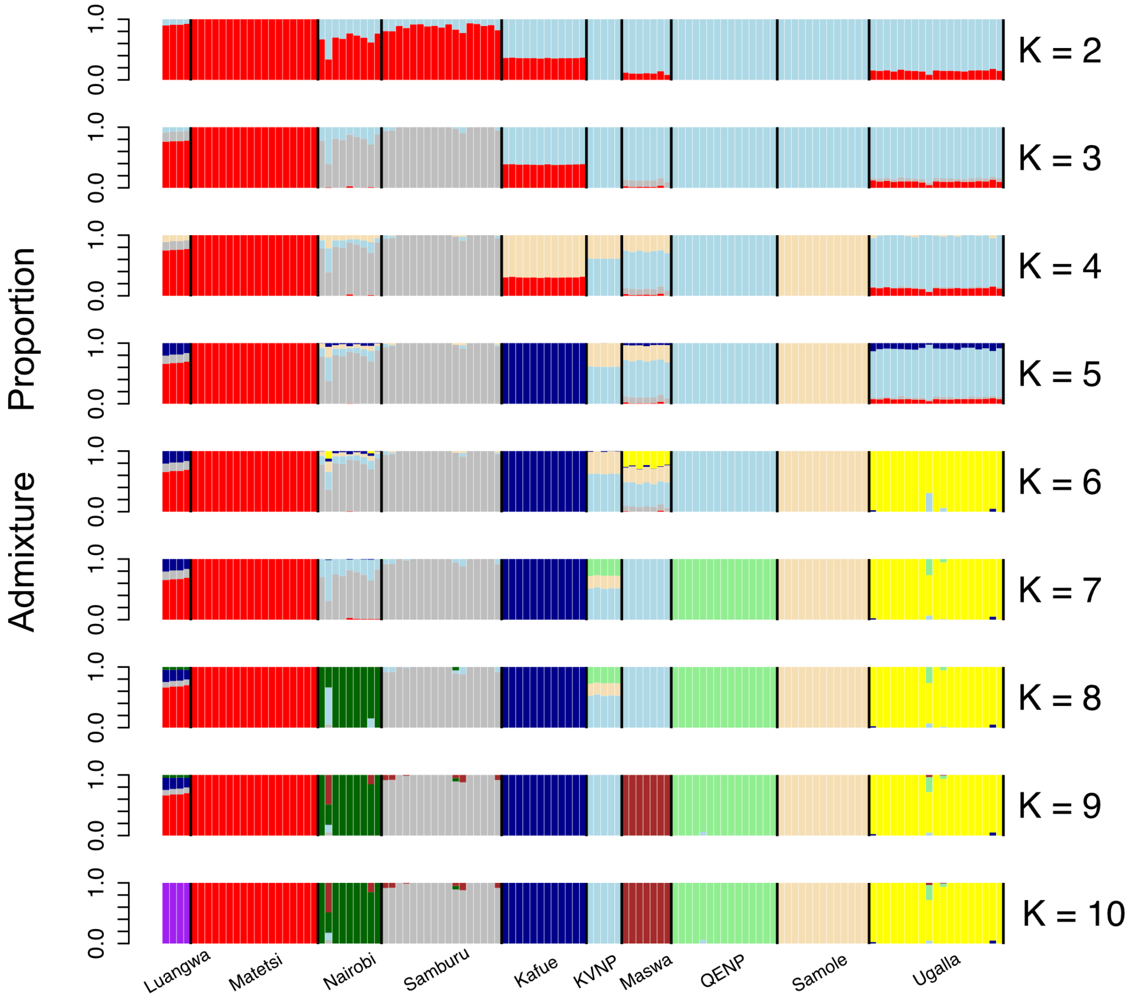

**Supplementary Figure 4**. Individual admixture proportions estimated with NGSadmix assuming from K = 2 to K = 10. Individuals are grouped by sampling locality.

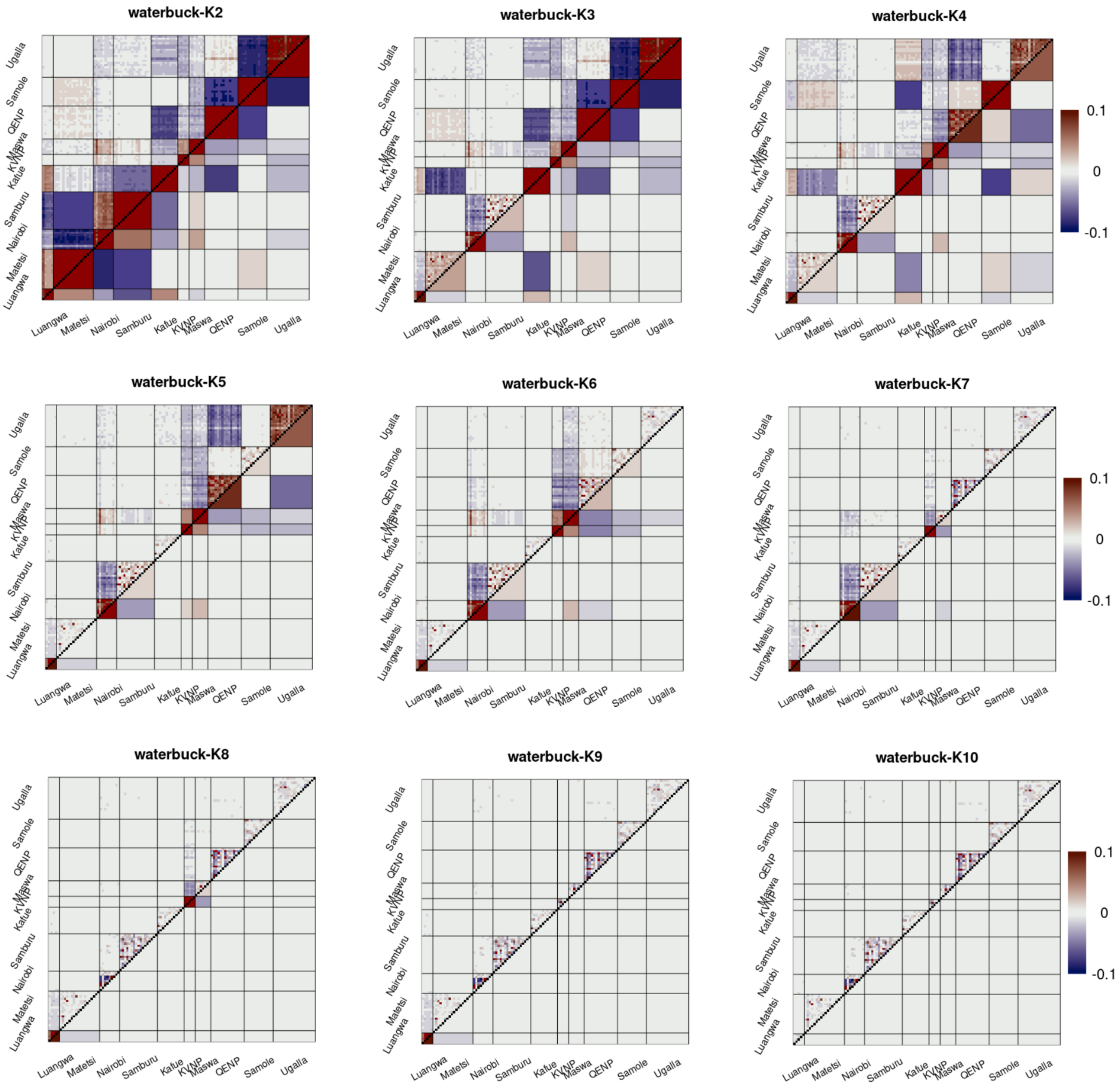

**Supplementary Figure 5.** Evaluation of NGSadmix results assuming from K = 2 to K = 10 as the correlation of residuals obtained with evalAdmix. The ordering of individuals is the same as in Fig. S4.

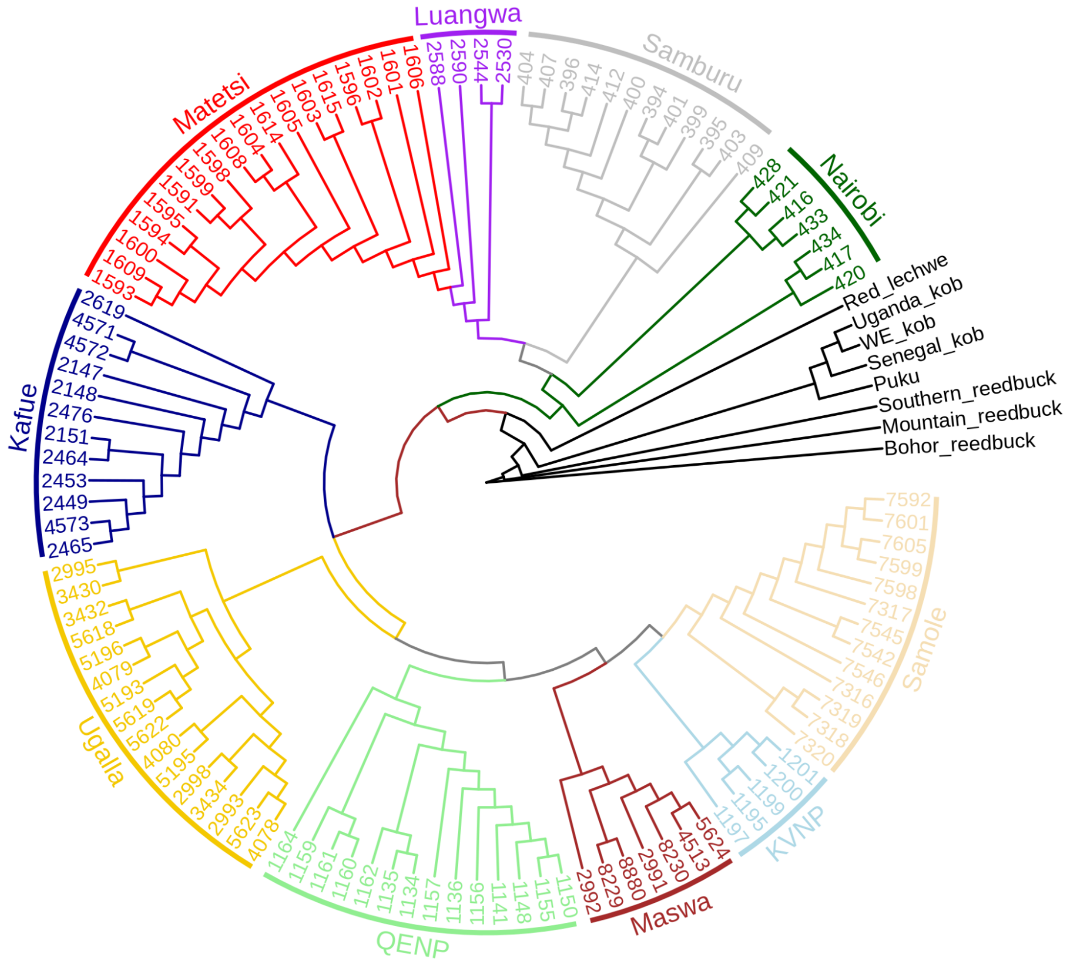

**Supplementary Figure 6.** Neighbor-joining tree based on an identity-by-state (IBS) matrix of 108 individuals (removing recently admixed samples identified by NGSadmix) and 8 outgroups. IBS was calculated in ANGSD and processed using the R package *ape* version 5.6-2[65].

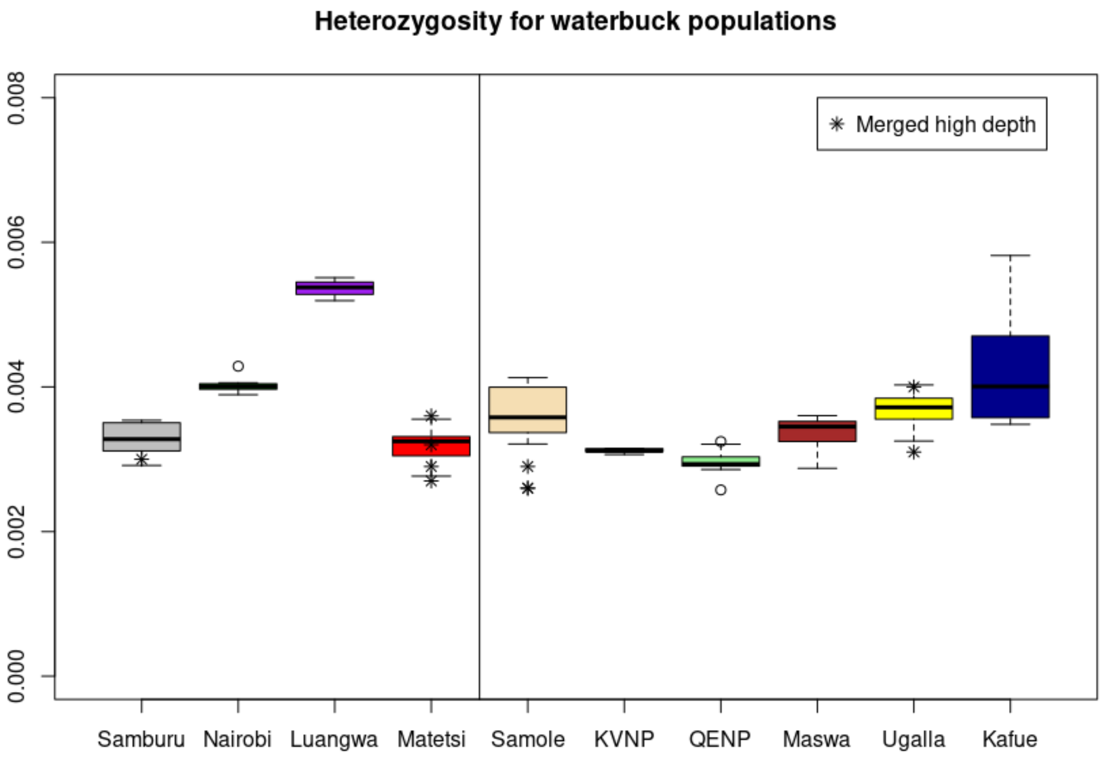

**Supplementary Figure 7**. Heterozygosity estimation using 108 waterbuck individuals without recently admixed samples based on genotype likelihoods (boxplots and small circles). The star symbols show the heterozygosity estimation based on called genotypes from high-depth individuals achieved by merging read data from duplicated samples. The overall similarity of estimates shows that the results are robust. The vertical line represents the split between subspecies; common waterbuck populations on the left and defassa waterbuck populations on the right.

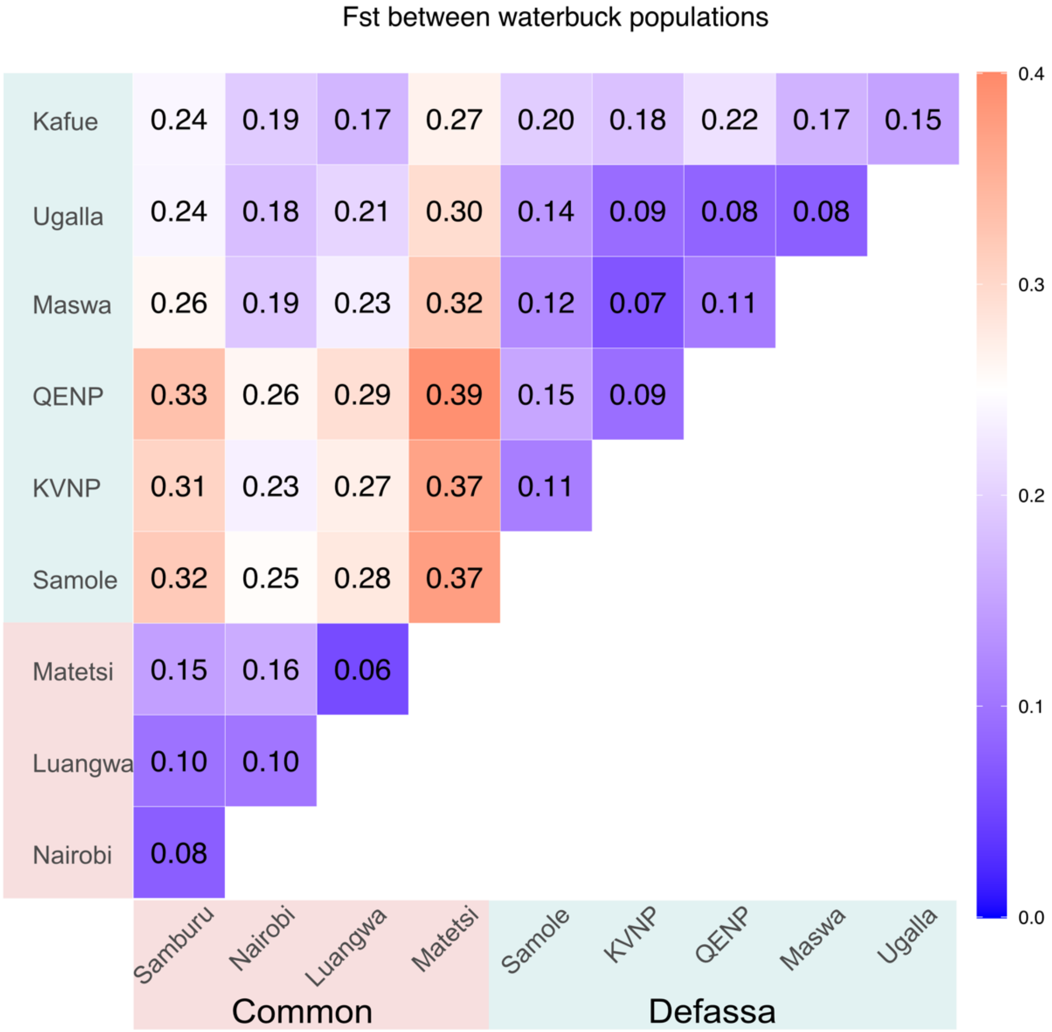

**Supplementary Figure 8**. Global genetic differentiation measured as $F_{st}$ values using Hudson's estimator between all population pairs estimated from the corresponding 2d-SFS.

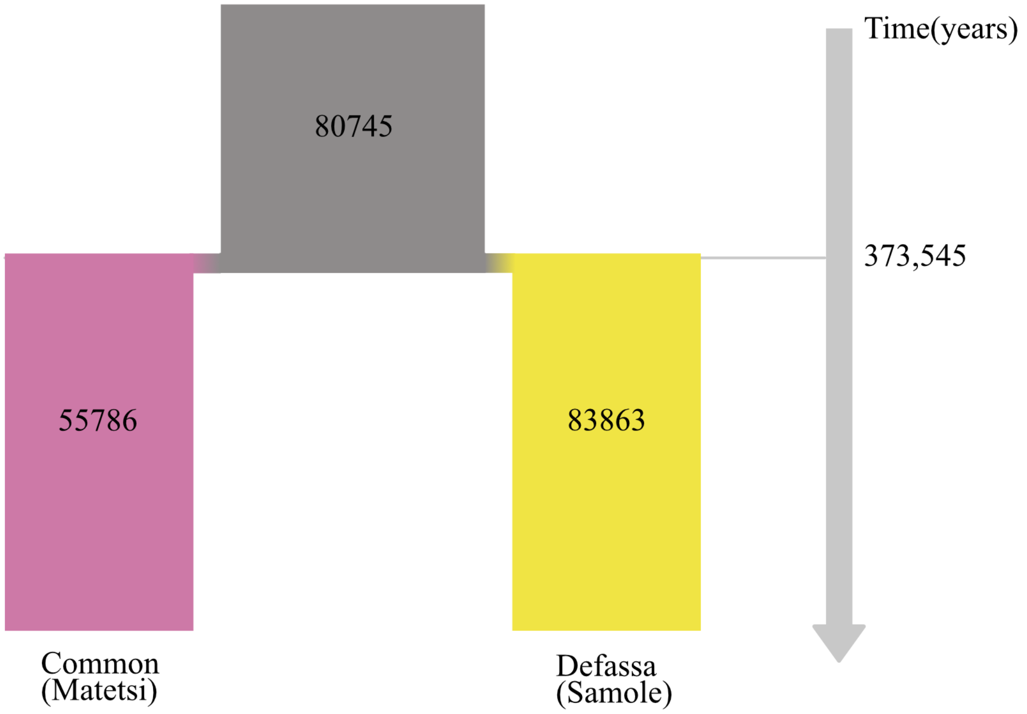

**Supplementary Figure 9**. Schematic diagram depicting the fastsimcoal2 demographic model and parameter point estimates, when considering a simple model to infer divergence time without gene flow between two subspecies. The Samole population was used to represent defassa waterbuck and Matetsi was used to represent common waterbuck, due to these populations having received the least gene flow from the other subspecies. All inferred demographic parameters and 95% confidence intervals are shown in Supplementary Table 5.

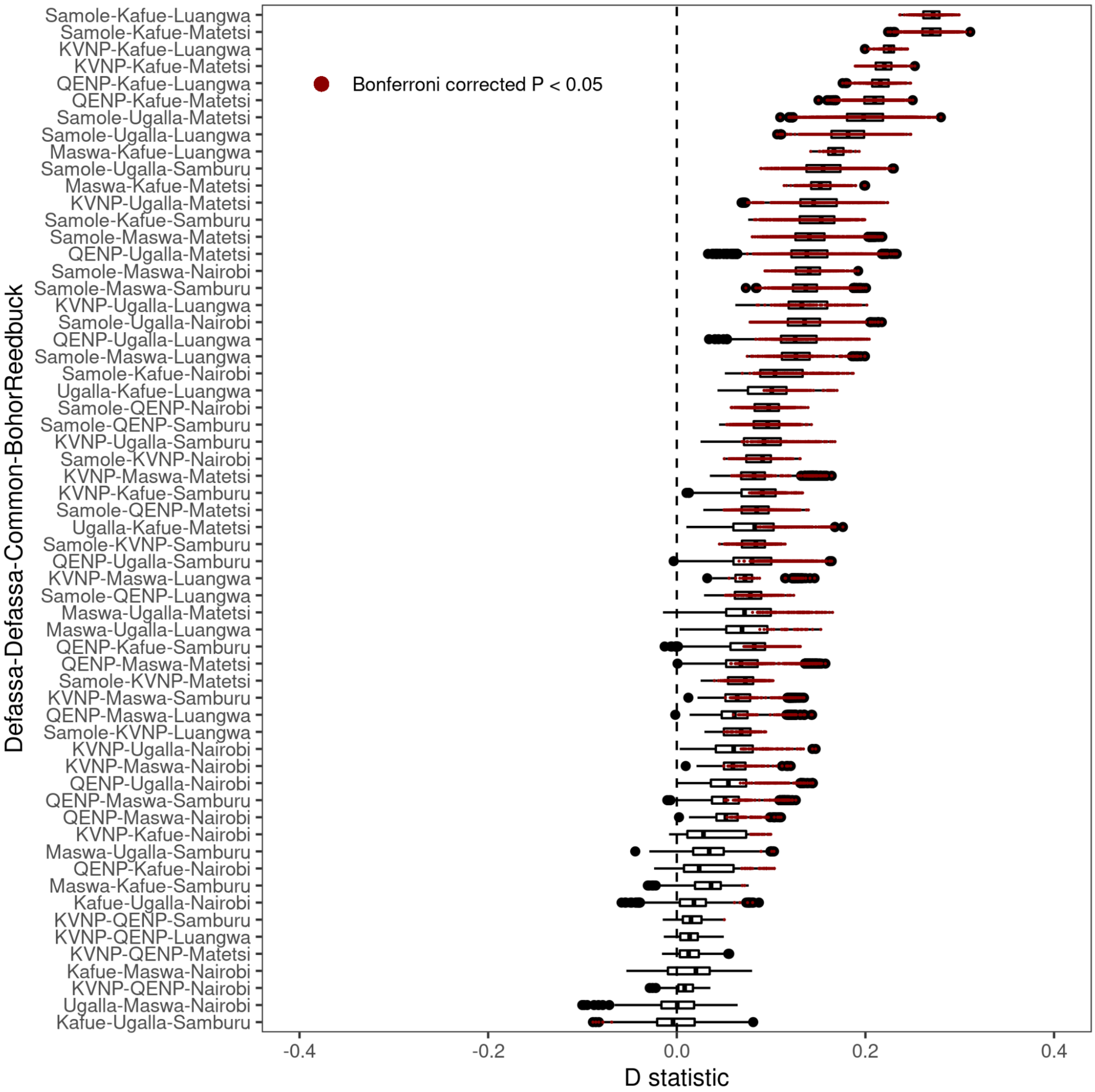

**Supplementary Figure 10.** D-statistics when using Bohor reedbuck as outgroup P4, various defassa waterbuck population pairs as P1, P2 and various common waterbuck populations as P3. The label in each row thereby denotes different combinations of populations in the order P1-P2-P3. D-statistics were estimated by single read sampling for all possible combinations of samples within the population trio using ANGSD. Red points represent a significant P value after Bonferroni correction calculated from Z scores.

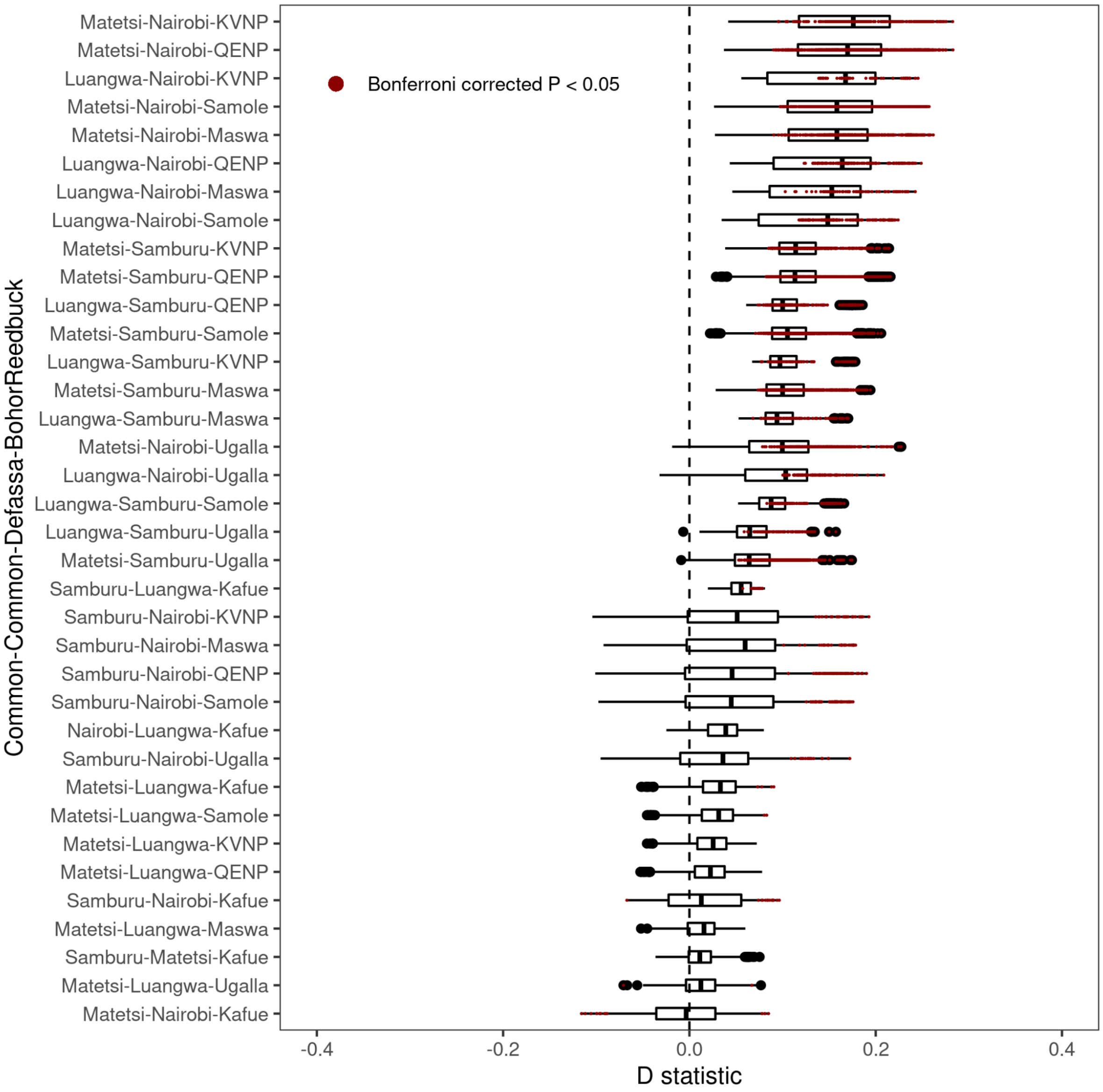

**Supplementary Figure 11.** As Fig. S10, but using Bohor reedbuck as outgroup P4, various common waterbuck populations as P1, P2 and various defassa waterbuck as P3. Red points represent the significant P value after Bonferroni correction calculated from Z scores.

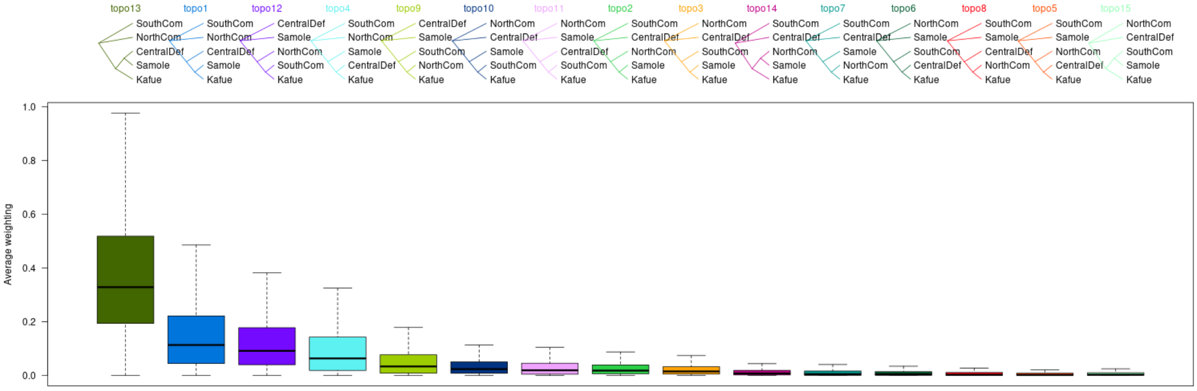

**Supplementary Figure 12.** The proportion of the 15 possible topologies in 100kb windows across the autosomal genome for five taxa, combining Nairobi+Samburu into one taxon (NorthCom); Matetsi+Luangwa into another taxon (SouthCom); Maswa+KVNP+QENP+Ugalla into another (CentralDef); and treating Kafue and Samole as separate taxa. Here we show the topology proportions along windows on chromosome 2 as an example.

**Supplementary Figure 13.** Plot of $F_{st}$ between two subspecies and variation in local topological relationships between two subspecies using TWISST, pairwise diversity, Tajima's D, and the pattern of localized linkage disequilibrium (LD) for defassa waterbuck (orange) and common waterbuck (light purple) of all 11 chromosomes where divergence islands were found.

**Supplementary Figure 14.** Plot of $F_{st}$ within all 14 outlying $F_{st}$ windows and its surrounding genomic region, together with its annotated protein coding genes.

**Supplementary Figure 15.** No signatures for reduced gene flow in 14 divergence islands. From left to right: population structure within each divergence island inferred by PCAngsd; the information of subspecies and populations that grouped individuals based on 5 bins split by PC1 in A belonging to; and heterozygosity estimation using realSFS in ANGSD by grouping individuals based on 5 bins split by PC1 in A.

**Supplementary Figure 16.** Highlighting D-statistics values of 10 "disparate" samples identified in Supplementary Figure 15.

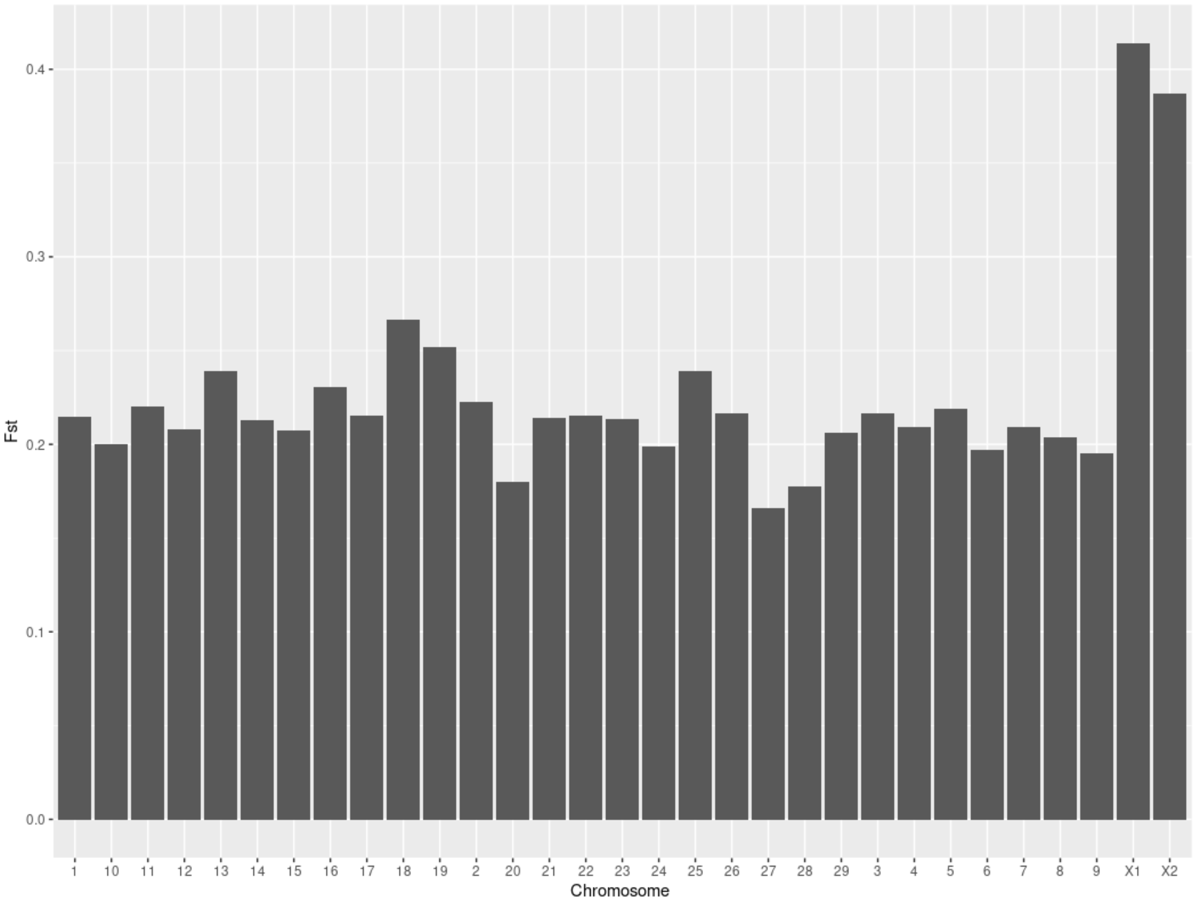

**Supplementary Figure 17.** Average Fst across each chromosome.

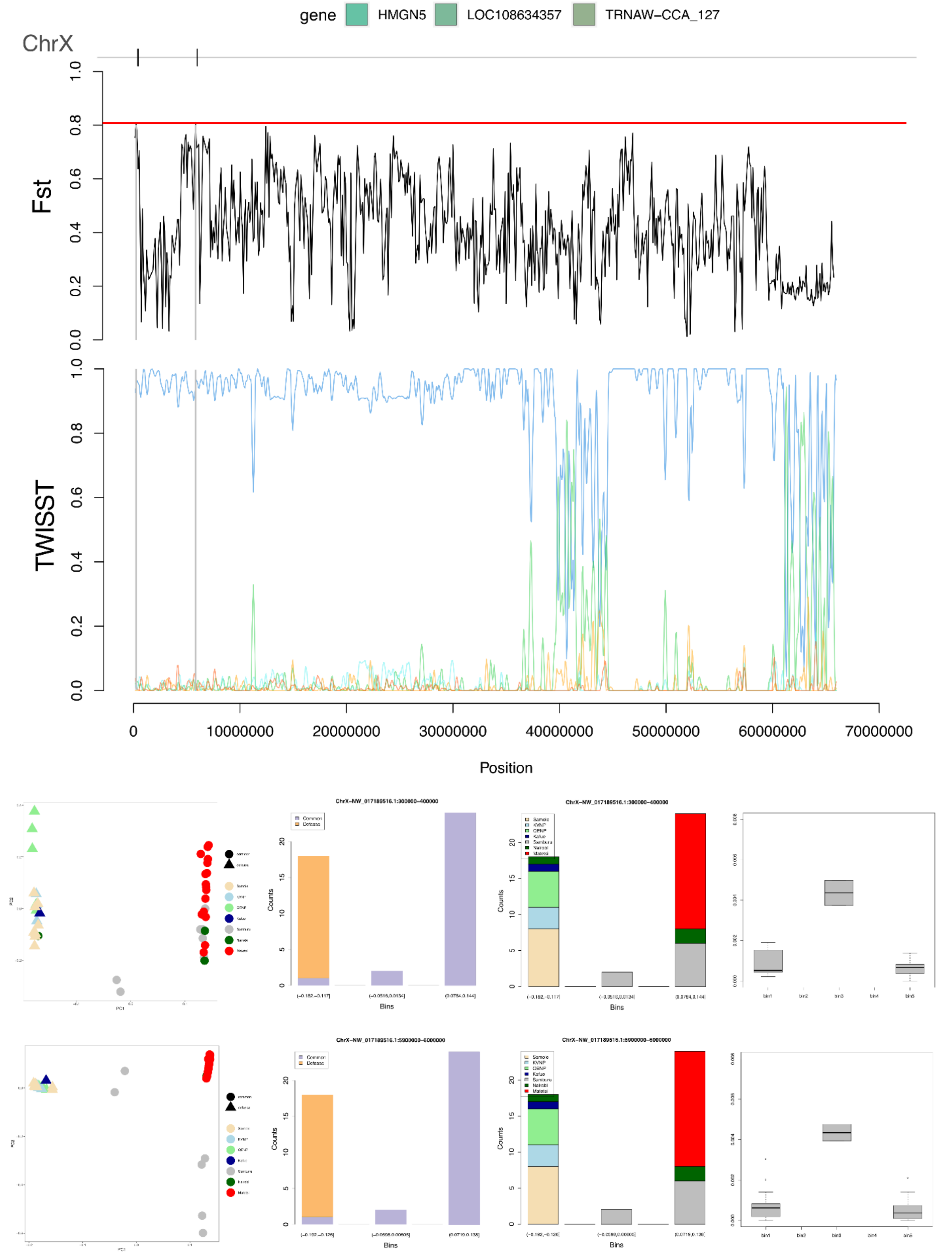

**Supplementary Figure 18.** Identification of two possible divergence islands on the X chromosome, using the distribution of $F_{st}$ across windows on the X chromosomal scaffolds to define top 0.1% outlier windows.

### Supplementary Materials Tables

**Supplementary Table 1.** Data sets after different steps of quality control.

| Population | Original samples | QC control Step1 (Basic mapping statistics filtering; Error rates: Heterozygosity; Duplicates filtering based on King) | QC control Step1 QC control Step2 (NGSrelate within each location) | QC control Step1 QC control Step2 removing admixed samples (NGSadmix) |
|---|---|---|---|---|
| **Samole** | 26 | 15 | 13 | 13 |
| **Maswa** | 7 | 7 | 7 | 7 |
| **KVNP** | 7 | 5 | 5 | 5 |
| **QENP** | 15 | 15 | 15 | 14 |
| **Ugalla** | 21 | 20 | 19 | 16 |
| **Kafue** | 15 | 12 | 12 | 12 |
| **Samburu** | 18 | 17 | 17 | 12 |
| **Matetsi** | 23 | 20 | 18 | 18 |
| **Nairobi** | 9 | 9 | 9 | 7 |
| **Luangwa** | 4 | 4 | 4 | 4 |
| **Total** | 145 | 124 | 119 | 108 |
| **Dataset type** | A | B | C | D |

**Supplementary Table 2.** Sample inclusion after different steps of sample filtering.

**Supplementary Table 3.** Summary statistics for sites filtering.

| | **Defassa Waterbuck (ref)** **88935 scaffolds** **sites:2895549025** | | **Goat (ref)** **29907 scaffolds** **sites:2922813246** | |
|---|---|---|---|---|
| | sites removed | sites remained (proportion%) | sites removed | sites remained (proportion%) |

| | | | | |
|---|---|---|---|---|
| **Sex Linked And Abnormal Scaffolds** | 250’295’945 | 2’645’253’080<br><br>(91%) | 133’653’460<br><br>(Keep 4Chr, so all 29 were maintained) | 2’789’159’786<br><br>(95%) |
| **RepeatMasker(ref)** | 622’184’946 | 2’273’364’079<br><br>(79%) | 1’295’118’187 | 1’627’695’059<br><br>(56%) |
| **Mappibility(ref)** | 580’144’885 | 2’315’404’140<br><br>(80%) | 273’776’835 | 2’649’036’411<br><br>(91%) |
| **ExcessHeterozygosity** | 49’862’593 | 2’845’686’432<br><br>(98%) | 138’264’407 | 2’784’548’839<br><br>(95%) |
| **Depth**<br><br>**(375<x<median * 1.5 )** | 1’246’315’525 | 1’649’233’500<br><br>(57%) | 1’355’482’338 | 1’567’330’908<br><br>(54%) |
| **All_combined**<br><br>**(all scaffolds)** | 1’748’758’452 | 114’6790’573<br><br>(40%) | 1’991’560’501 | 931’252’745<br><br>(32%) |
| **All_combined**<br><br>**(with scaffolds length > 100kb)** | 1’775’681’421 | 1’119’867’604<br><br>(39%) | - | - |
| **All_combined**<br><br>**(with Goat only include 29 Chr)** | - | - | 2’003’874’943 | 918’938’303<br><br>(31%) |

**Supplementary Table 4.** Summary table with distance to potential pedigrees and summary indices using the recent admixture inference method (Garcia- Erill et al. in prep) applied to all admixed samples identified by NGSAdmix assuming K=10. The “admixture index” can be interpreted as the number of generations since admixture, and is only shown for samples verified to be the result of recent admixture using this method. The “Distance to pedigree x” is the Jensen-Shannon distance of the estimated proportions to the expected under an independent pedigree, and to the pedigree identified as the most compatible recent admixture pedigrees.

| SampleID | Inconsistency Index | Distance to independent pedigree | Distance to pedigree 1 | Distance to pedigree 2 | Admixture index |
|---|---|---|---|---|---|
| 415_Samburu | 0.06 | 0.00 | 0.09 | 0.21 | |

| | | | | | |
|---|---|---|---|---|---|
| 2535_Luangwa | 0.09 | 0.00 | 0.02 | 0.10 | |
| 1133_QENP | 0.05 | 0.00 | 0.15 | 0.20 | |
| 393_Samburu | 0.06 | 0.00 | 0.10 | 0.21 | |
| 427_Nairobi | 0.23 | 0.22 | 0.11 | 0.13 | 5.16 |
| 2994_Ugalla | 0.17 | 0.00 | 0.24 | 0.26 | |
| 2996_Ugalla | 0.05 | 0.01 | 0.17 | 0.21 | |
| 392_Samburu | 0.08 | 0.00 | 0.25 | 0.27 | |
| 398_Samburu | 0.02 | 0.28 | 0.02 | 0.13 | 3.07 |
| 429_Nairobi | 0.12 | 0.00 | 0.04 | 0.12 | |
| 408_Samburu | 0.06 | 0.00 | 0.10 | 0.21 | |

**Supplementary Table 5.** Inferred parameters of demographic history between defassa and common waterbuck under a simple demographic model of population divergence without gene flow. We assumed a mutation rate of 1.43e-8 per site per generation and 7.1 years of generation time.

| **Parameters** | | | | **Point estimation** | **95% CI**[a] | | |
|---|---|---|---|---|---|---|---|
| | | | | | **Lower bound** | **Upper bound** | **Standard Error** |
| **Effective population sizes** | **Ancest** | Common (Samole) & Defassa(Matesis) | NANC | 80745 | 78044 | 82272 | 11330 |
| | **Population** | Common (Samole) | NPOPC | 55786 | 54516 | 57421 | 8261 |
| | | Defassa(Matesis) | NPOPD | 83863 | 82644 | 88254 | 14367 |
| **Time** | **Divergence** | Common (Samole) & | TDIV | 373545 | 364791 | 385803 | 57908 |

| | | |
|---|---|---|
| | | Defassa( Matesis) |

[a] Non-parametric bootstrap estimates obtained by 100 resampled SFS using jackknife sampling with same parameters setting up for the original data sets.

**Supplementary Table 6.** Summary statistics for 14 Fst outlier windows.

| **Divergence island** | **Position** | **Twisst** (quantile) | **Pairwise diversity** (quantile) (Common) | **Pairwise diversity** (quantile) (Defassa) | **Tajima's D** (quantile) (Common) | **Tajima's D** (quantile) (Defassa) | **LD** (quantile) (Common) | **LD** (quantile) (Defassa) | **Haplotype frequency**[a] |
|---|---|---|---|---|---|---|---|---|---|
| **Chr2 (NC_030809.1)** | 134700000-135900000 | 1.000 | 0.0426 | 0.0685 | 0.0474 | 0.0815 | 0.893 | 0.921 | 0.611 |
| **Chr5 (NC_030812.1)** | 55400000-55500000 | 1.000 | 0.0105 | 0.0165 | 0.00673 | 0.0661 | 0.749 | 0.941 | - |
| **Chr6 (NC_030813.1)** | 400000-1000000 | 0.881 | 0.193 | 0.00914 | 0.215 | 0.00341 | 0.0310 | - | 0.347 |
| **Chr7 (NC_030814.1)** | 56200000-56300000 | 0.941 | 0.0289 | 0.0687 | 0.0190 | 0.0907 | 0.715 | 0.982 | 0.384 |
| **Chr13 (NC_030820.1)** | 22400000-22500000 | 0.818 | 0.0501 | 0.0751 | 0.0204 | 0.0675 | 0.910 | 0.977 | 0.389 |
| **Chr14 (NC_030821.1)** | 26500000-26600000 | 0.810 | 0.0557 | 0.133 | 0.0411 | 0.161 | 0.992 | 0.0407 | 0.412 |
| **Chr16 (NC_030823.1)** | 41400000-41500000 | 1.000 | 0.0884 | 0.0308 | 0.0474 | 0.00810 | 0.879 | 0.736 | - |
| **Chr16 (NC_030823.1)** | 43200000-43300000 | 0.851 | 0.00254 | 0.00104 | 0.00175 | 0.000623 | 0.926 | 0.982 | 0.380 |

| | | | | | | | | | |
|---|---|---|---|---|---|---|---|---|---|
| **Chr16 (NC_030823.1)** | 49300000-49600000 | 0.962 | 0.00985 | 0.0460 | 0.00116 | 0.0168 | 0.964 | 0.980 | 0.384 |
| **Chr17 (NC_030824.1)** | 35800000-36000000 | 0.965 | 0.00557 | 0.0856 | 0.00116 | 0.173 | 0.986 | 0.355 | 0.435 |
| **Chr18 (NC_030825.1)** | 38300000-39300000 | 0.929 | 0.0133 | 0.0159 | 0.0184 | 0.0225 | 0.910 | 0.943 | - |
| **Chr22 (NC_030829.1)** | 17000000-17100000 | 1.000 | 0.00212 | 0.0298 | 0.00237 | 0.0312 | 0.886 | 0.980 | - |
| **Chr25 (NC_030832.1)** | 1100000-1700000 | 1.000 | 0.0354 | 0.0168 | 0.0190 | 0.00578 | 0.869 | 0.976 | - |
| **Chr25 (NC_030832.1)** | 3200000-3300000 | 0.961 | 0.00262 | 0.000790 | 0.00378 | 0.00112 | 0.965 | 0.994 | 0.611 |

[a] The “haplotype” frequency was only calculated for windows showing three distinct clusters.
