## SupplementaryFigure13 for "Persistent gene flow suggests an absence of reproductive isolation in an African antelope speciation model"

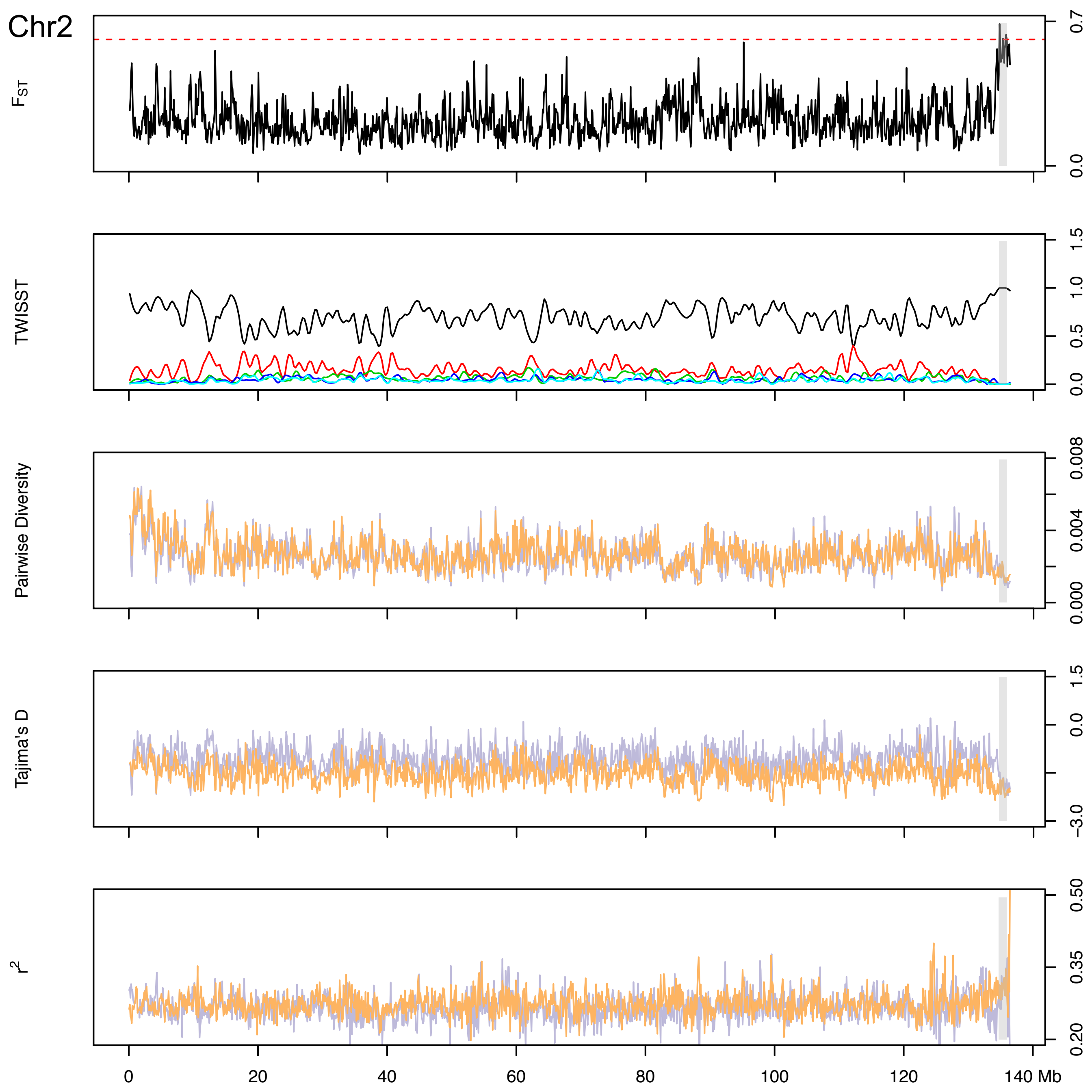

Chr2
$F_{ST}$
0.0
0.7
TWISST
0.0
0.5
1.0
1.5
Pairwise Diversity
0.000
0.004
0.008
Tajima's D
−3.0
−1.5
0.0
1.5
$r^2$
0.20
0.35
0.50
0
20
40
60
80
100
120
140 Mb

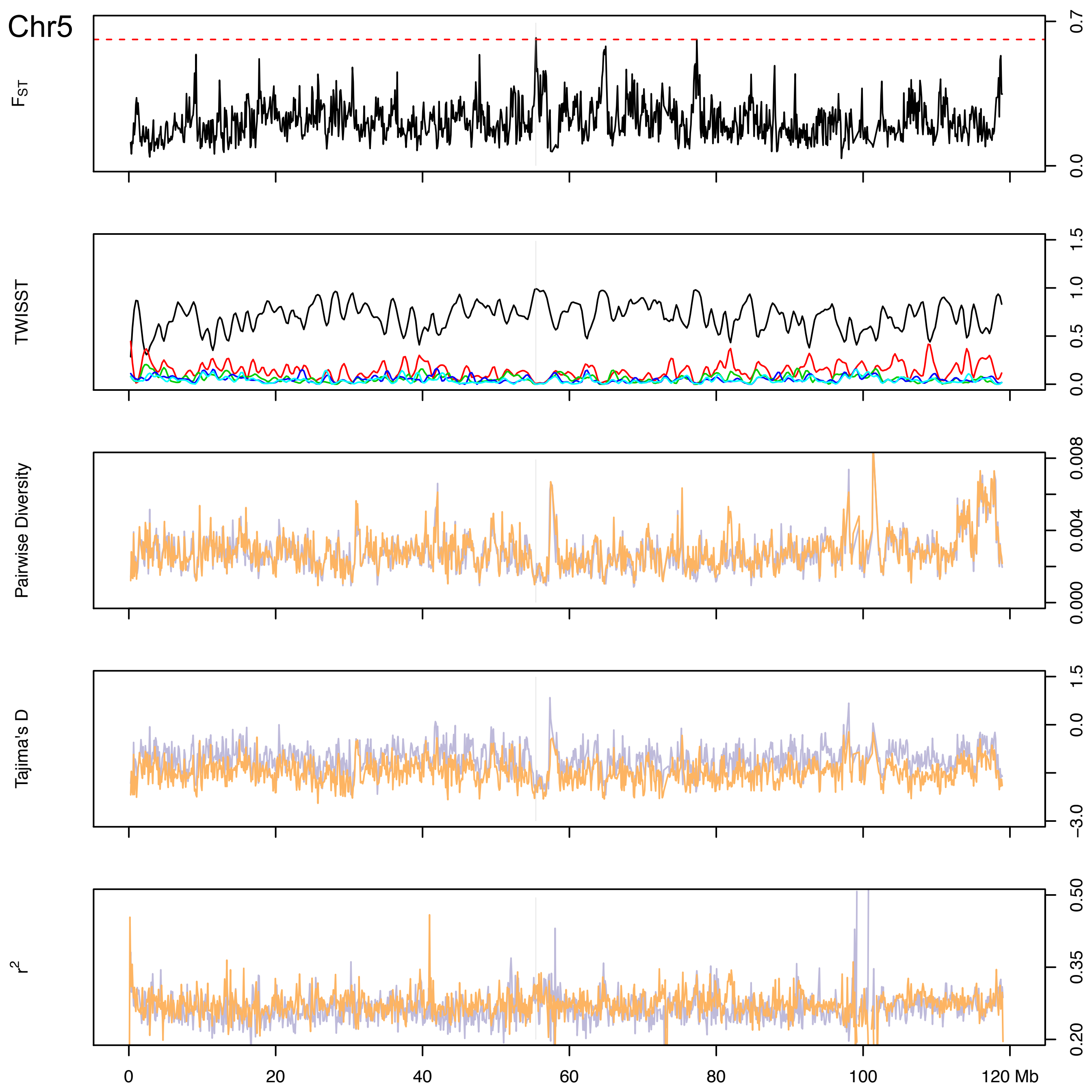

Chr5
$F_{ST}$
0.0
0.7
TWISST
0.0
0.5
1.0
1.5
Pairwise Diversity
0.000
0.004
0.008
Tajima's D
−3.0
0.0
1.5
$r^2$
0.20
0.35
0.50
0
20
40
60
80
100
120 Mb

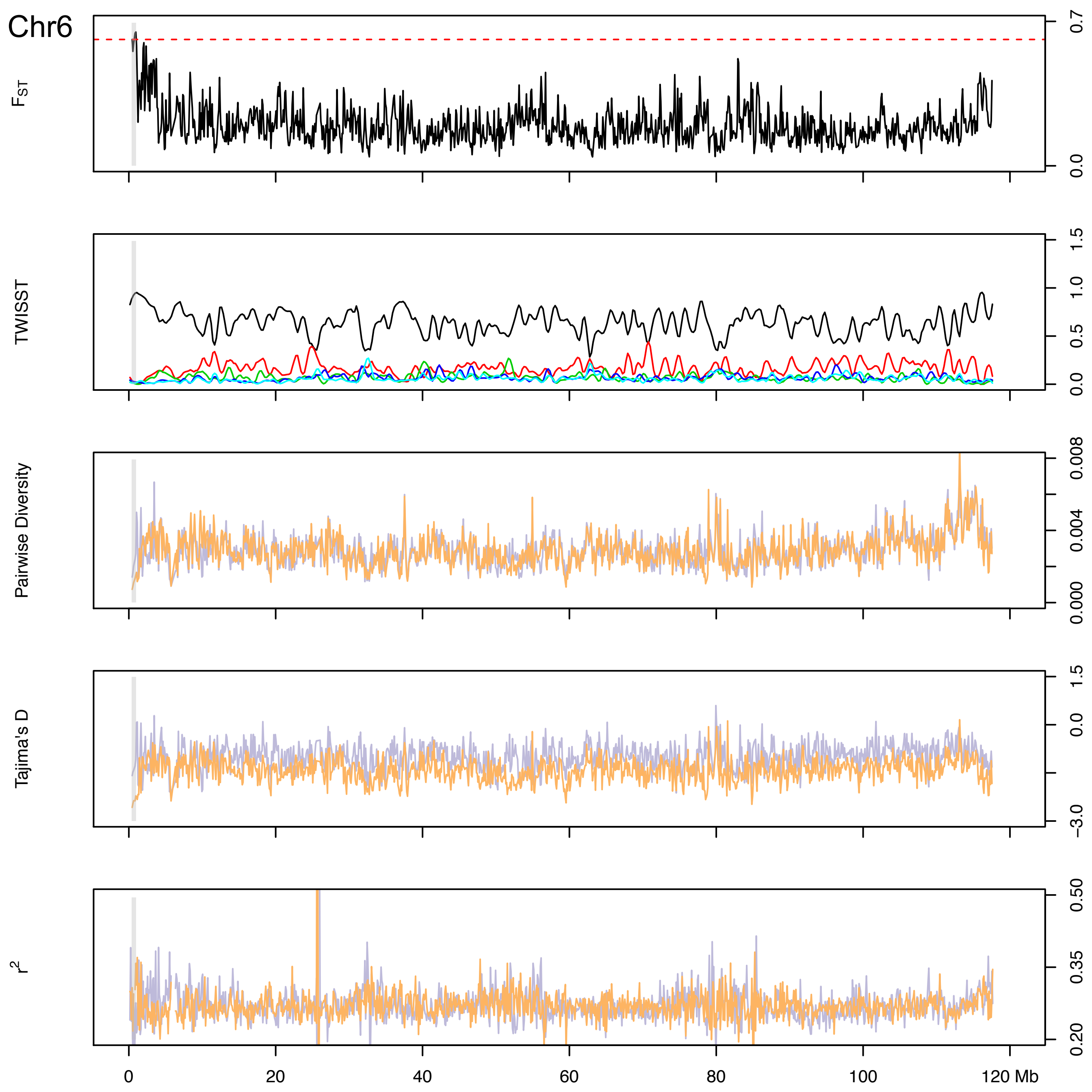

Chr6
F_ST
0.7
0.0
TWISST
1.5
1.0
0.5
0.0
Pairwise Diversity
0.008
0.004
0.000
Tajima's D
1.5
0.0
−3.0
r²
0.50
0.35
0.20
0
20
40
60
80
100
120 Mb

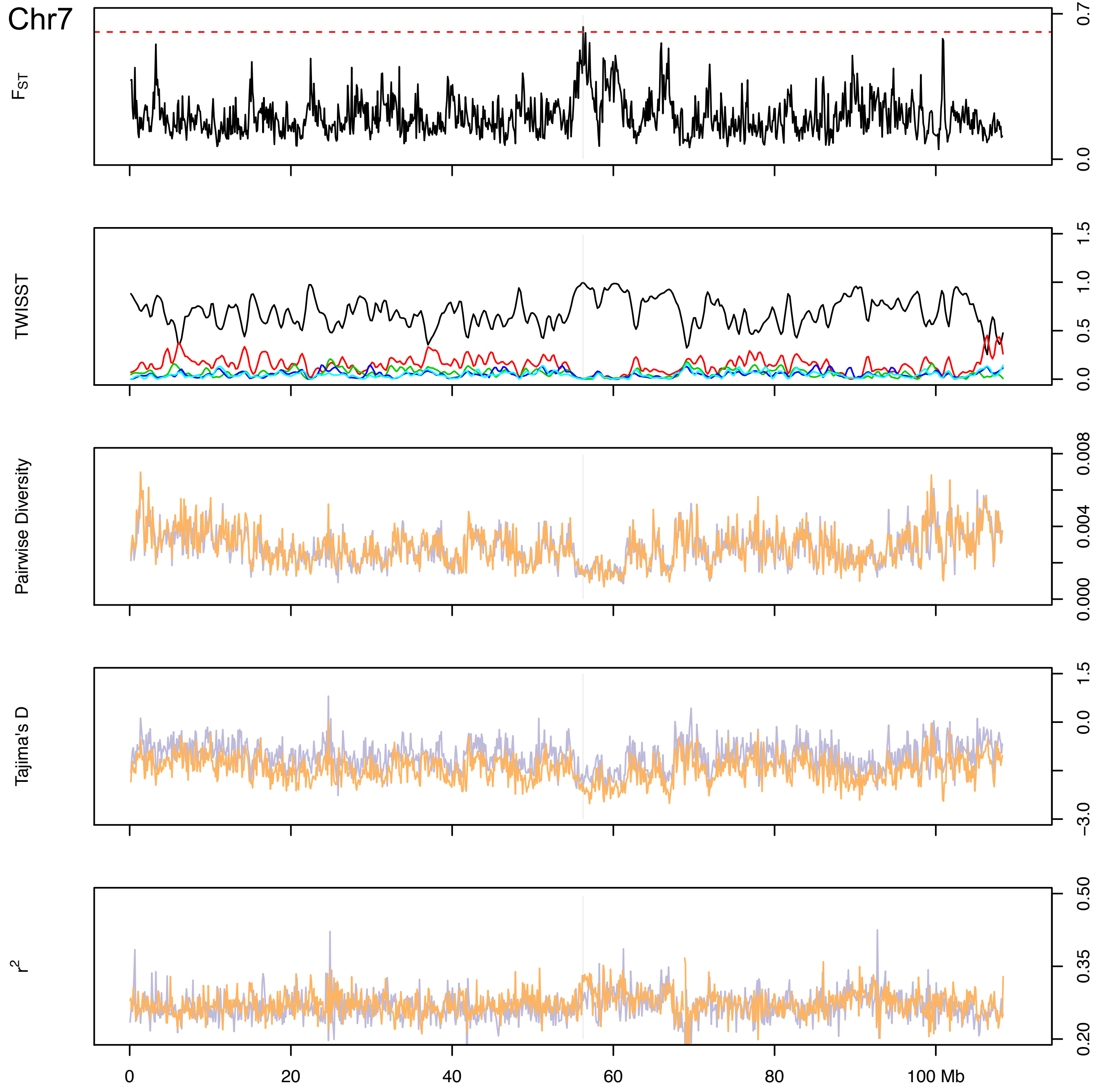

Chr7
$F_{ST}$
0.7
0.0
TWISST
1.5
1.0
0.5
0.0
Pairwise Diversity
0.008
0.004
0.000
Tajima's D
1.5
0.0
−3.0
$r^2$
0.50
0.35
0.20
0
20
40
60
80
100 Mb

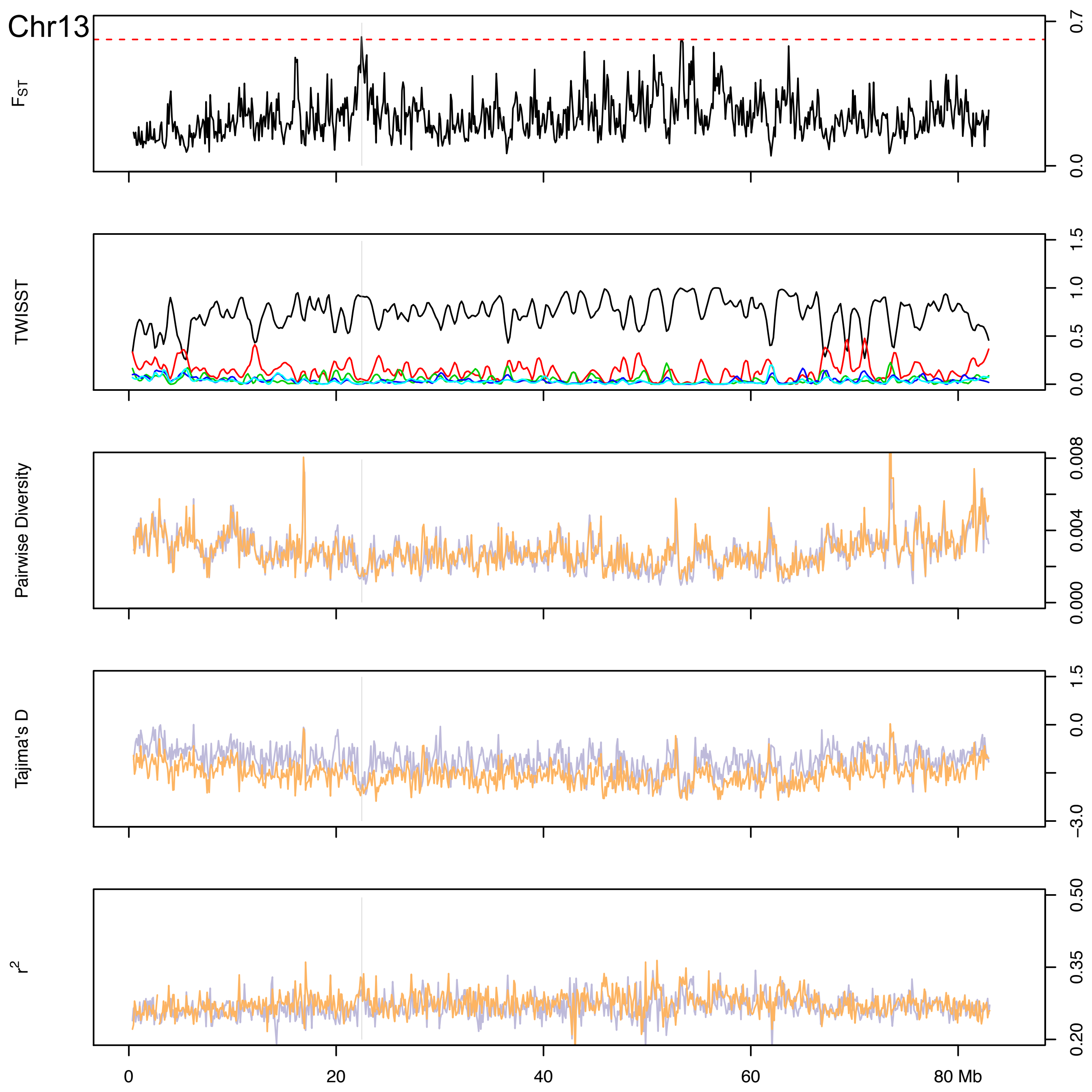
Chr13
$F_{ST}$
0.7
0.0
TWISST
1.5
1.0
0.5
0.0
Pairwise Diversity
0.008
0.004
0.000
Tajima's D
1.5
0.0
−3.0
$r^2$
0.50
0.35
0.20
0
20
40
60
80 Mb

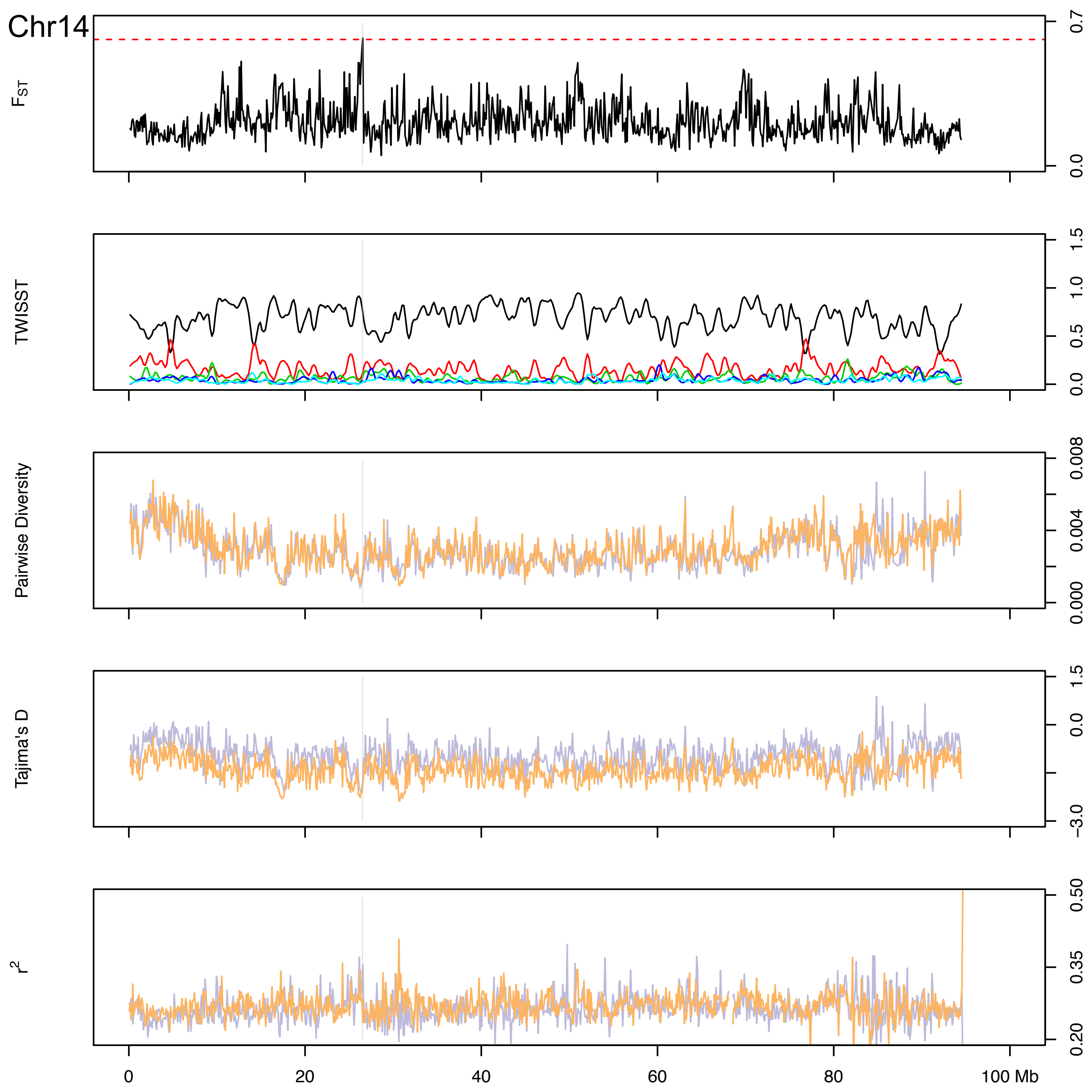

Chr14
$F_{ST}$
0.7
0.0
TWISST
1.5
1.0
0.5
0.0
Pairwise Diversity
0.008
0.004
0.000
Tajima's D
1.5
0.0
−3.0
$r^2$
0.50
0.35
0.20
0
20
40
60
80
100 Mb

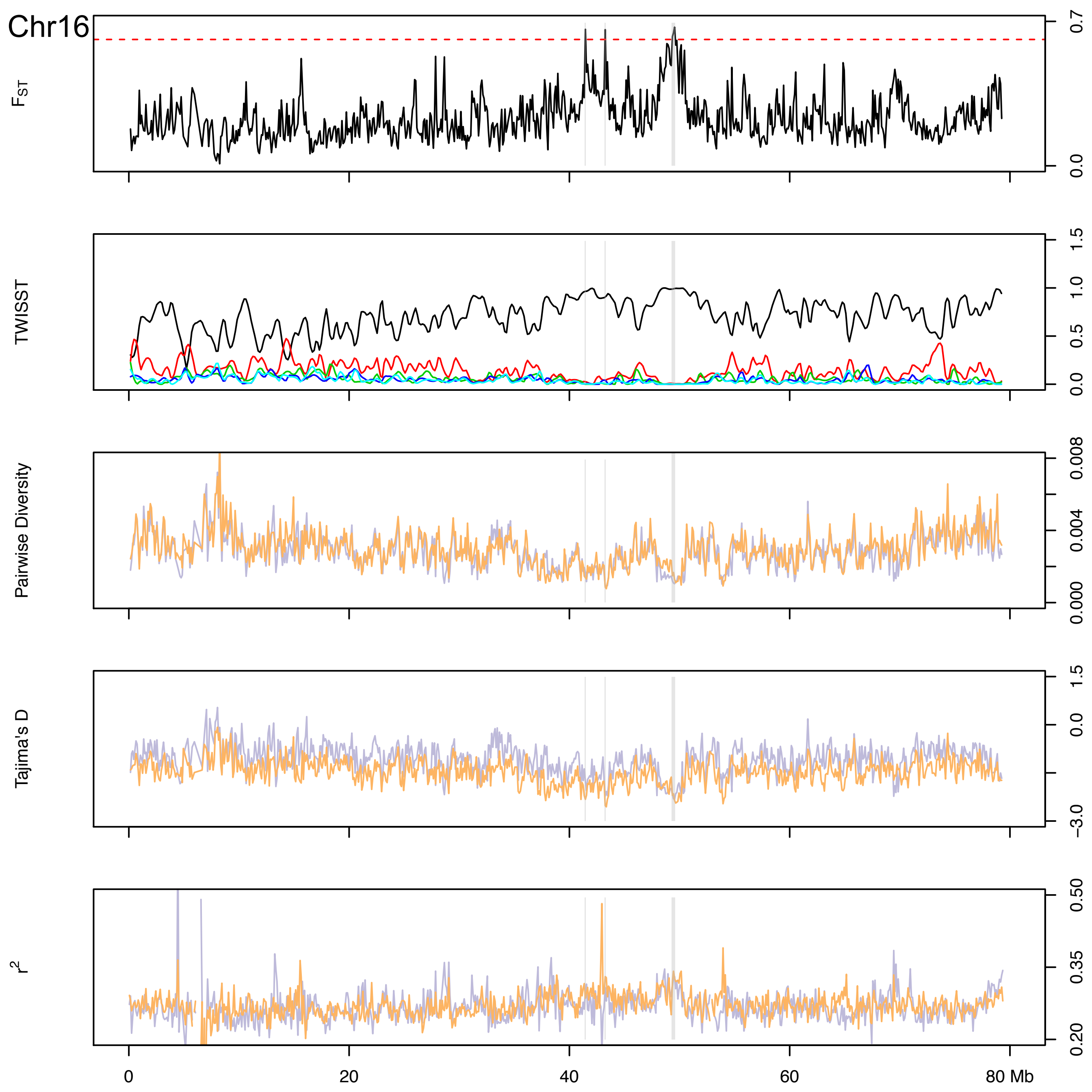

Chr16
$F_{ST}$
0.7
0.0
TWISST
1.5
1.0
0.5
0.0
Pairwise Diversity
0.008
0.004
0.000
Tajima's D
1.5
0.0
−3.0
$r^2$
0.50
0.35
0.20
0
20
40
60
80 Mb

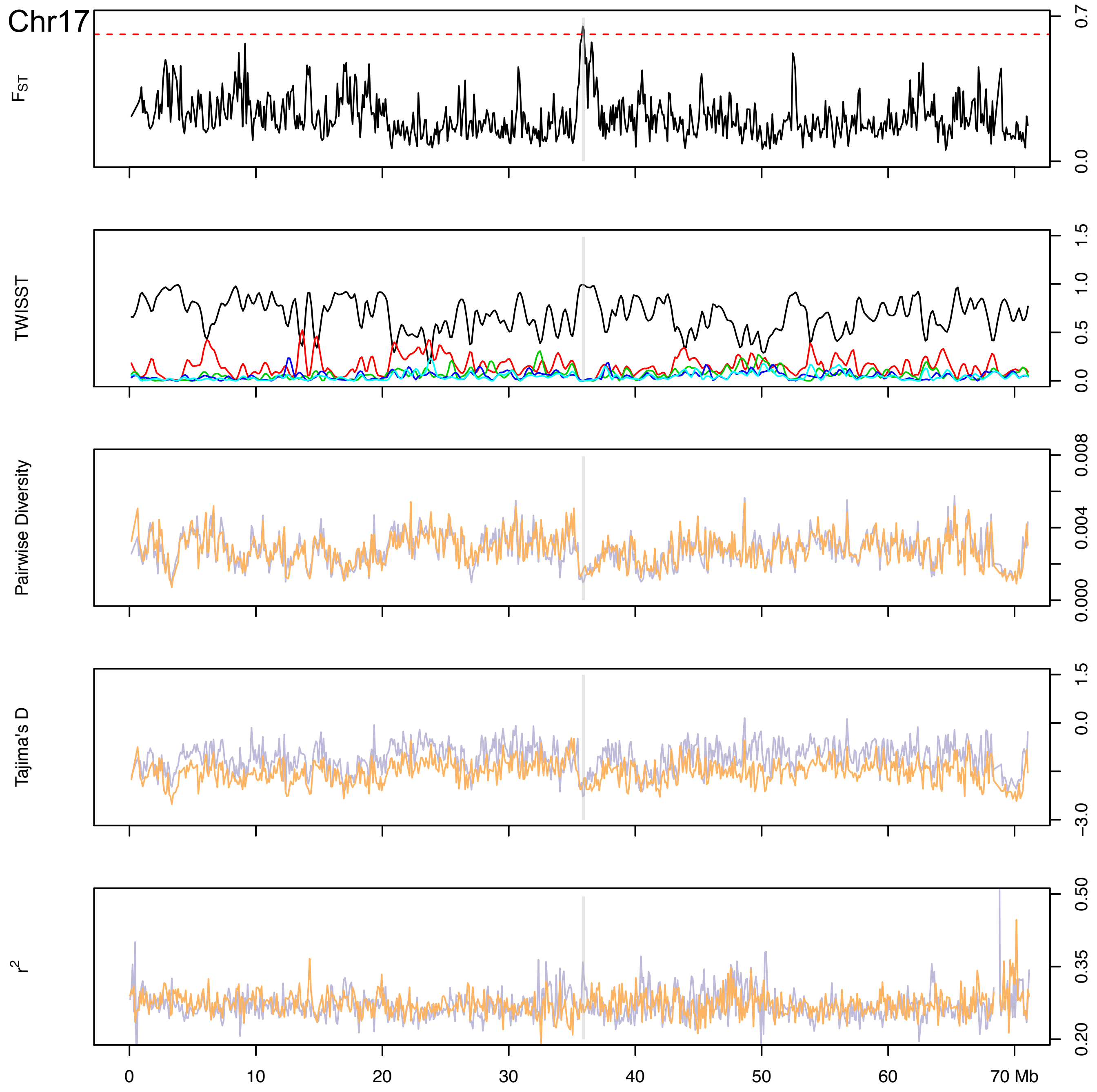

Chr17
F_ST
TWISST
Pairwise Diversity
Tajima's D
r^2
0.7
0.0
1.5
1.0
0.5
0.0
0.008
0.004
0.000
1.5
0.0
−3.0
0.50
0.35
0.20
0
10
20
30
40
50
60
70 Mb

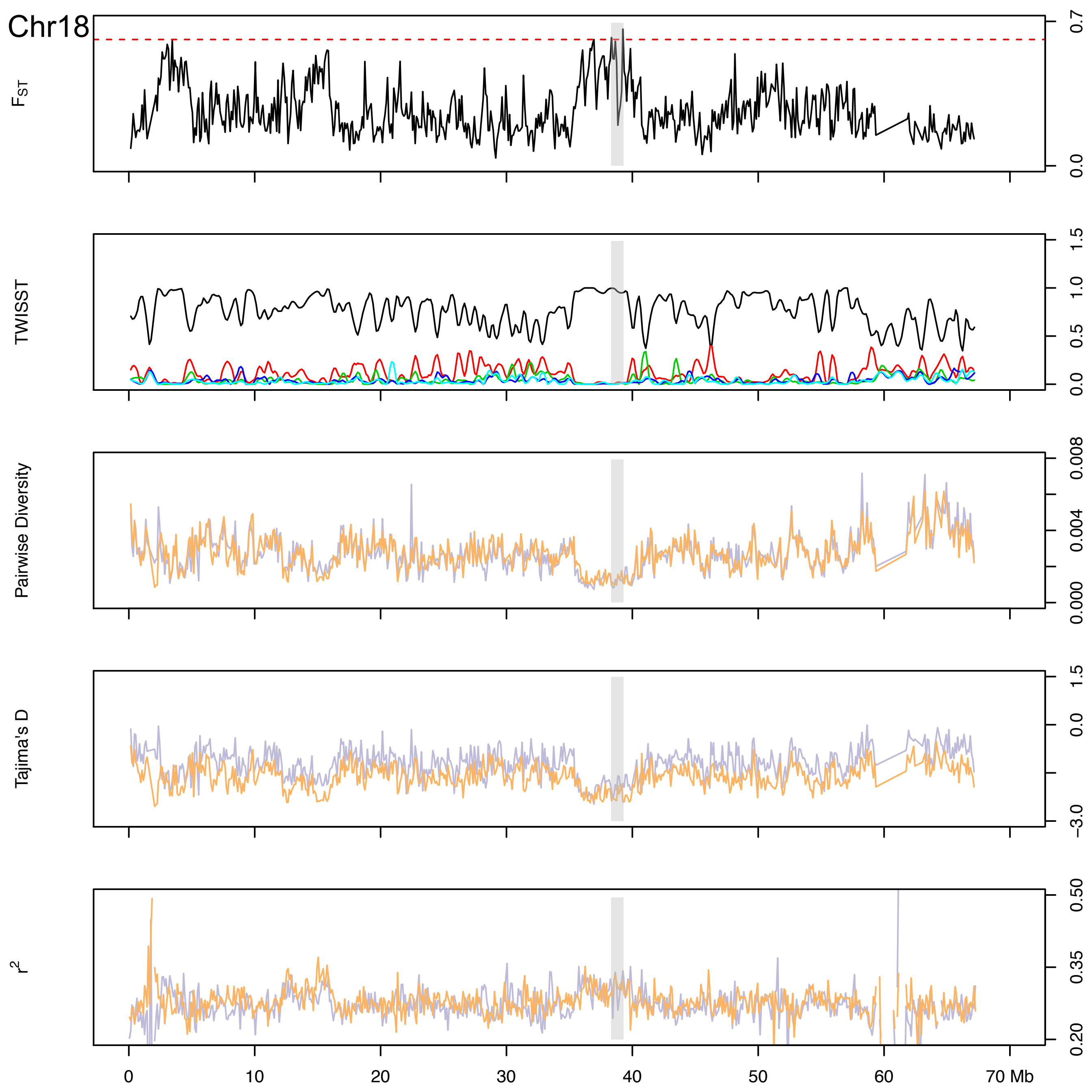

Chr18
FST
TWISST
Pairwise Diversity
Tajima's D
r2
0.7
0.0
1.5
1.0
0.5
0.0
0.008
0.004
0.000
1.5
0.0
−3.0
0.50
0.35
0.20
0
10
20
30
40
50
60
70 Mb

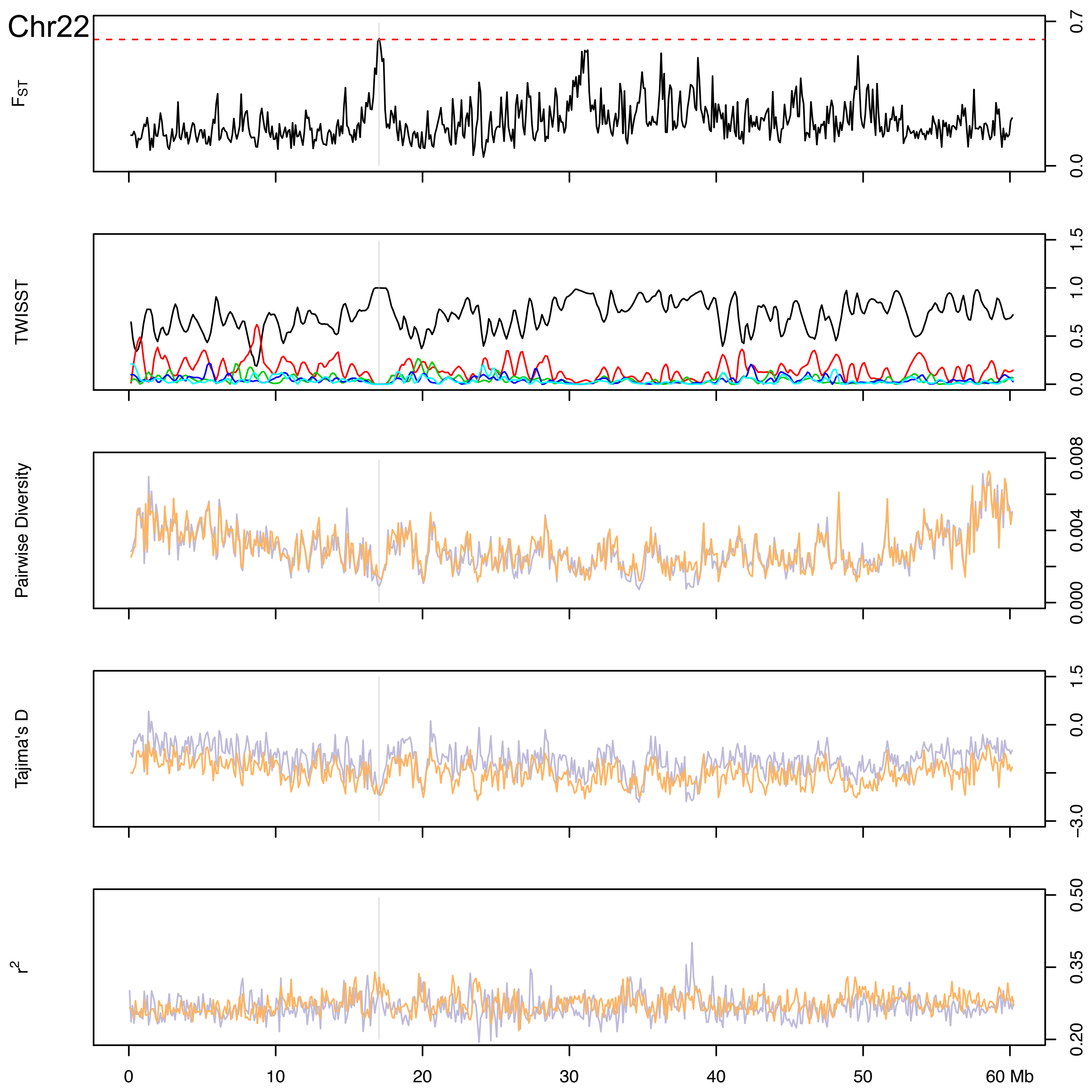

Chr22
$F_{ST}$
0.7
0.0
TWISST
1.5
1.0
0.5
0.0
Pairwise Diversity
0.008
0.004
0.000
Tajima's D
1.5
0.0
−3.0
$r^2$
0.50
0.35
0.20
0
10
20
30
40
50
60 Mb

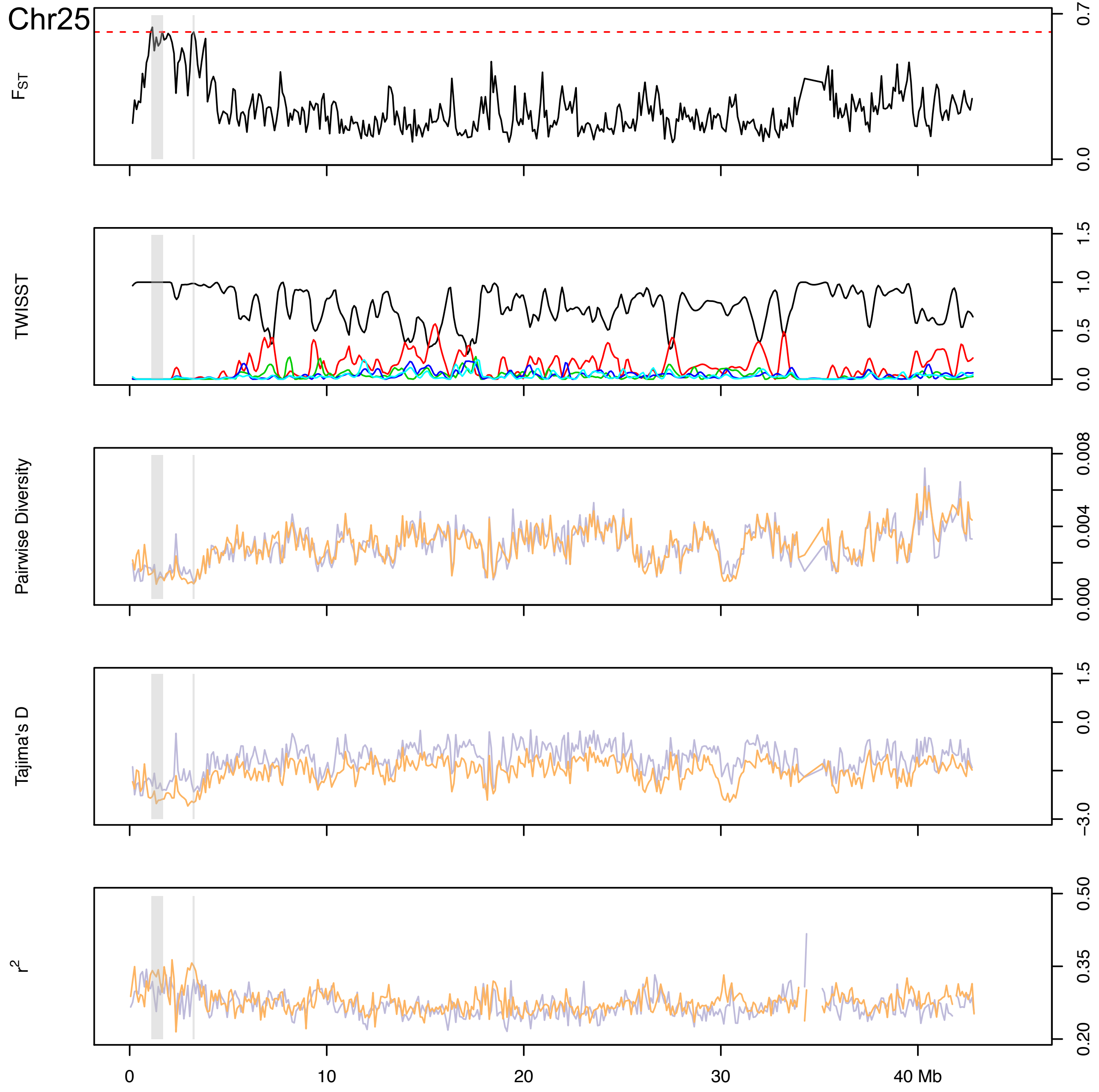
Chr25
$F_{ST}$
0.7
0.0
TWISST
1.5
1.0
0.5
0.0
Pairwise Diversity
0.008
0.004
0.000
Tajima's D
1.5
0.0
−3.0
$r^2$
0.50
0.35
0.20
0
10
20
30
40 Mb
