## SupplementaryFigure14 for "Persistent gene flow suggests an absence of reproductive isolation in an African antelope speciation model"

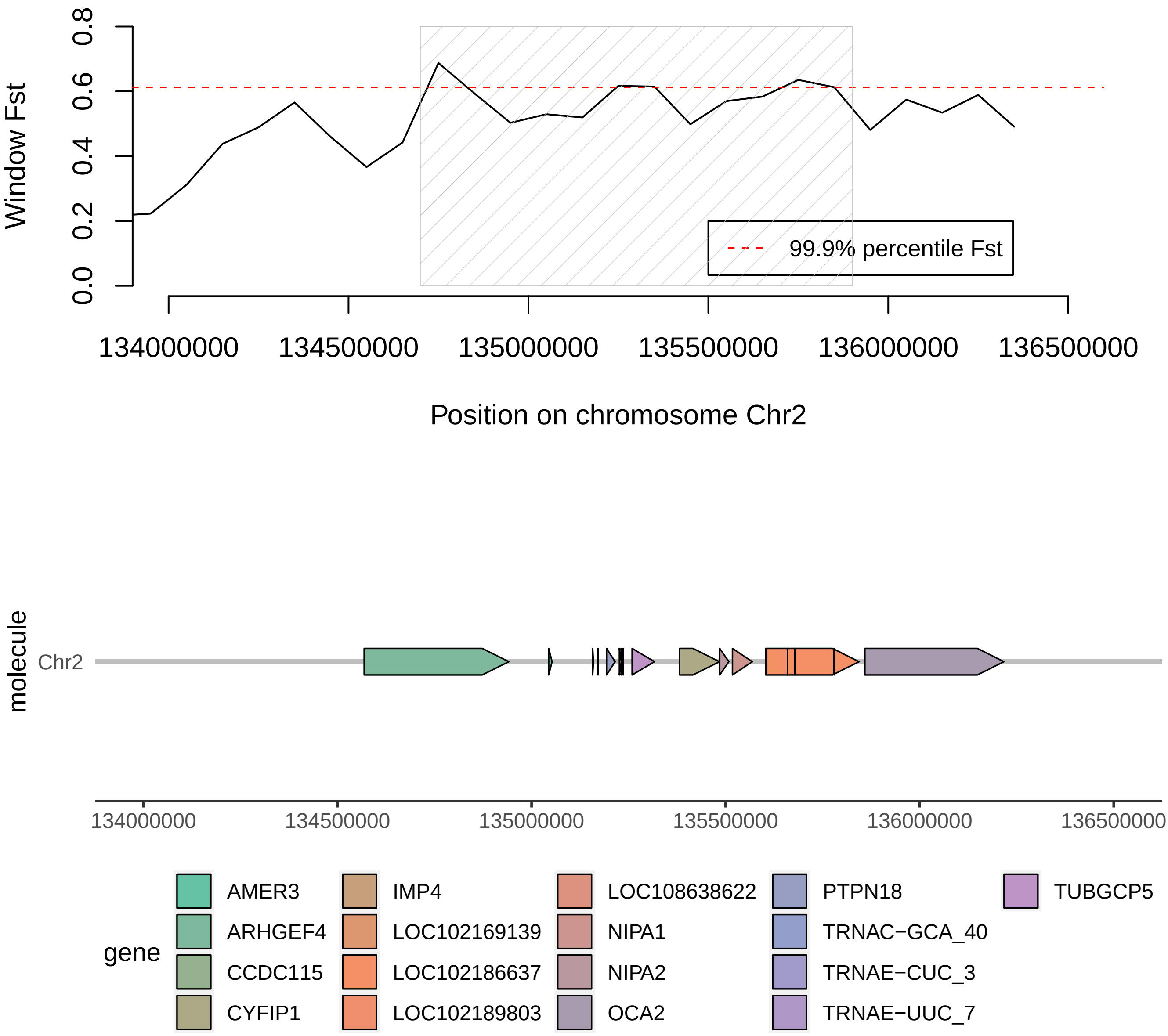

Window Fst
0.0
0.2
0.4
0.6
0.8
99.9% percentile Fst
134000000
134500000
135000000
135500000
136000000
136500000
Position on chromosome Chr2
molecule
Chr2
134000000
134500000
135000000
135500000
136000000
136500000
gene
AMER3
ARHGEF4
CCDC115
CYFIP1
IMP4
LOC102169139
LOC102186637
LOC102189803
LOC108638622
NIPA1
NIPA2
OCA2
PTPN18
TRNAC−GCA_40
TRNAE−CUC_3
TRNAE−UUC_7
TUBGCP5

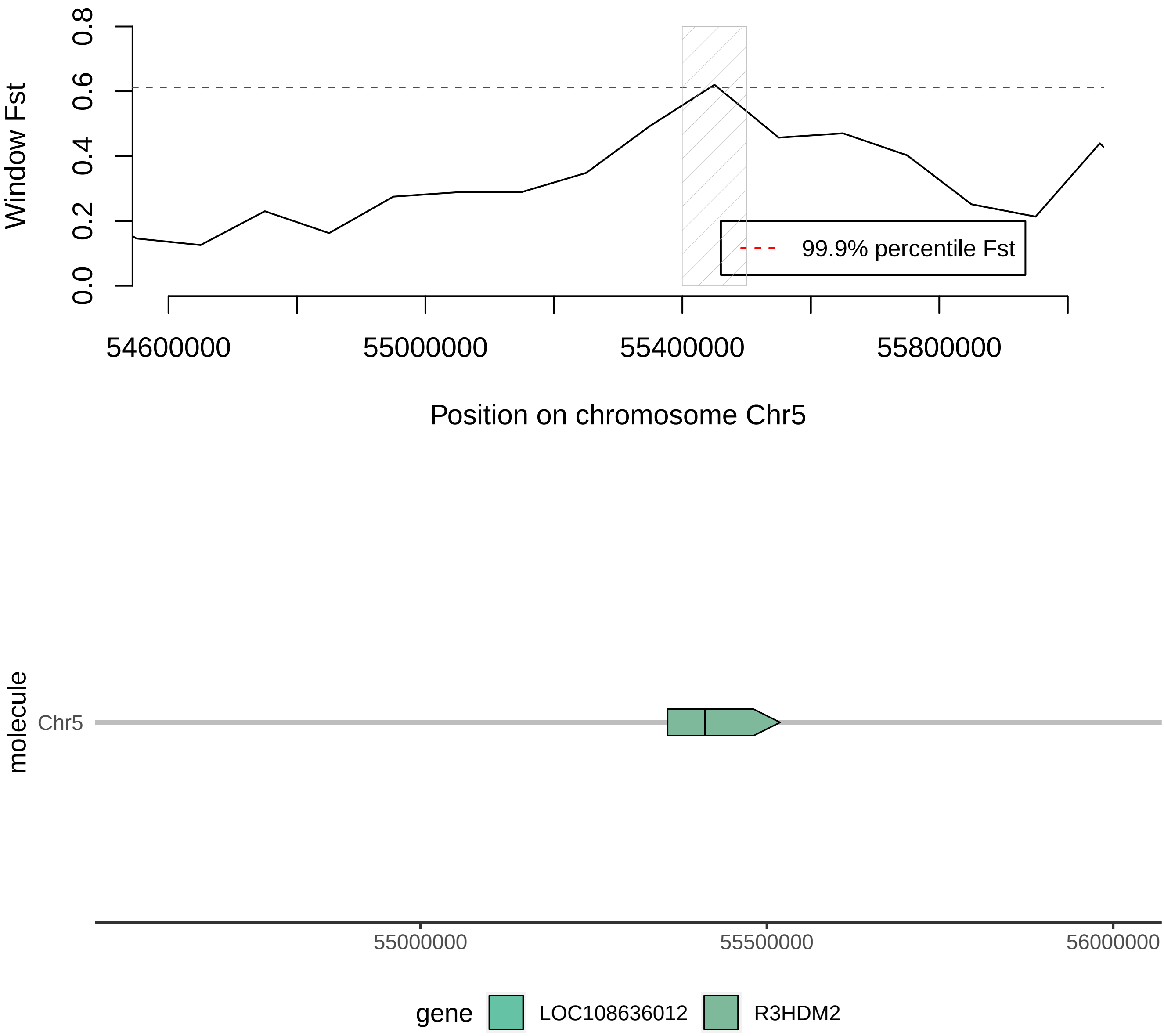
Window Fst
0.0
0.2
0.4
0.6
0.8
99.9% percentile Fst
54600000
55000000
55400000
55800000
Position on chromosome Chr5
molecule
Chr5
55000000
55500000
56000000
gene
LOC108636012
R3HDM2

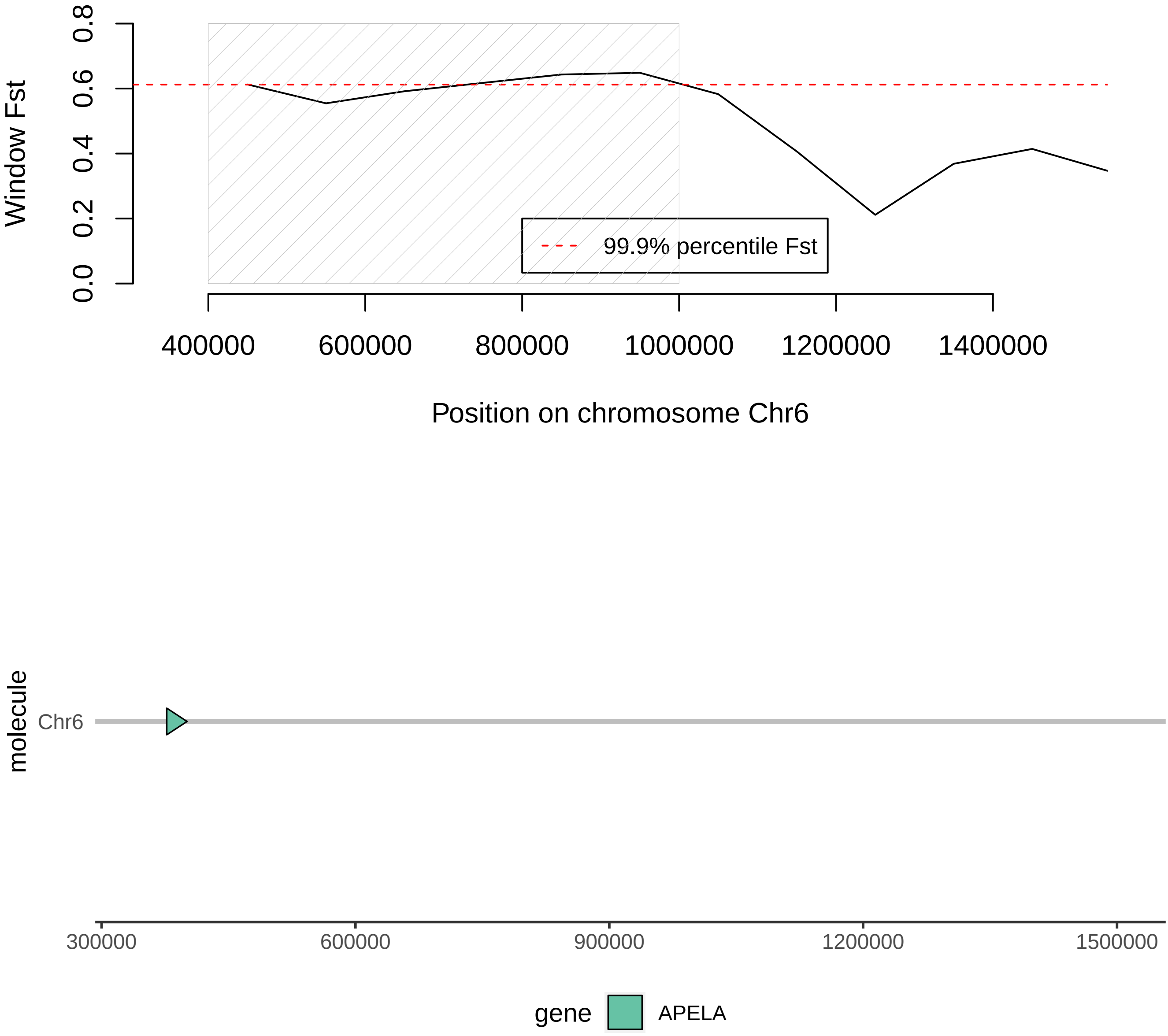

Window Fst
0.0
0.2
0.4
0.6
0.8
99.9% percentile Fst
400000
600000
800000
1000000
1200000
1400000
Position on chromosome Chr6
molecule
Chr6
300000
600000
900000
1200000
1500000
gene
APELA

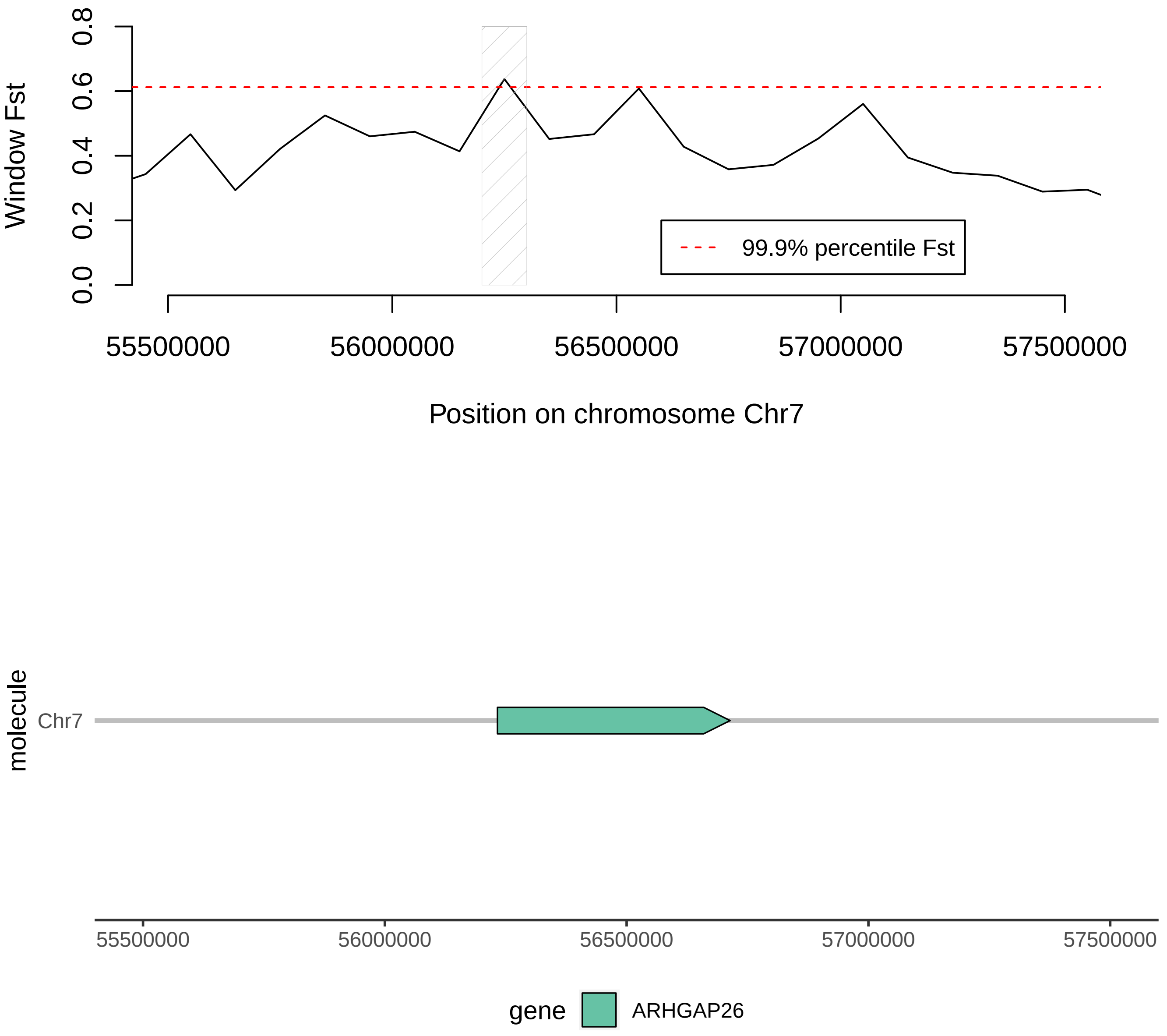

Window Fst
0.0
0.2
0.4
0.6
0.8
99.9% percentile Fst
55500000
56000000
56500000
57000000
57500000
Position on chromosome Chr7
molecule
Chr7
55500000
56000000
56500000
57000000
57500000
gene
ARHGAP26

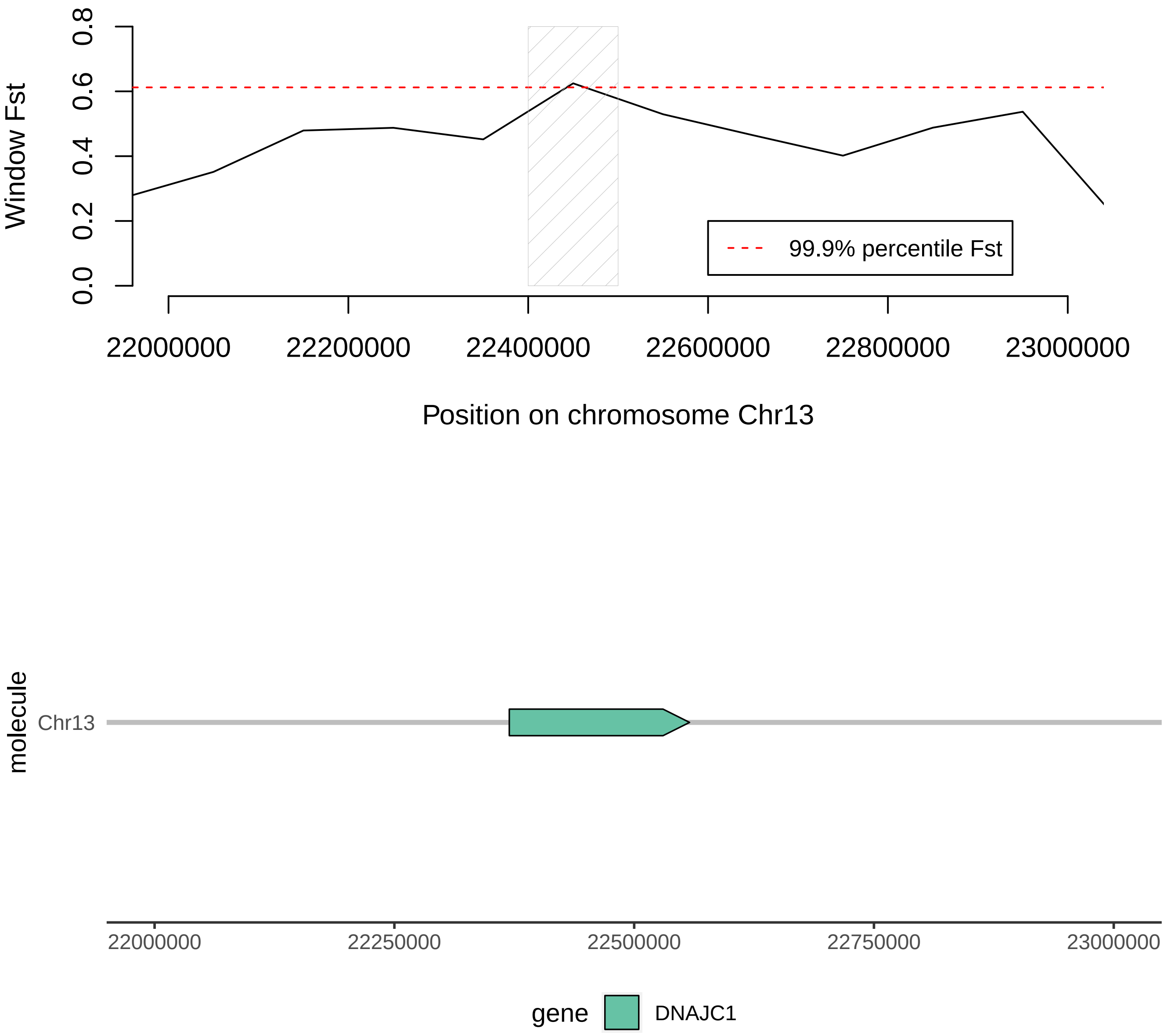

Window Fst
0.0
0.2
0.4
0.6
0.8
99.9% percentile Fst
22000000
22200000
22400000
22600000
22800000
23000000
Position on chromosome Chr13
molecule
Chr13
22000000
22250000
22500000
22750000
23000000
gene
DNAJC1

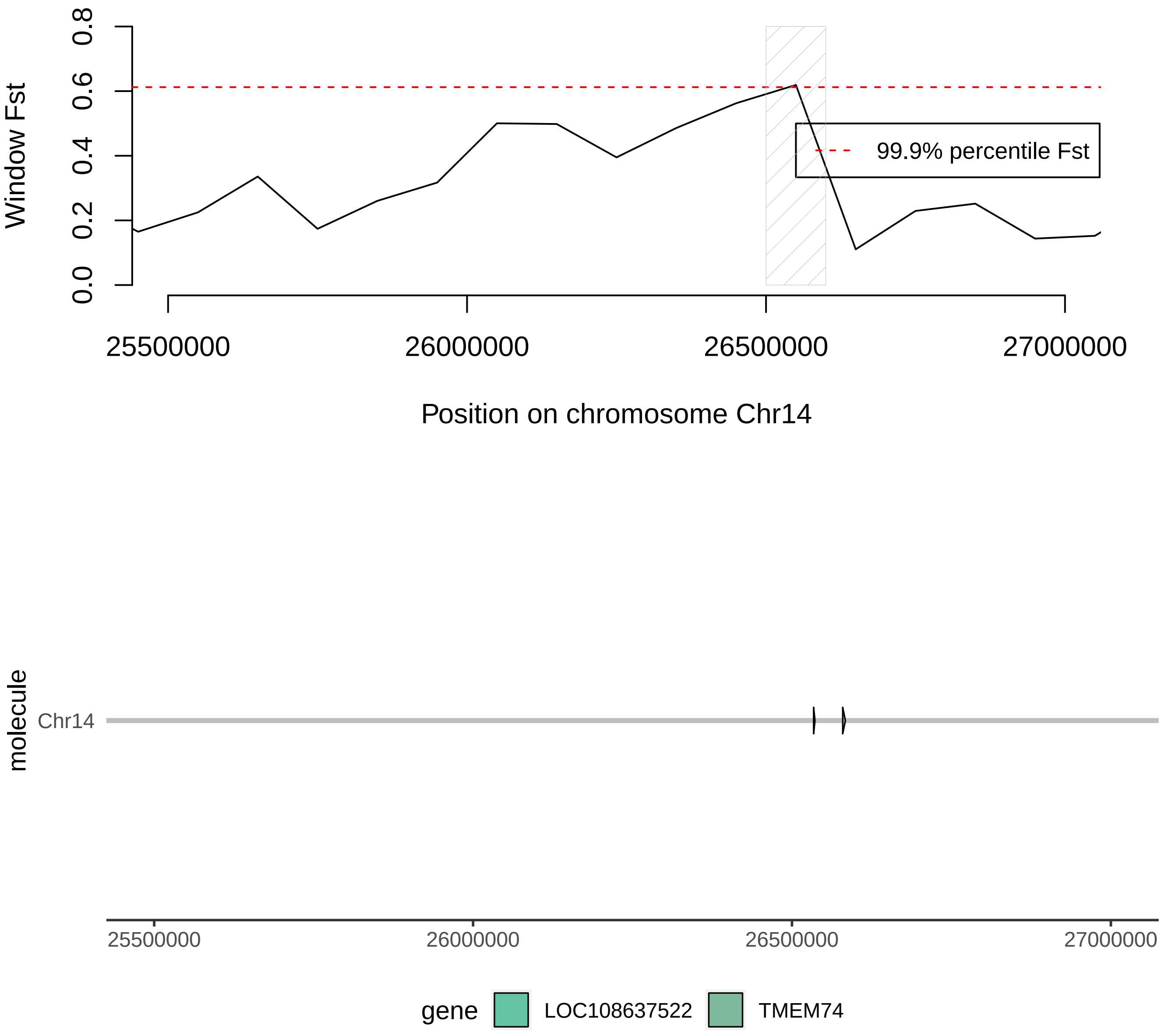

Window Fst
0.0
0.2
0.4
0.6
0.8
99.9% percentile Fst
25500000
26000000
26500000
27000000
Position on chromosome Chr14
molecule
Chr14
25500000
26000000
26500000
27000000
gene
LOC108637522
TMEM74

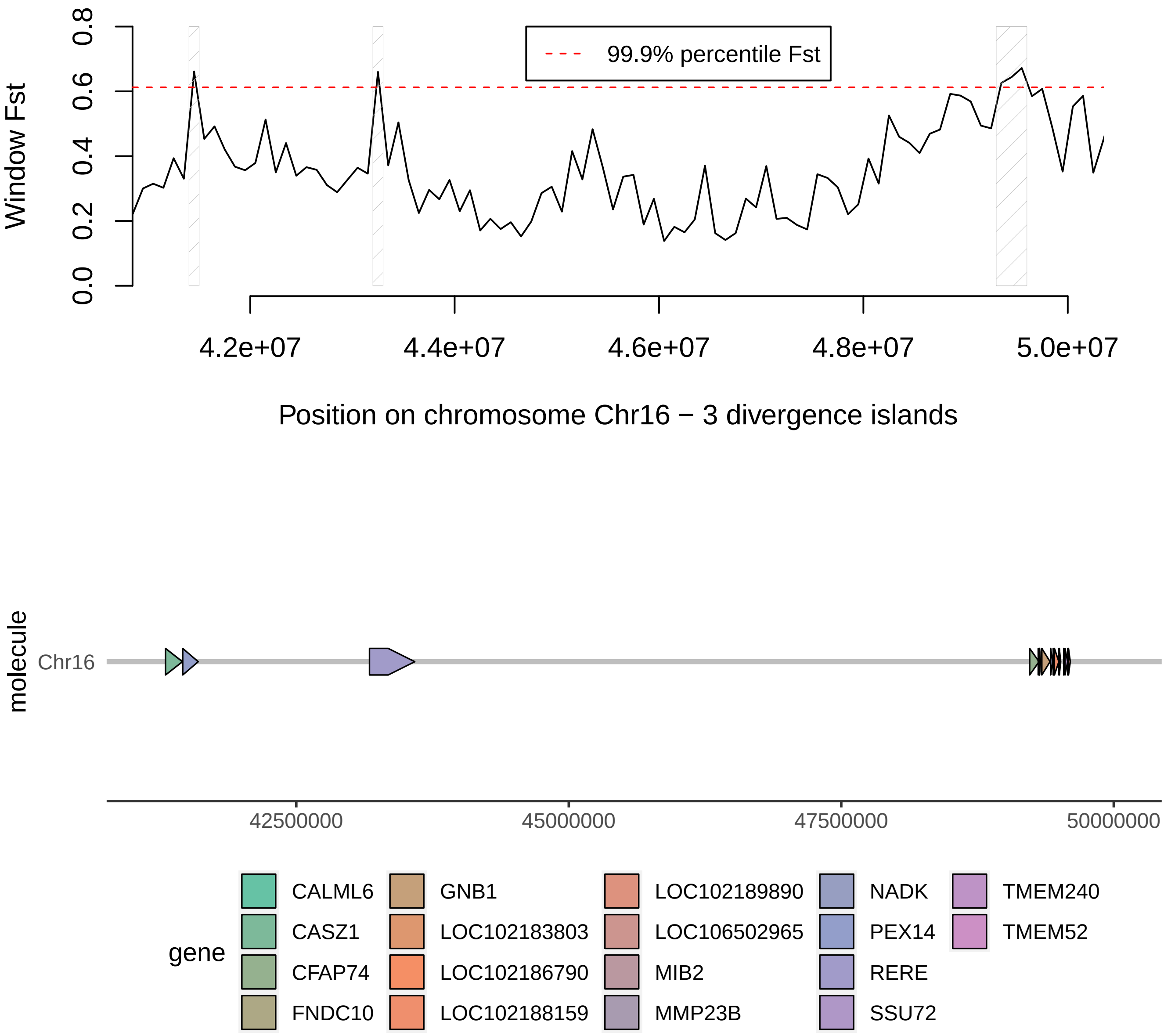
Window Fst
0.0
0.2
0.4
0.6
0.8
99.9% percentile Fst
4.2e+07
4.4e+07
4.6e+07
4.8e+07
5.0e+07
Position on chromosome Chr16 − 3 divergence islands
molecule
Chr16
42500000
45000000
47500000
50000000
gene
CALML6
CASZ1
CFAP74
FNDC10
GNB1
LOC102183803
LOC102186790
LOC102188159
LOC102189890
LOC106502965
MIB2
MMP23B
NADK
PEX14
RERE
SSU72
TMEM240
TMEM52

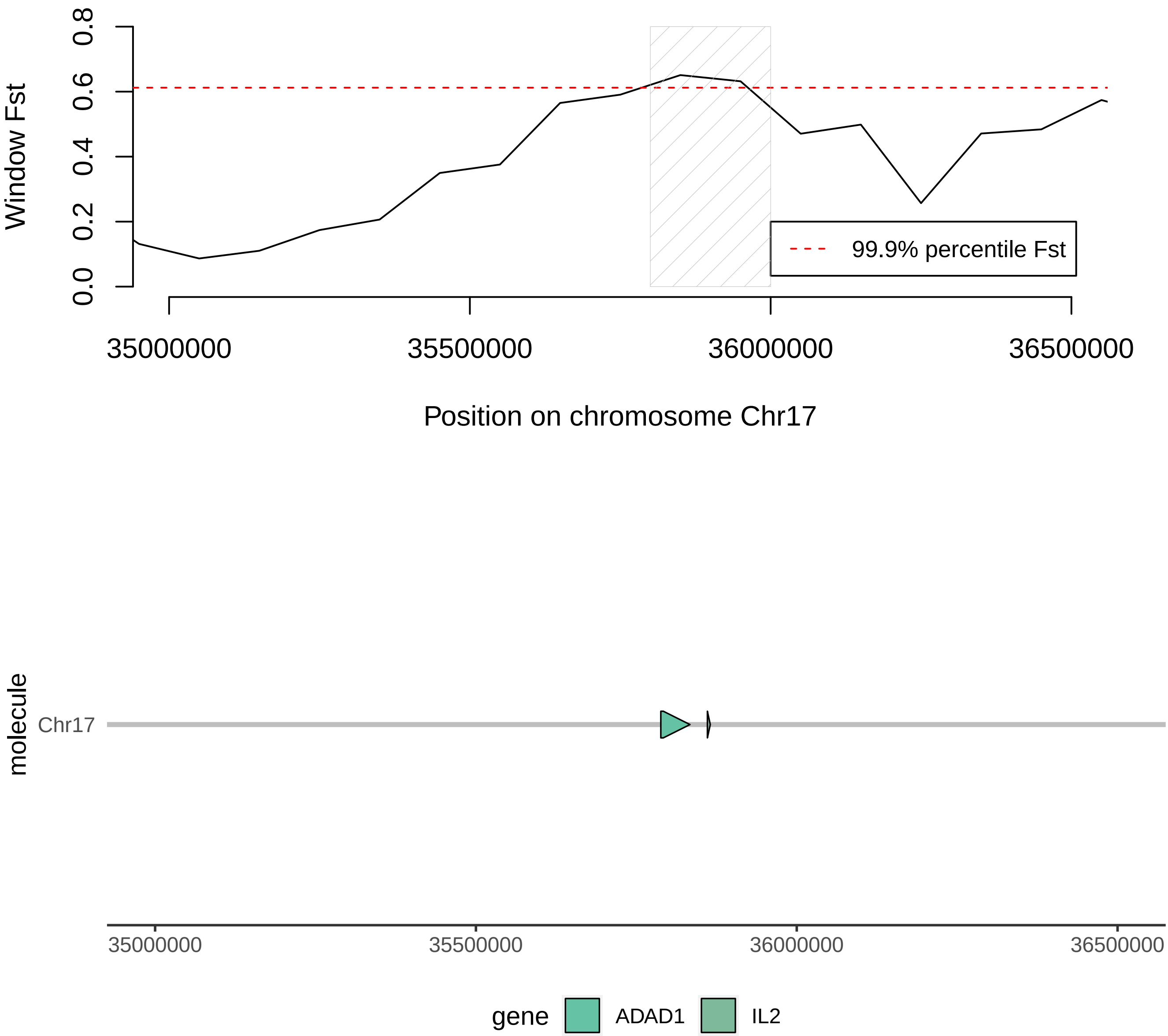
Window Fst
0.0
0.2
0.4
0.6
0.8
99.9% percentile Fst
35000000
35500000
36000000
36500000
Position on chromosome Chr17
molecule
Chr17
35000000
35500000
36000000
36500000
gene
ADAD1
IL2

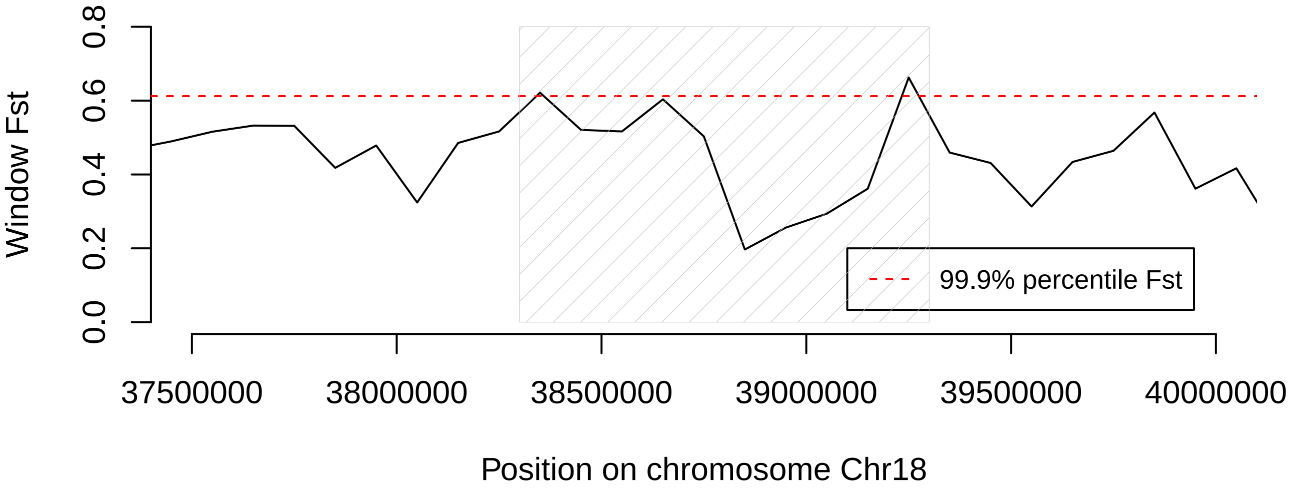

Window Fst
0.0
0.2
0.4
0.6
0.8
99.9% percentile Fst
37500000
38000000
38500000
39000000
39500000
40000000
Position on chromosome Chr18


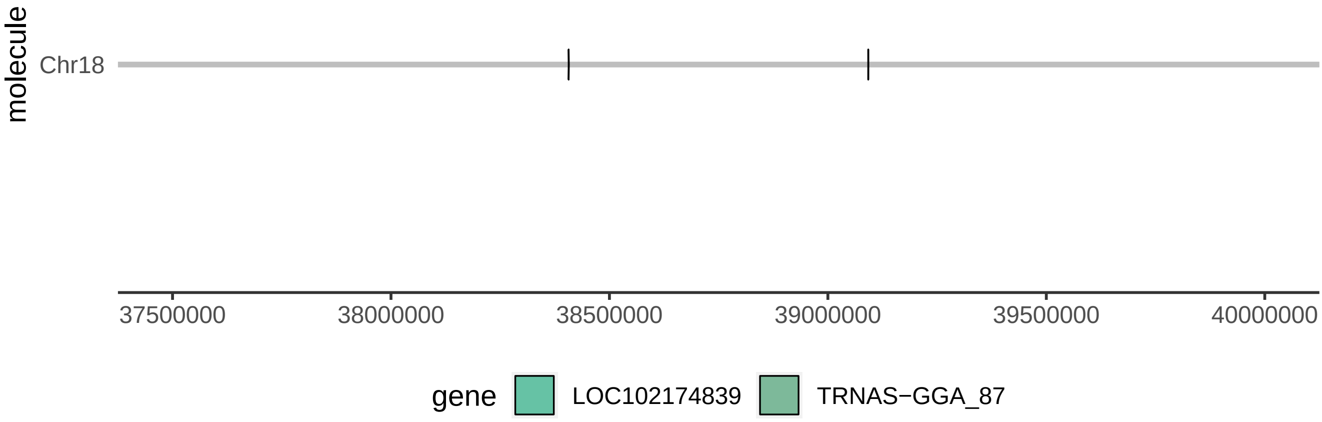

molecule
Chr18
37500000
38000000
38500000
39000000
39500000
40000000
gene
LOC102174839
TRNAS−GGA_87

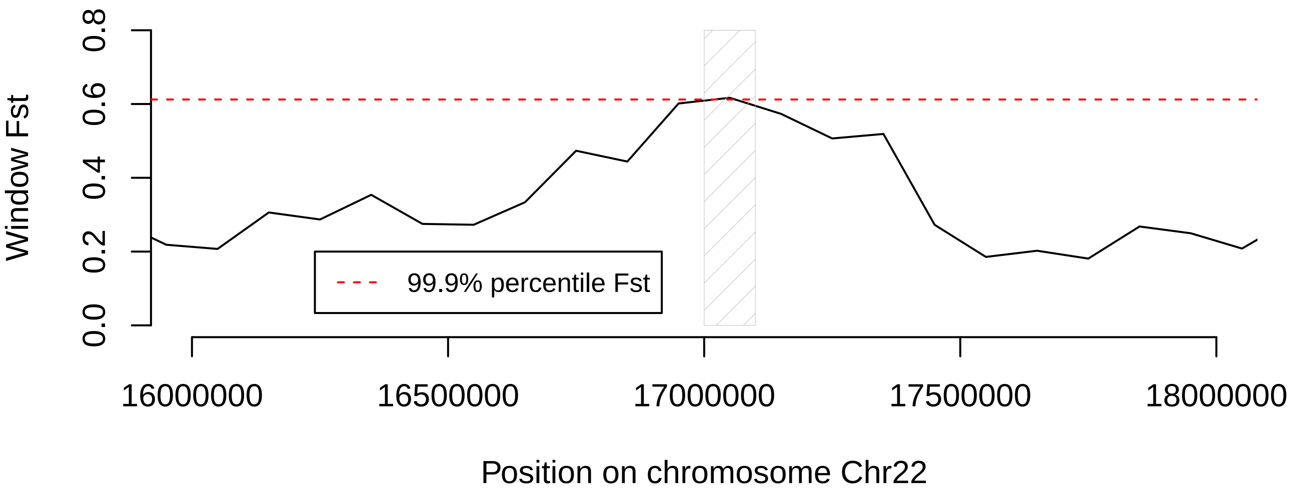

Window Fst
0.0
0.2
0.4
0.6
0.8
99.9% percentile Fst
16000000
16500000
17000000
17500000
18000000
Position on chromosome Chr22


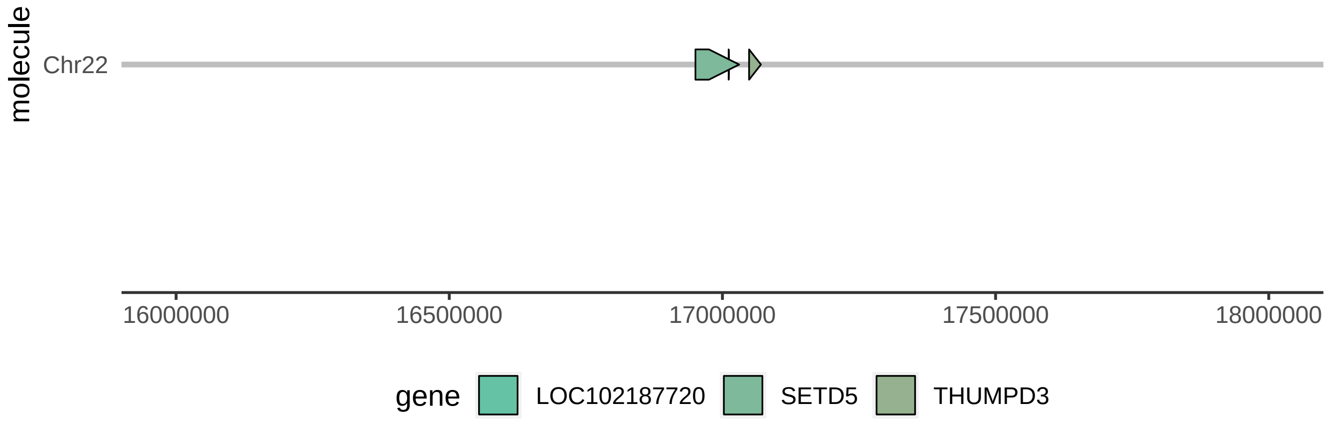

molecule
Chr22
16000000
16500000
17000000
17500000
18000000
gene
LOC102187720
SETD5
THUMPD3

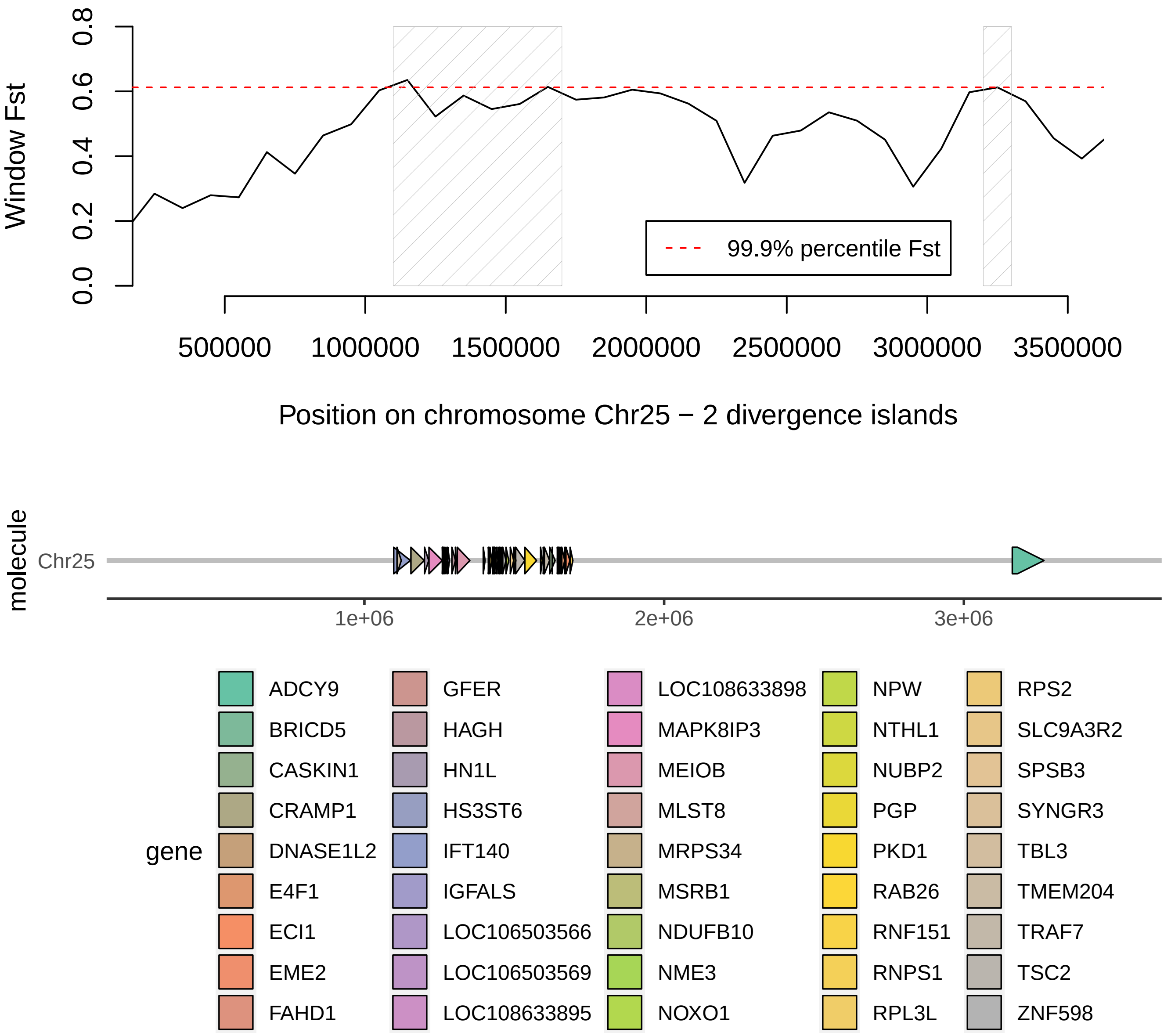

Window Fst
0.0
0.2
0.4
0.6
0.8
99.9% percentile Fst
500000
1000000
1500000
2000000
2500000
3000000
3500000
Position on chromosome Chr25 − 2 divergence islands
molecule
Chr25
1e+06
2e+06
3e+06
gene
ADCY9
BRICD5
CASKIN1
CRAMP1
DNASE1L2
E4F1
ECI1
EME2
FAHD1
GFER
HAGH
HN1L
HS3ST6
IFT140
IGFALS
LOC106503566
LOC106503569
LOC108633895
LOC108633898
MAPK8IP3
MEIOB
MLST8
MRPS34
MSRB1
NDUFB10
NME3
NOXO1
NPW
NTHL1
NUBP2
PGP
PKD1
RAB26
RNF151
RNPS1
RPL3L
RPS2
SLC9A3R2
SPSB3
SYNGR3
TBL3
TMEM204
TRAF7
TSC2
ZNF598
