## Supplementary figures and images for "Persistent gene flow suggests an absence of reproductive isolation in an African antelope speciation model"

### SupplementaryFigure15

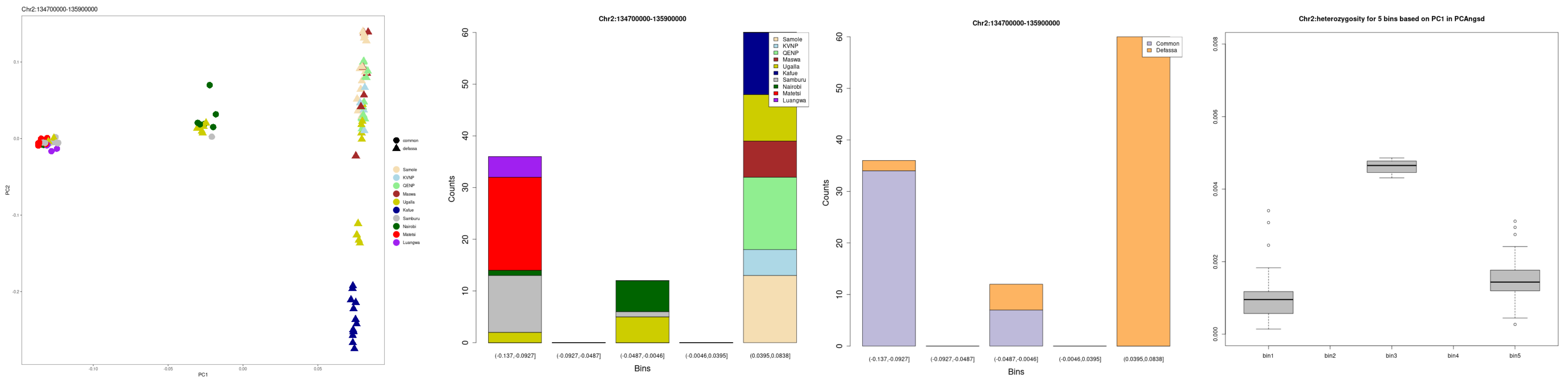

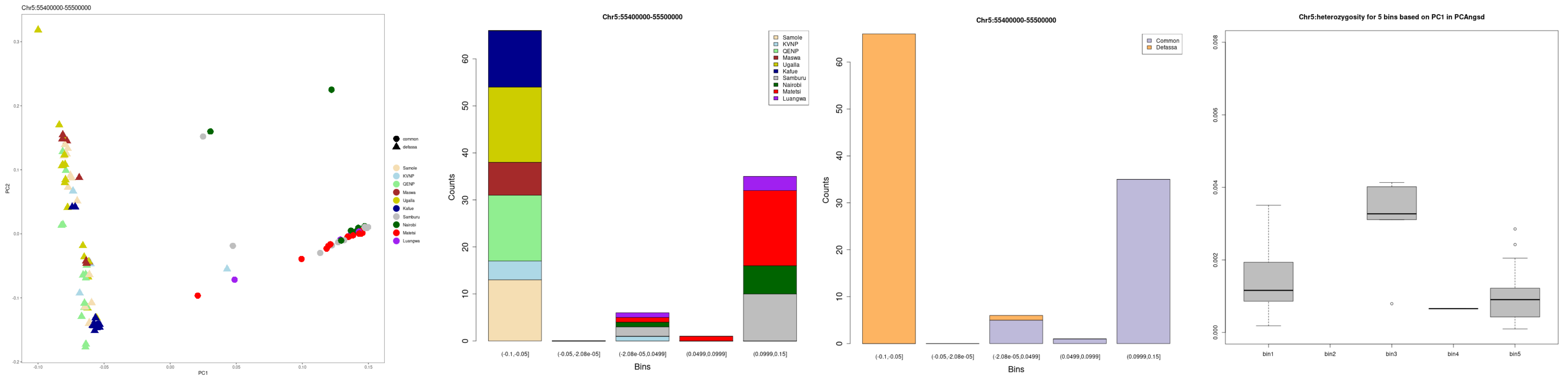

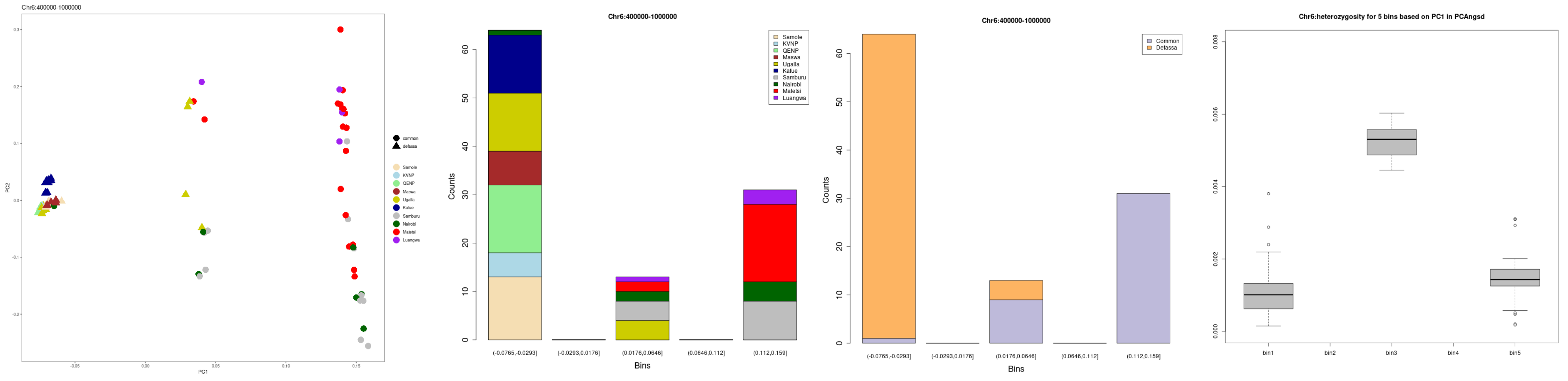

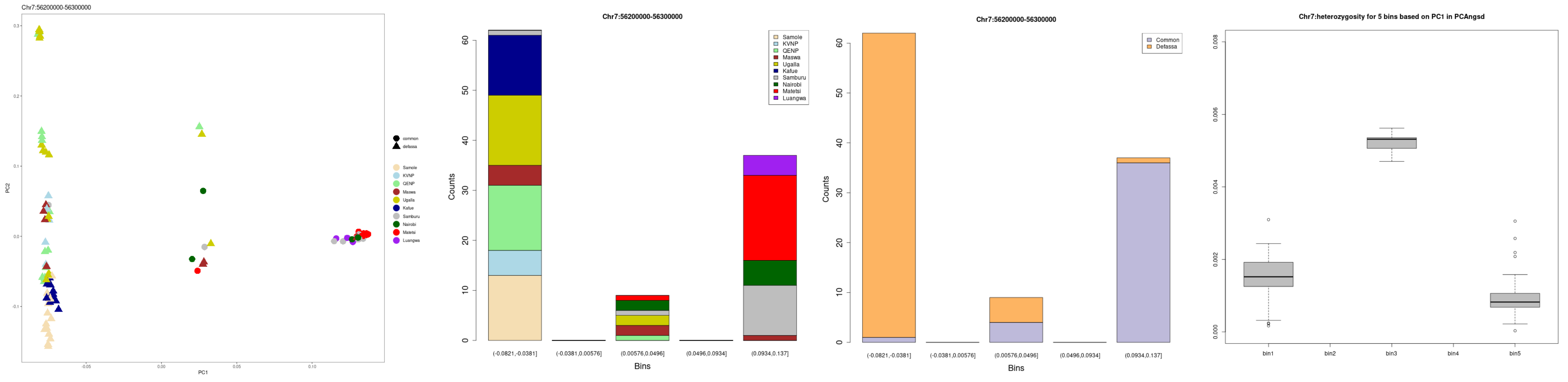

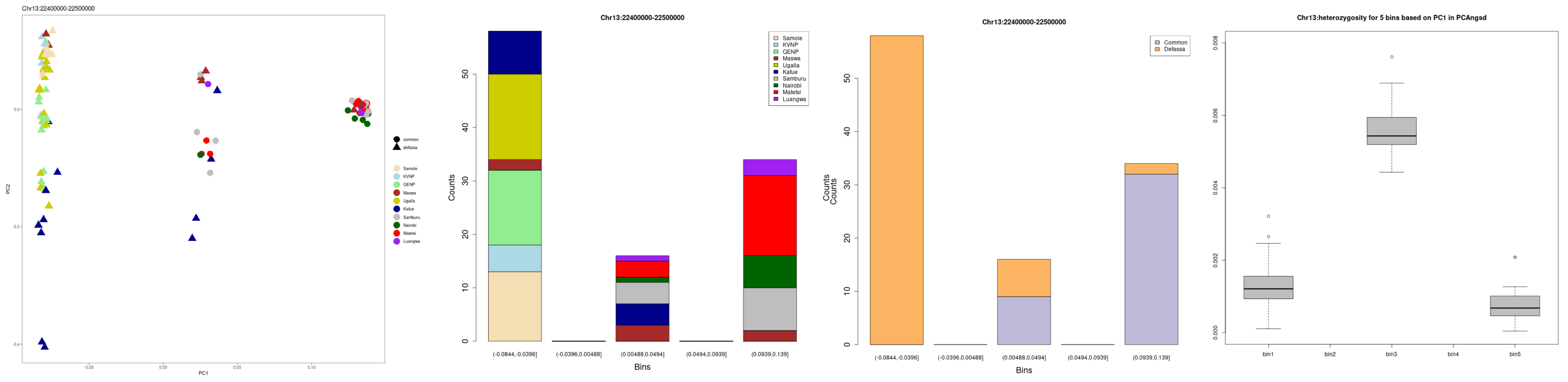

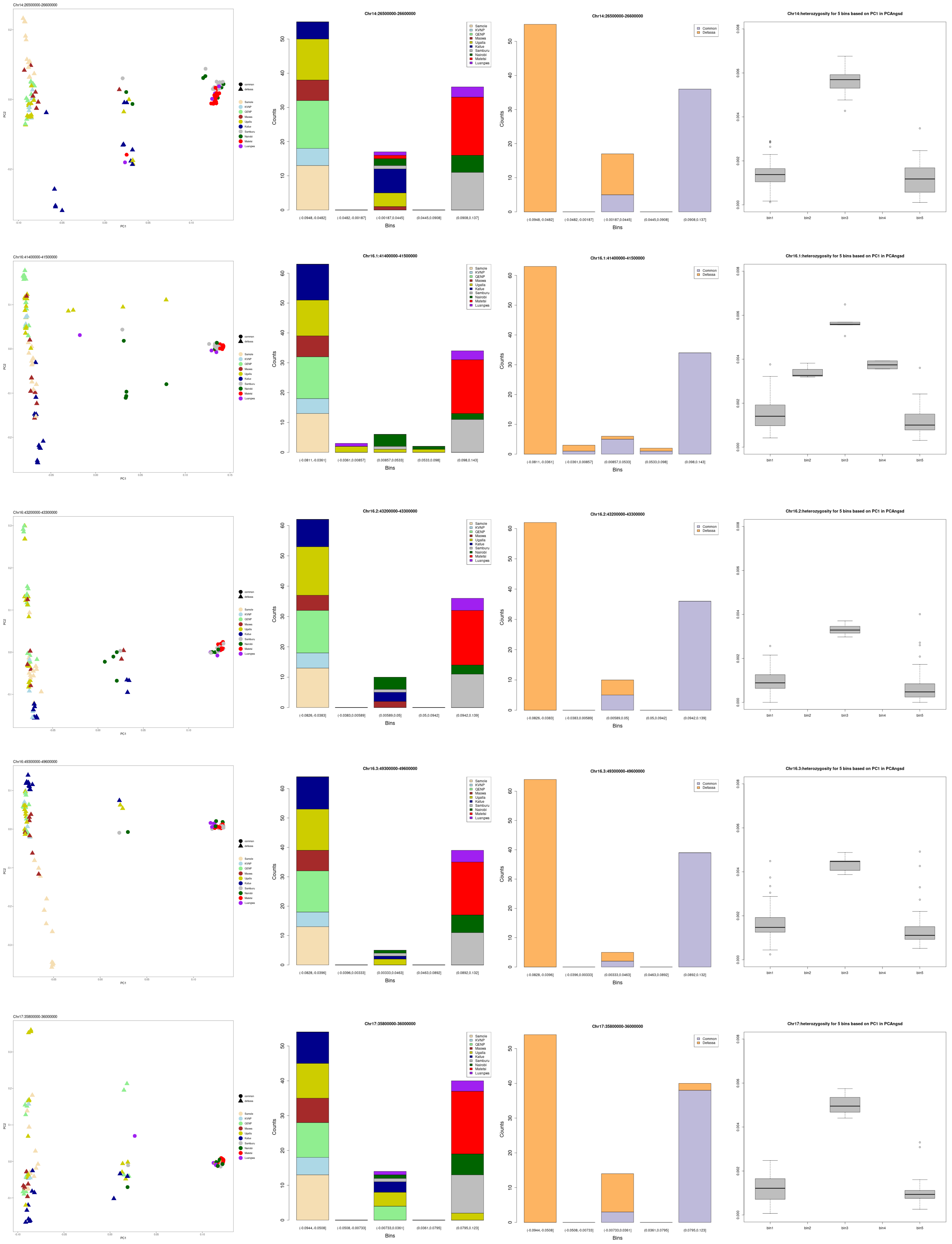

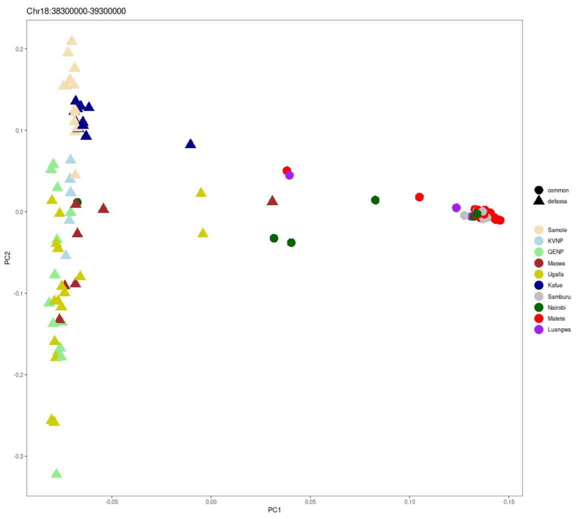

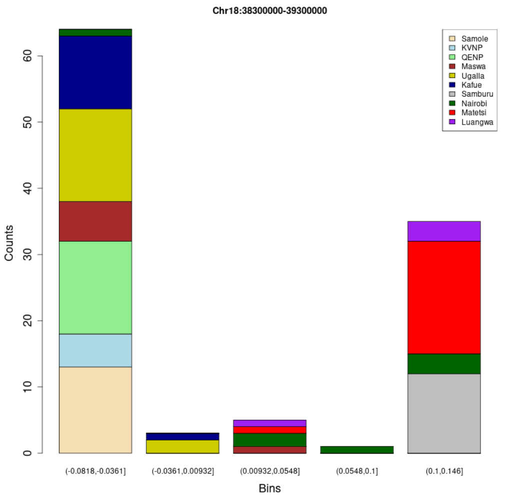

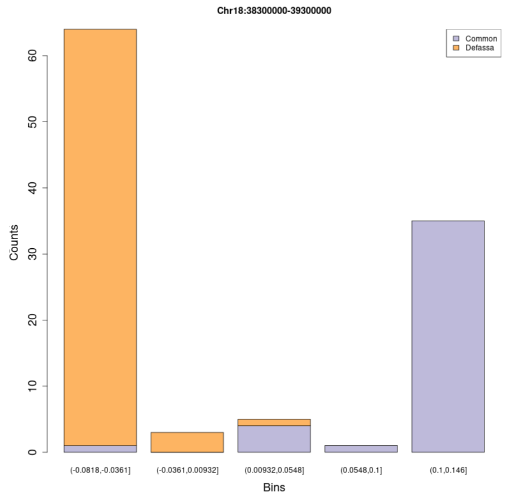

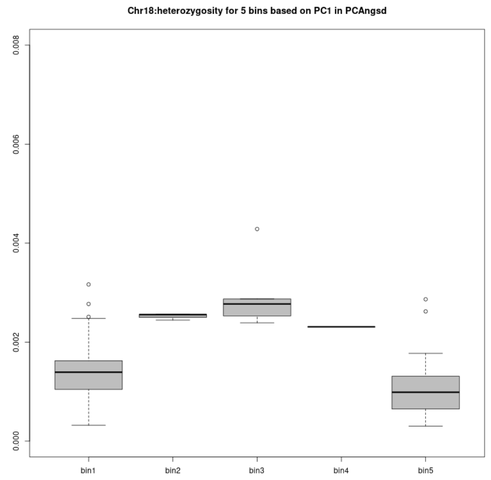

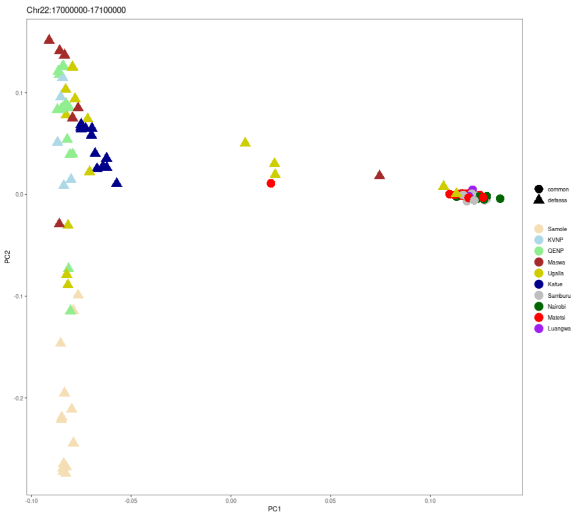

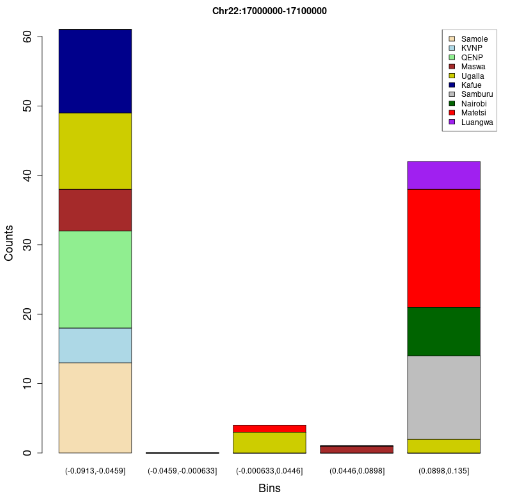

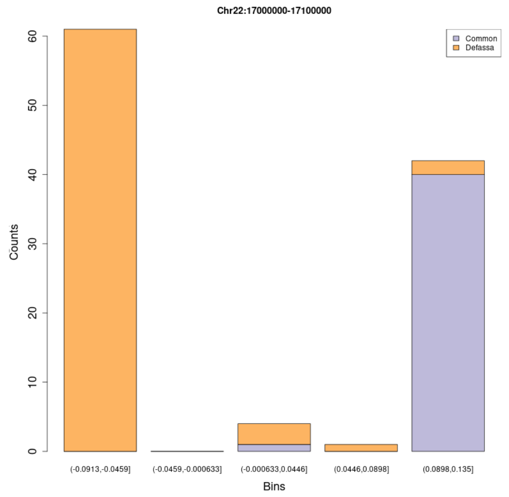

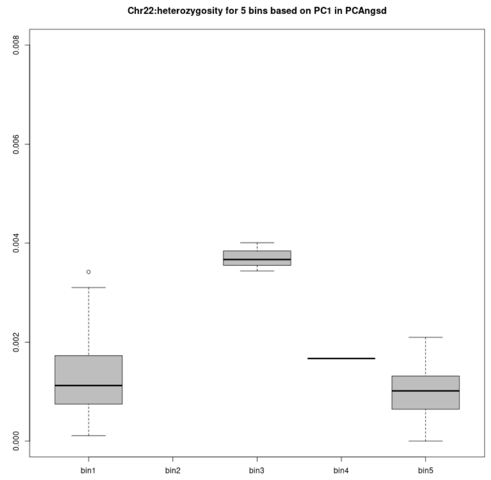

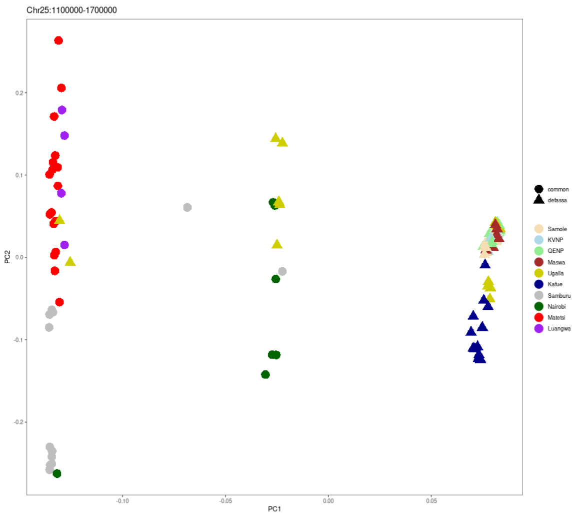

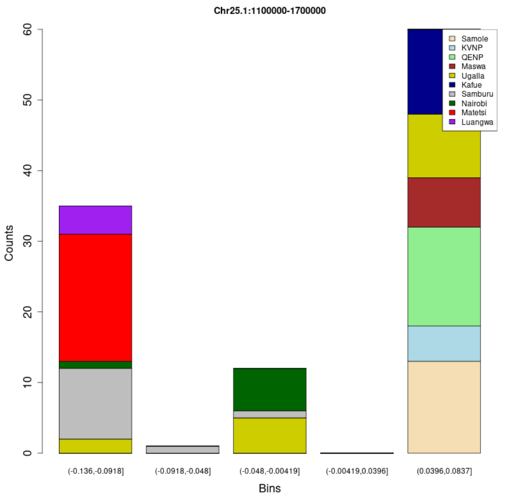

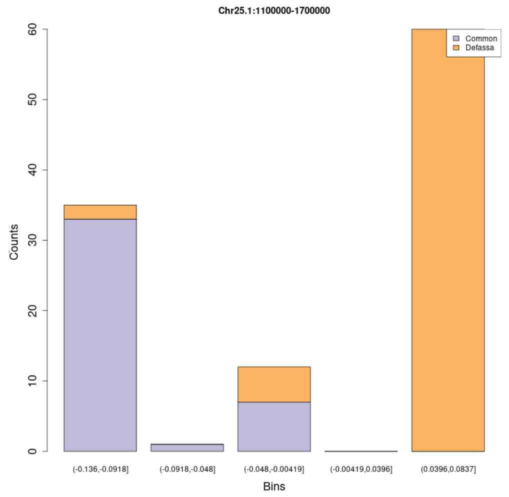

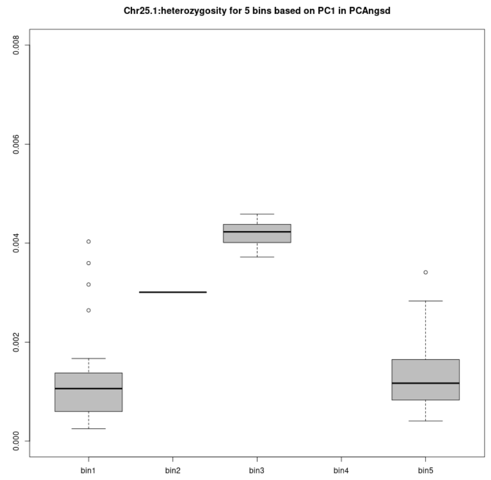

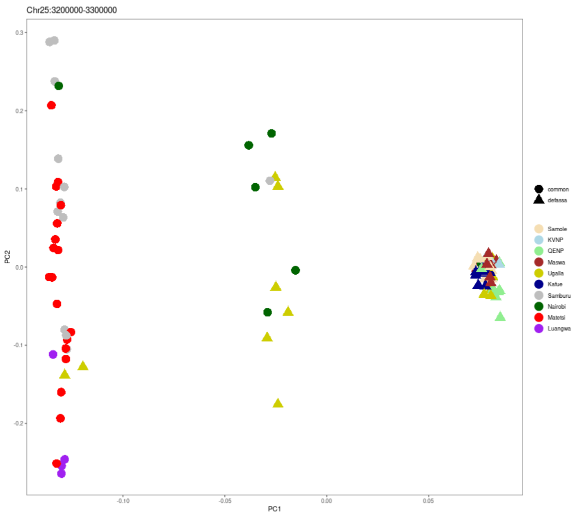

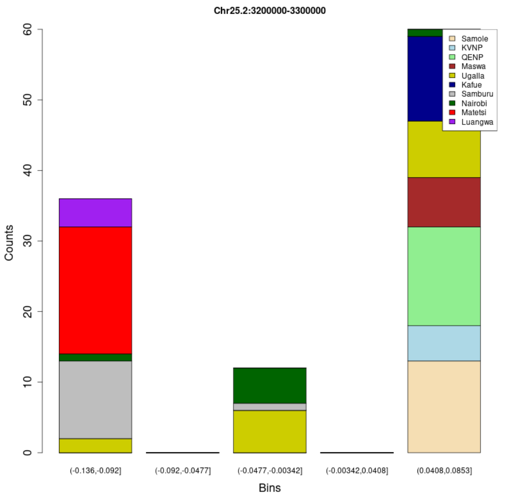

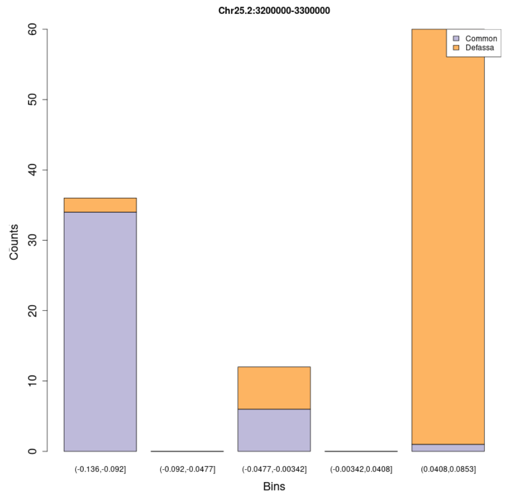

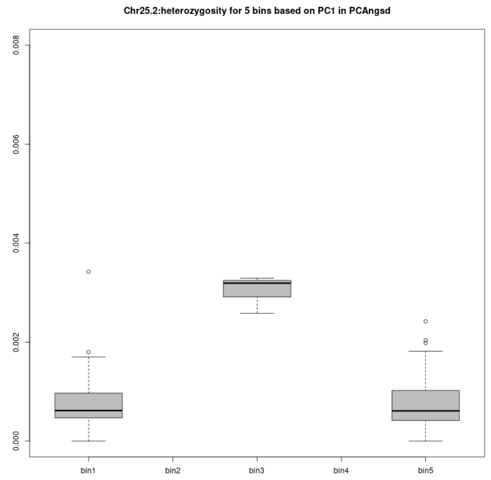

### SupplementaryFigure16

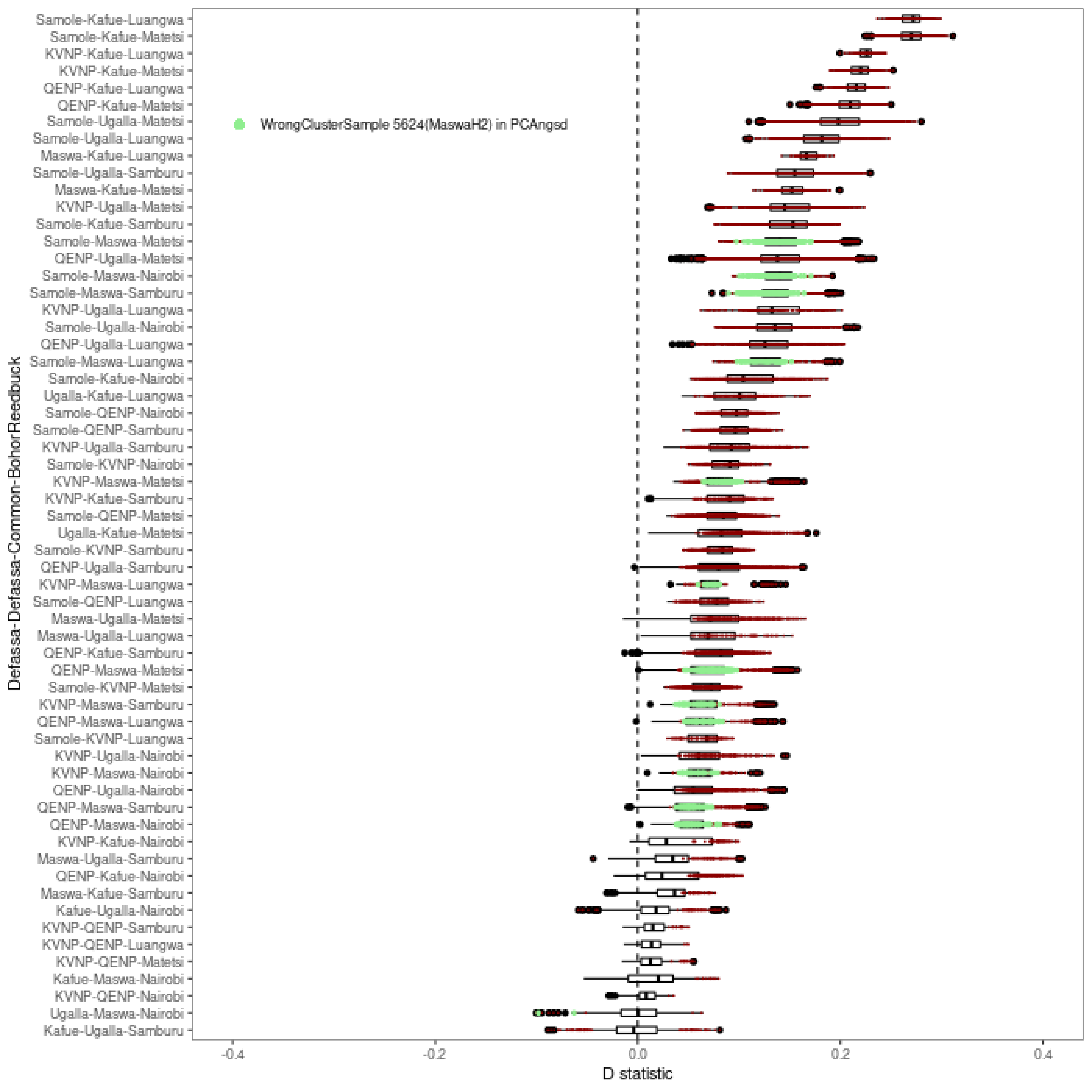

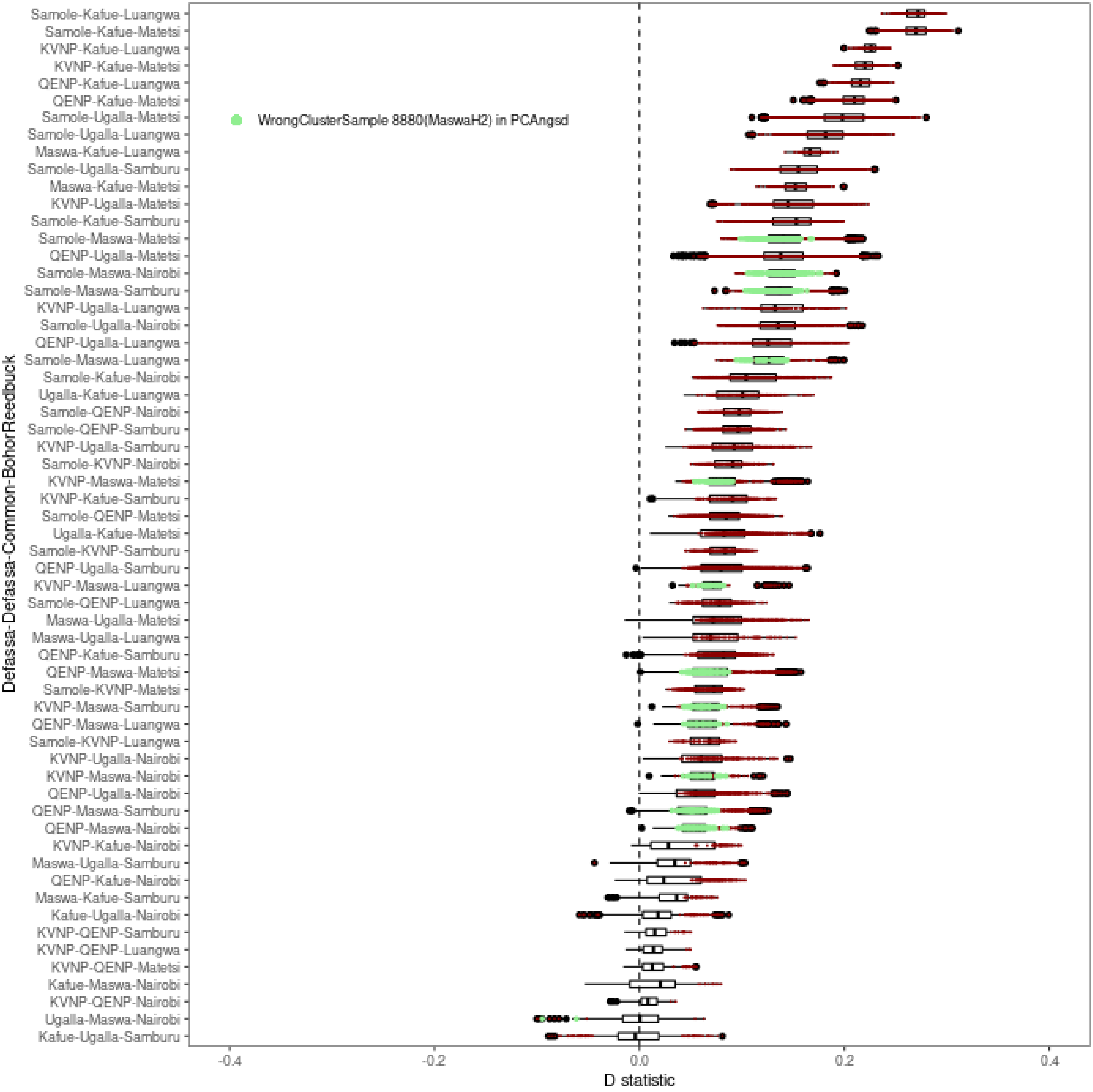

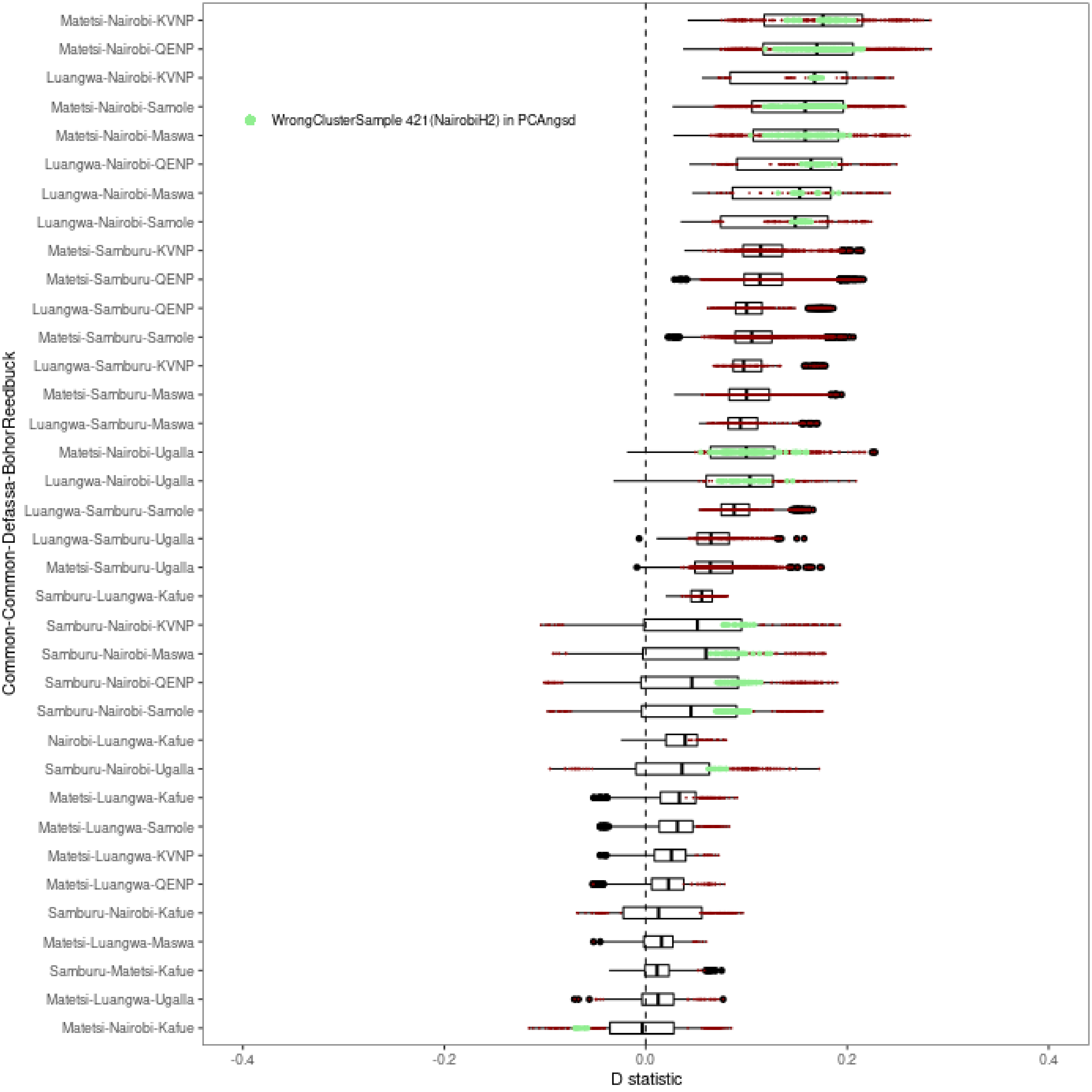

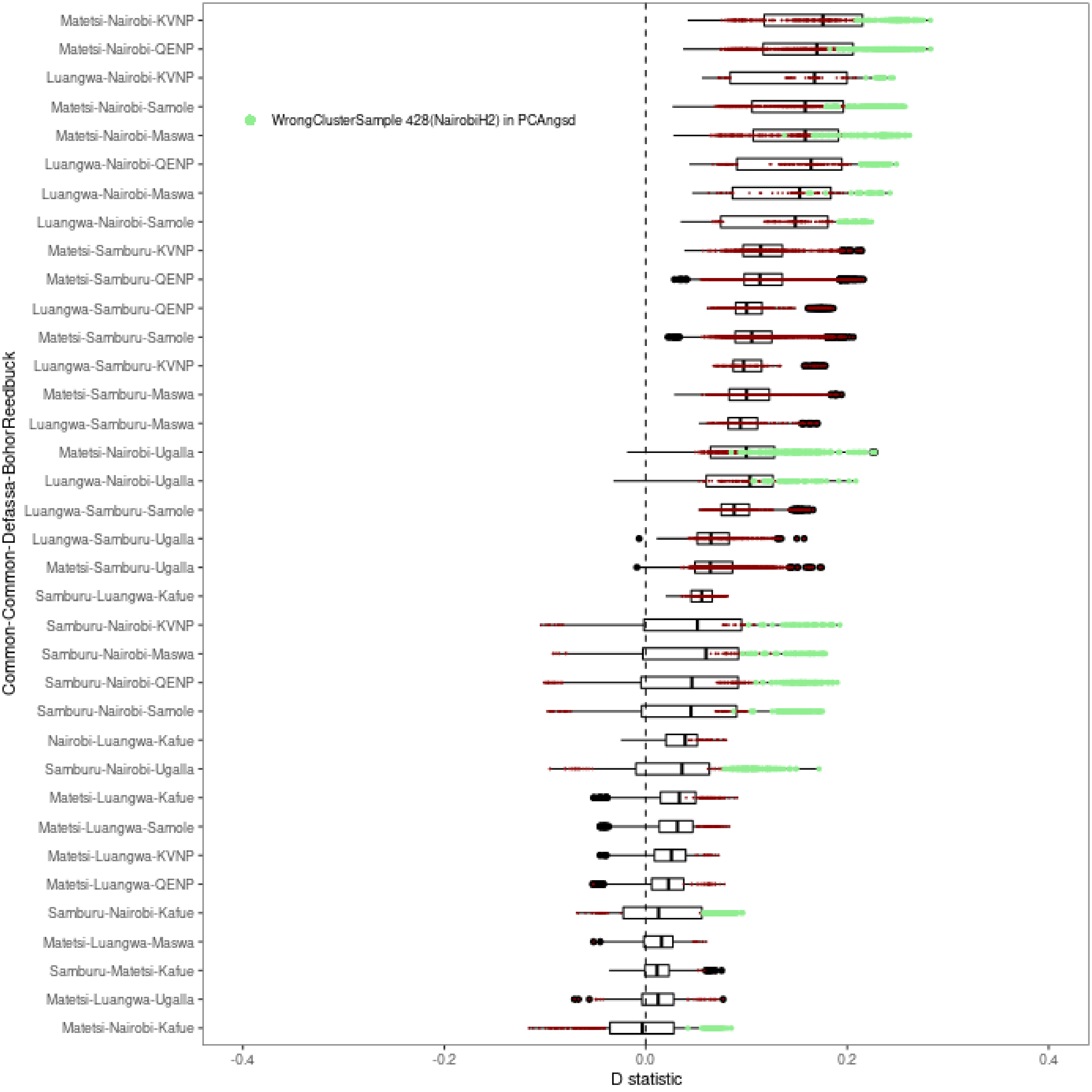

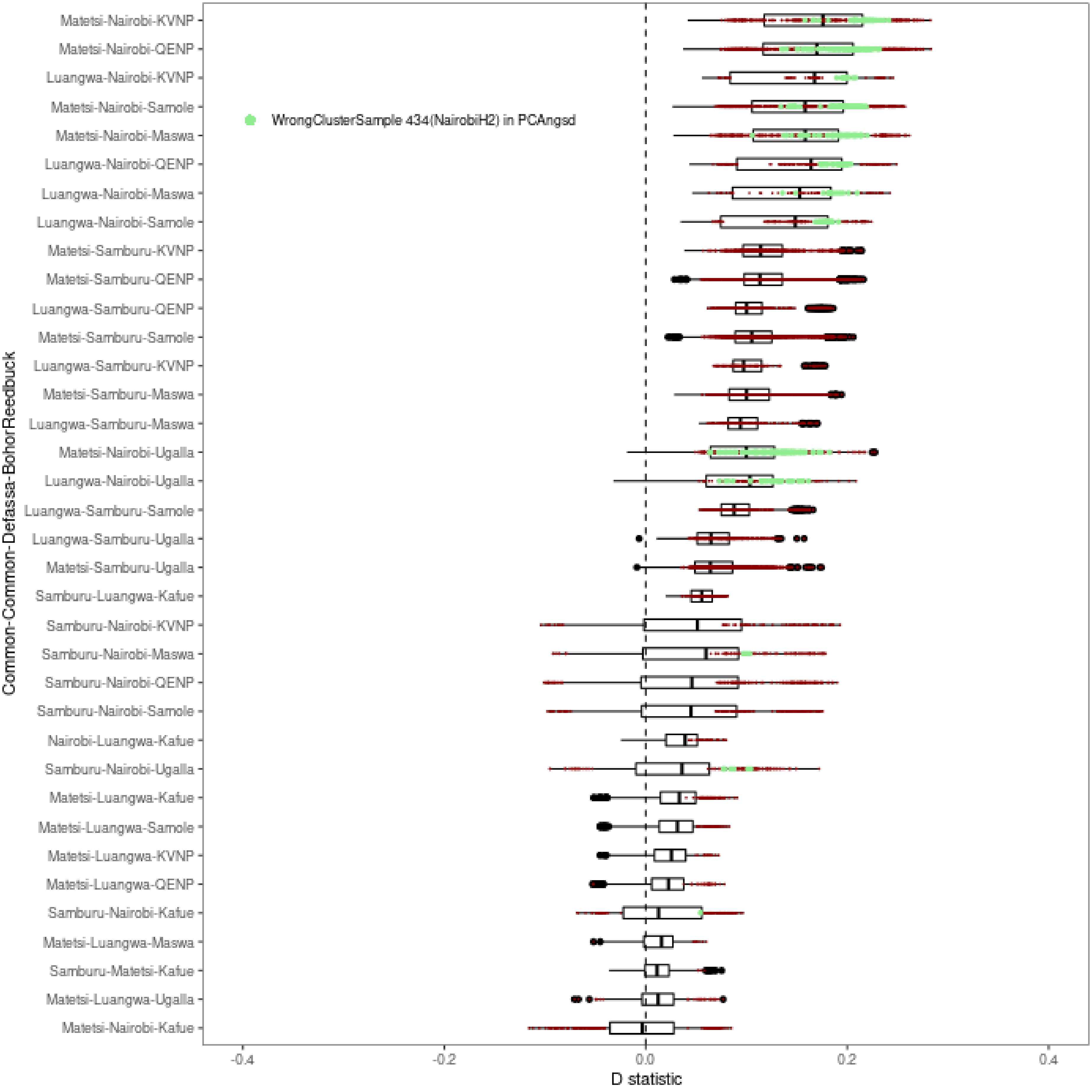

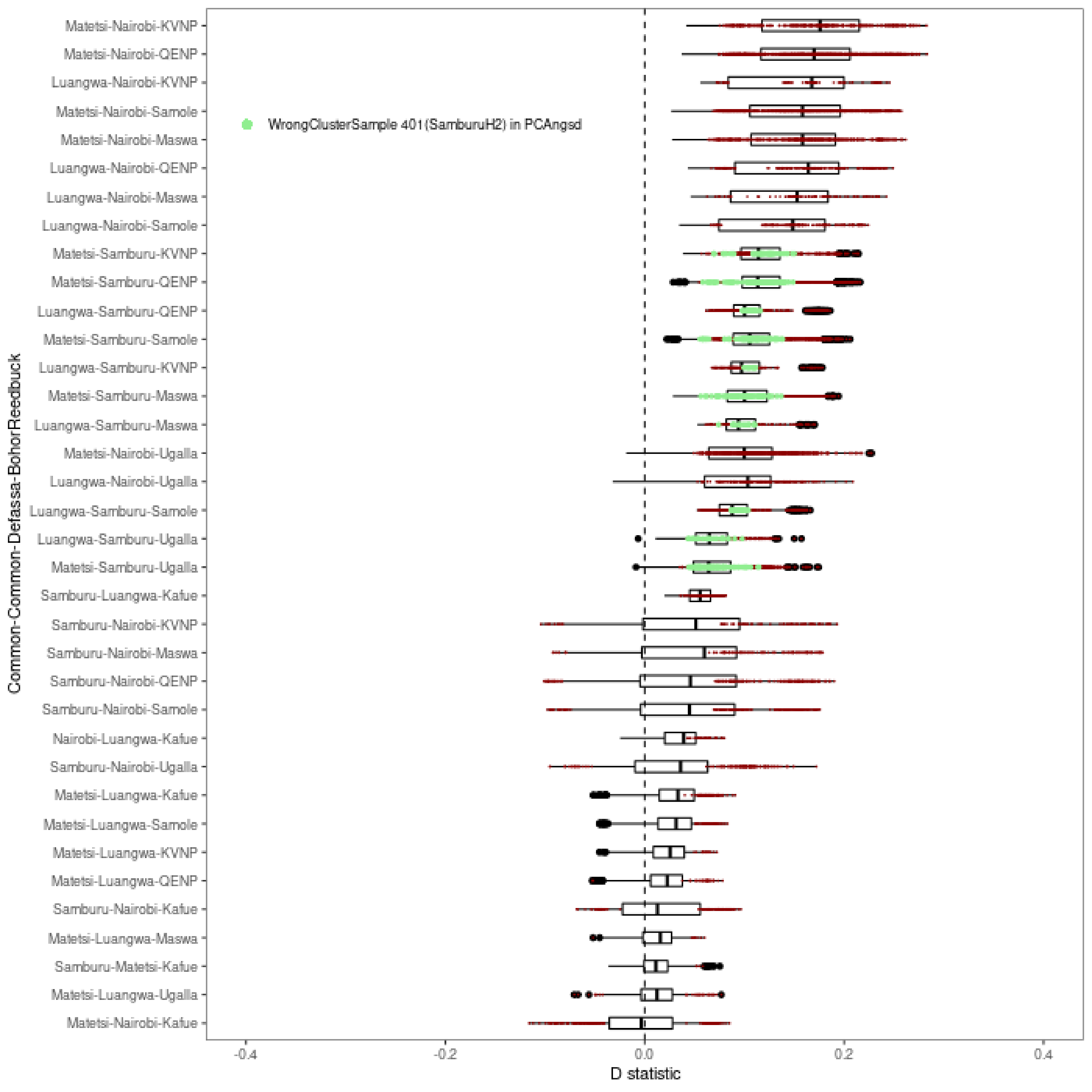

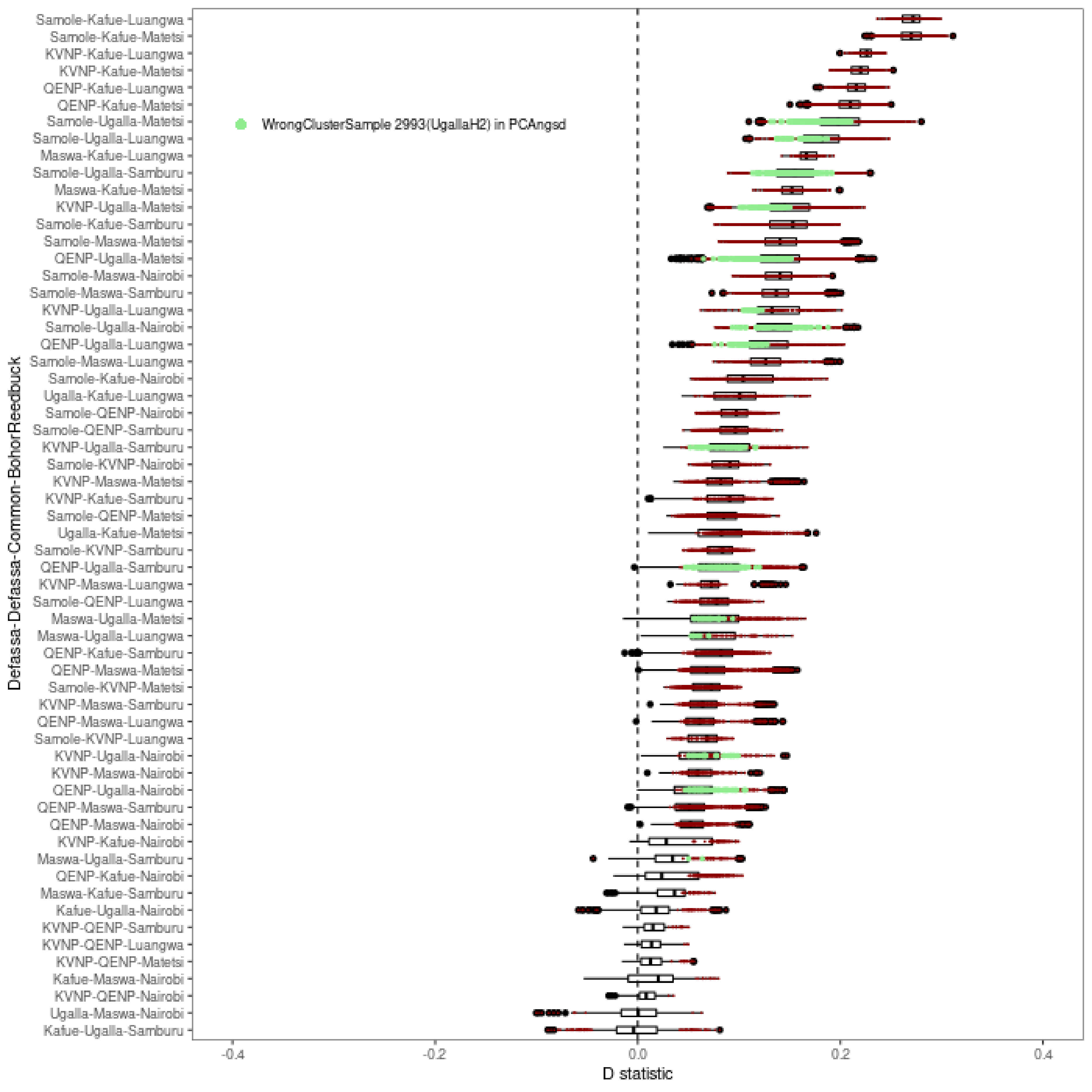

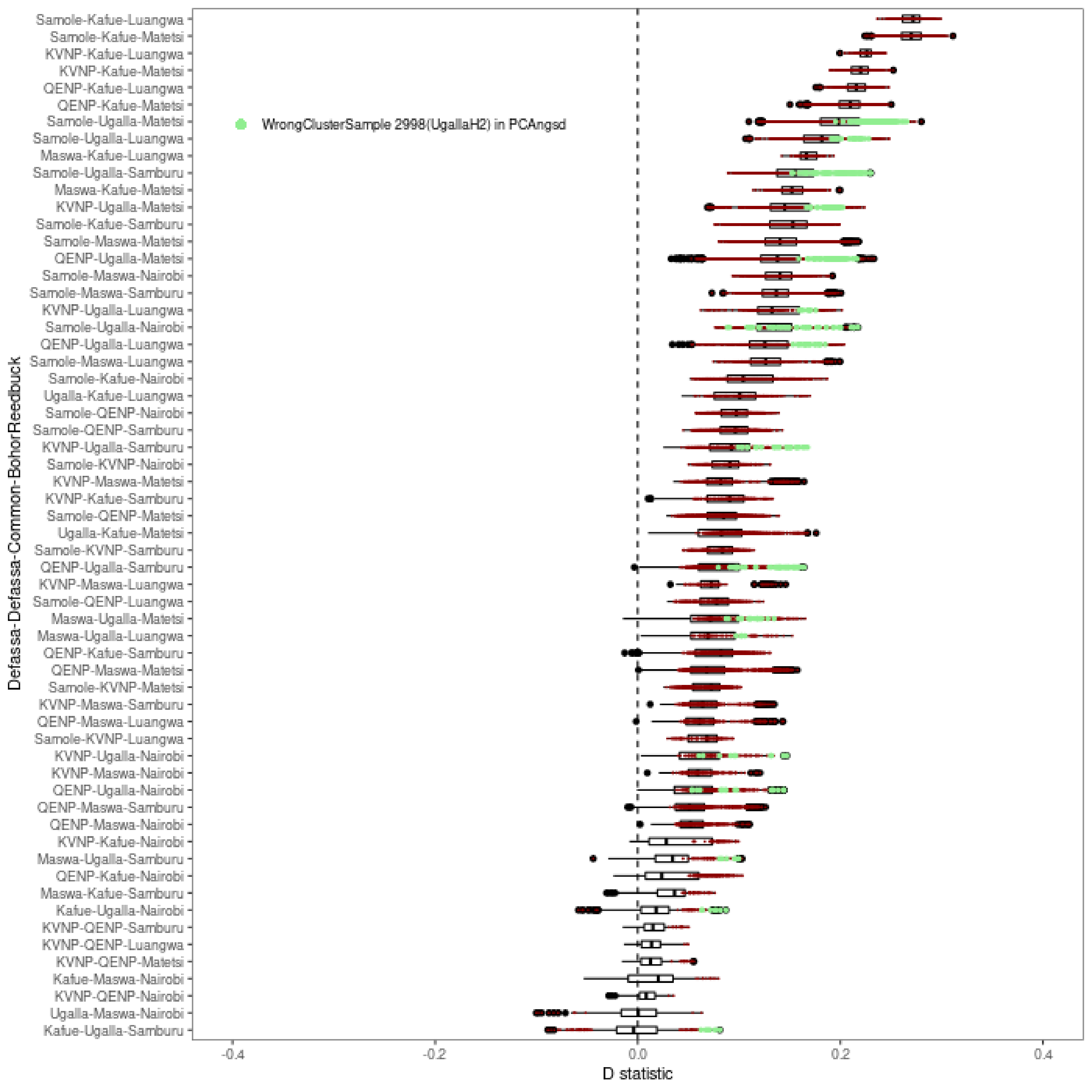

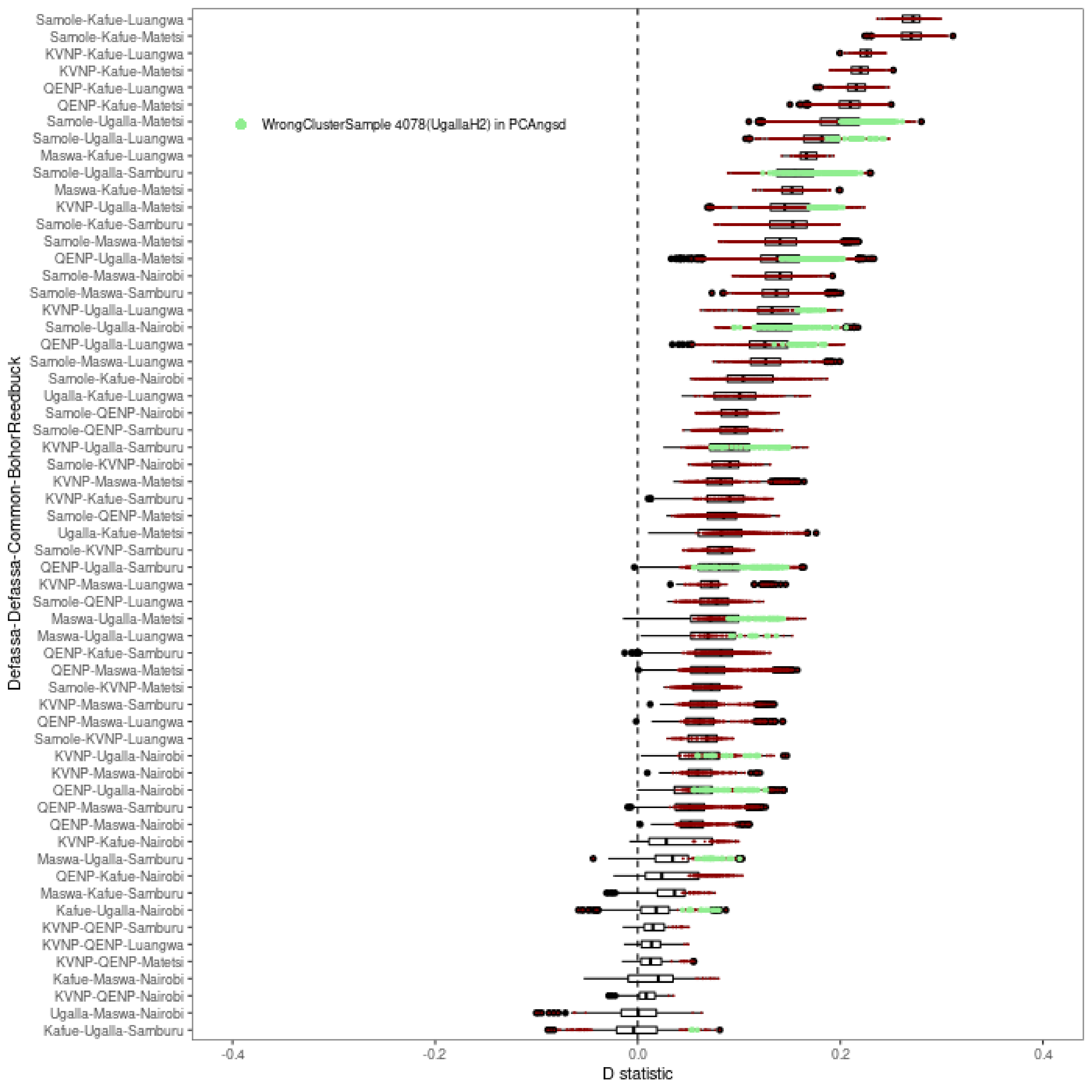

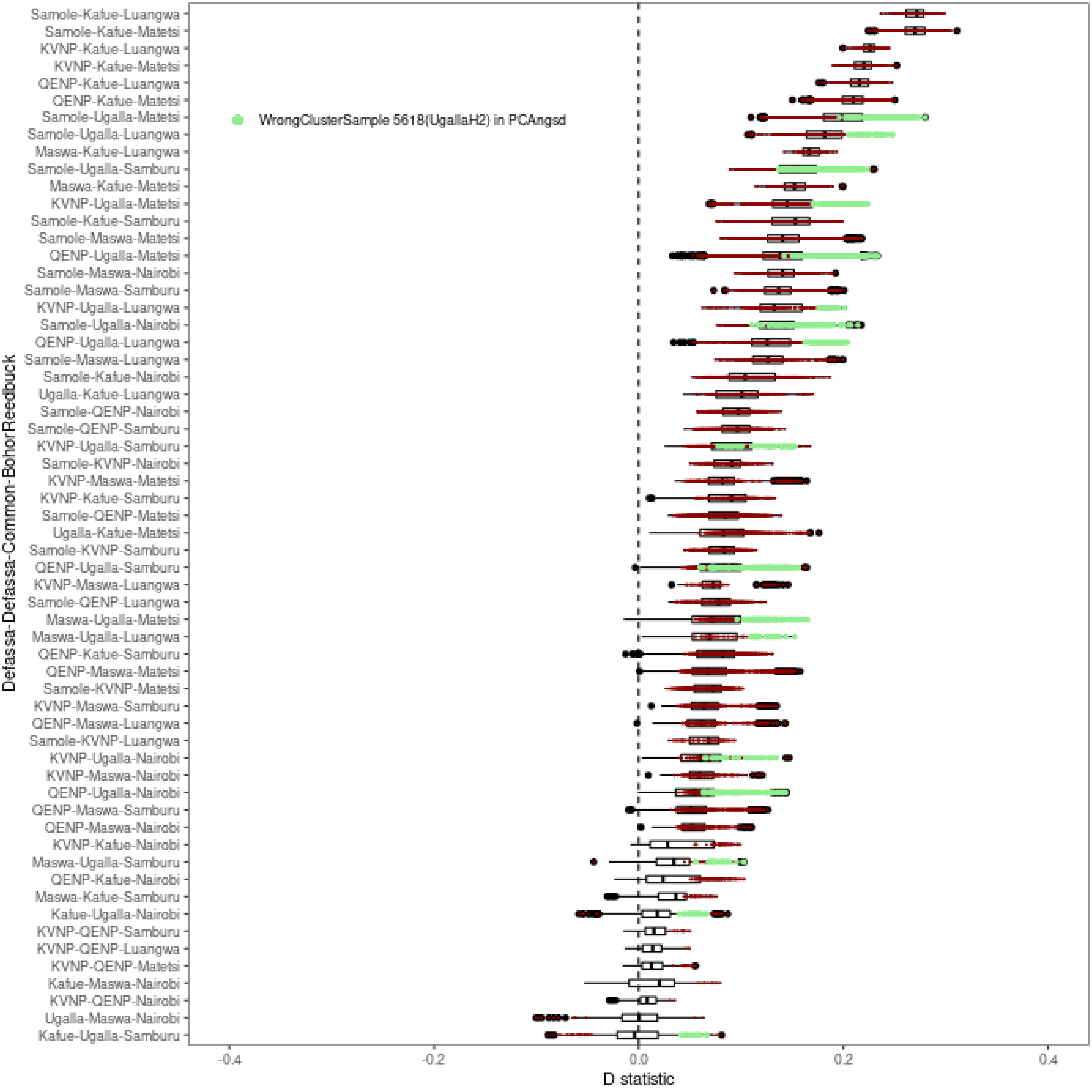
